# Human cerebellum and ventral tegmental area interact during extinction of learned fear

**DOI:** 10.1101/2024.11.06.622063

**Authors:** Enzo Nio, Patrick Pais Pereira, Nicolas Diekmann, Mykola Petrenko, Alice Doubliez, Thomas M. Ernst, Giorgi Batsikadze, Stefan Maderwald, Cornelius Deuschl, Metin Üngör, Sen Cheng, Christian J. Merz, Harald H. Quick, Dagmar Timmann

## Abstract

The key elements for fear extinction learning are unexpected omissions of expected aversive events, which are considered to be rewarding. Given its reception of reward information, we tested the hypothesis that the cerebellum contributes to reward-like prediction error processing driving extinction learning via its connections with the ventral tegmental area (VTA). Forty-three young and healthy participants performed a three-day fear conditioning paradigm in a 7T MR scanner. The cerebellum and VTA were active during unexpected omissions of aversive unconditioned stimuli in the initial extinction trials and in other learning phases, in line with the proposed role of prediction-error processing. Increased functional connectivity was observed between the cerebellum and VTA, indicating that they are functionally coupled during fear extinction learning. These results suggest that an interaction between the cerebellum and VTA should be incorporated into the existing model of the fear extinction network.

## Introduction

Deficits in learning to extinguish previously associated threat responses to cues that no longer signal danger are thought to be one of the main causes in the development of anxiety disorders, including post-traumatic stress disorder and social anxiety disorder^1^. Exposure therapy addresses this deficit and attempts to compensate for failed or absent extinction learning from the past^2^. This is modeled effectively with extinction training in classical fear conditioning paradigms where the previously paired conditioned stimulus (CS+) is no longer followed by the aversive unconditioned stimulus (US)^3,4^. At the beginning of extinction training the omission of the US is unexpected and this unexpected lack of the US is considered to be rewarding^5–7^. This reward-like prediction error is thought to drive safety learning. This newly learned safety association should then inhibit the initially learned fear association (which is never fully lost)^8^.

The reward system in the brain is primarily associated with the mesolimbic dopamine system^9^, i.e., the ventral tegmental area (VTA) and the ventral striatum. In fact, recent rodent studies provide strong evidence that the VTA is central for extinction learning, and that (reward-like) prediction errors are encoded in a subset of VTA neurons^5–7,10^.

Importantly, new evidence indicates that the cerebellum receives reward signals^11–13^, and has direct efferent connections with the mesolimbic dopaminergic system, in particular the VTA^14–17^.

Functional magnetic resonance imaging (fMRI) findings in healthy human participants suggest that the cerebellum is involved in processing of prediction errors in fear conditioning^18,19^. Ernst et al.^18^ studied acquisition of learned fear responses using a partial reinforcement rate, i.e., a subset of CS+ trials were not reinforced in acquisition training. Cerebellar activations were strongest in the unreinforced CS+ trials, when the US was expected but did not occur. An unreinforced CS+ trial can be considered as a very first extinction trial, and cerebellar activations may reflect prediction errors driving extinction learning. Likewise, recordings from the VTA in rodents showed strong activations at the time the US is expected and does not occur during initial extinction trials^6,10^. The aim of the present fMRI study is therefore to test the hypothesis that the cerebellum contributes to fear extinction learning via its connection to the VTA.

In contrast to the previous one-day study design^18^, here we used a three-day design with acquisition training on day 1, extinction training on day 2 and a recall test on day 3 (Figure 1A). This multi-day study design allowed enough time for consolidation of both fear and extinction learning. Cerebellar activity related to the unexpected omission of the US was assessed in four different learning phases, that is, during extinction training (in initial CS+ trials), during the recall test (in initial CS+ trials), during reacquisition (in interspersed non-reinforced CS+ trials), and during reextinction (the initial CS+ trials immediately following reinforced trials) (Figure 1A). The CSs were geometric shapes (i.e. square and diamond), and the US was an electrical stimulation (Figure 1B). Full (100%) reinforcement was used during acquisition on day 1 to maximize the size of the reward-like prediction error elicited by the initial unexpected US omissions during extinction. Learning phases differed in the underlying learning state, ranging from newly acquired fear, to spontaneous recovery after consolidated extinction^20^, to reactivated fear and reextinction, providing four distinct contexts in which to examine cerebellar and VTA responses to unexpected US omission^21^.

**Figure 1A:**
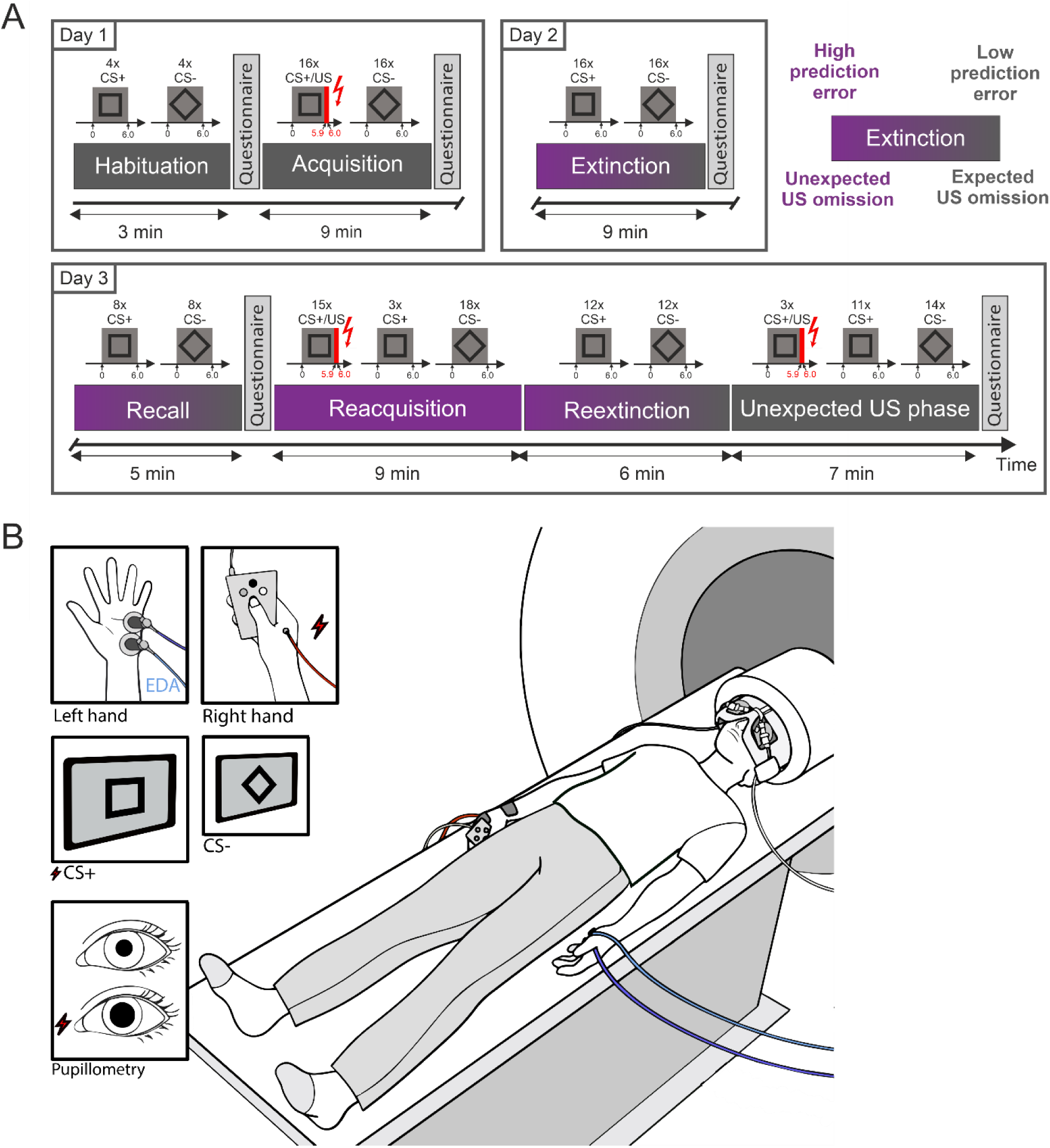
Overview of the three-day fear conditioning paradigm. Day 1 consisted of habituation and acquisition training, day 2 of extinction training, and day 3 of a recall test followed by reacquisition, reextinction, and the unexpected US phase. Extinction, recall, reacquisition, and reextinction are highlighted in purple, indicating phases in which unexpected US omissions occurred. Extinction, recall, and reextinction are shown with a gradient to illustrate the gradual decrease in unexpectedness of US omissions across trials, in contrast to the discrete omission events during partially reinforced reacquisition. **B:** Schematic of the experimental setup during 7 T MRI. Participants lay supine in the scanner and viewed visual conditioned stimuli (CS; geometric shapes) displayed on a screen at the end of the bore via a mirror system. The eyes were simultaneously imaged with an eye-tracking system. Electrodermal activity (EDA) was recorded from the left hand (blue/purple cables). The unconditioned stimulus (US; electrical stimulation) was delivered to the right hand (red cable), and responses were recorded using a button box.

We used a reinforcement-learning-based deep learning model fitted to group-averaged trial-by-trial skin conductance response (SCR) data to compute per-trial prediction errors. SCRs reflect autonomic arousal linked to stimulus salience and expectancy. Averaged SCRs were used here to capture predictive learning about the CS-US association at the population level, allowing us to derive model-based prediction error values. 7T fMRI allowed us to assess VTA activation.

Results showed activation in the cerebellum and VTA for all four phases related to unexpected US omissions, despite differences in learning contexts between phases. Results also showed significant functional interaction between the cerebellum and VTA related to prediction errors in unexpected omission trials. These findings suggest that the cerebellum contributes to extinction learning via the VTA, and thus supports safety learning inhibiting the initial fear association.

## Results

We first report behavioral results demonstrating that participants acquired and partially extinguished conditioned fear in the three-day fear conditioning paradigm (Figure 1). We then present fMRI results related to the prediction, presentation and unexpected omission of the US in the cerebellar cortex, deep cerebellar nuclei (DCN), and VTA. Finally, we report analyses of functional connectivity between the cerebellum and the VTA.

### Behavioral data

Behavioral responses were assessed using skin conductance responses (SCRs; Figure 2), pupil size responses (PSRs; Figure 3), and self-reported ratings. Results for all three measures are reported below, with non-parametric ANOVA-type statistics and descriptive summaries of self-reported ratings provided in the Supplementary Information (Table S1-S6).

**Figure 2:**
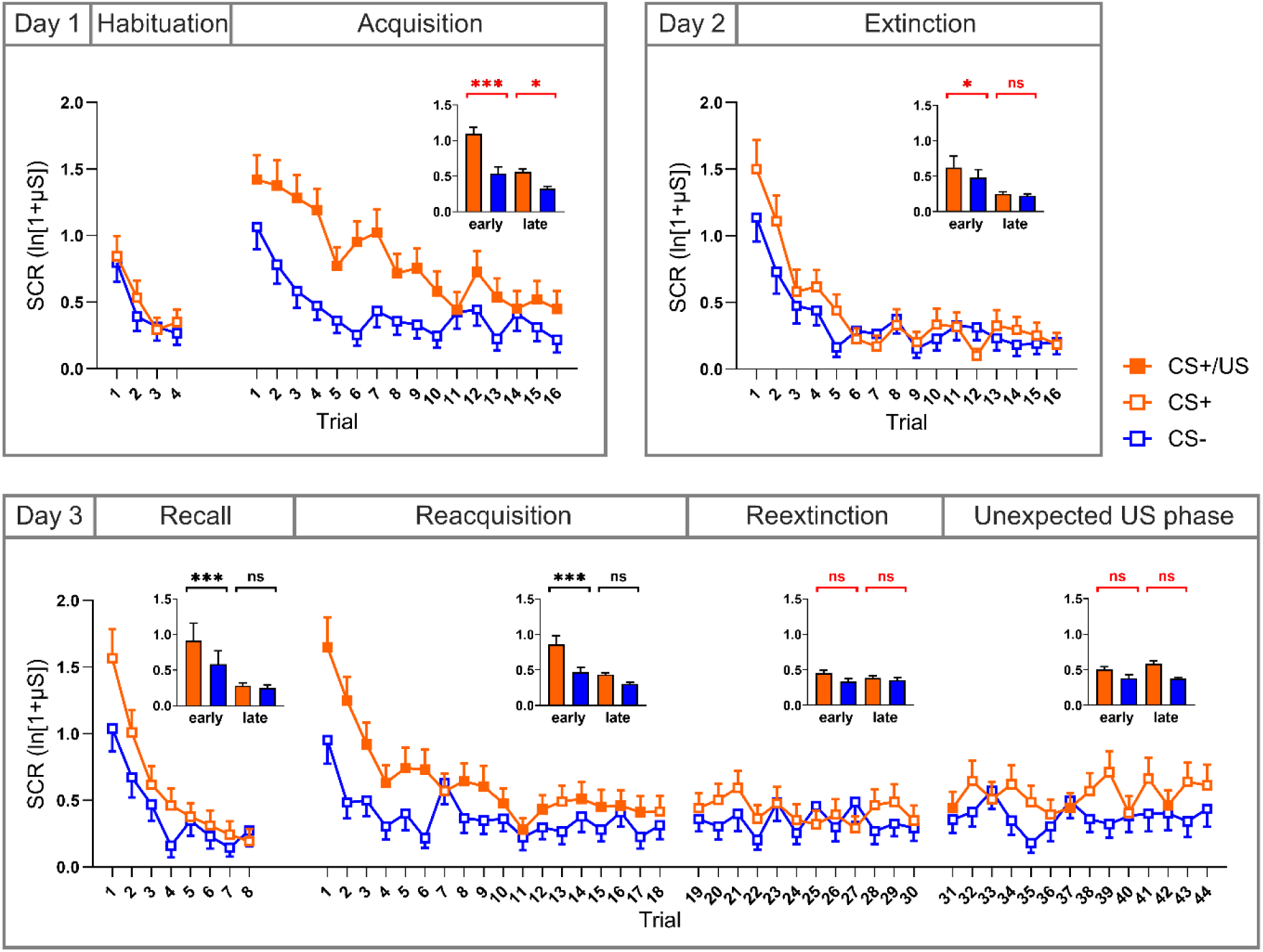
Skin conductance responses (SCRs) for each trial, with CS+ (shown in orange) and CS- (shown in blue) responses paired in blocks. Reinforcement of the CS+ by a US (CS+/US) is indicated by filled squares. The CS- is never reinforced. Bar plots on the top right show mean SCRs for early and late trials, defined as the first and second halves of each phase, respectively. Significance markers indicate post hoc comparisons between CS+ and CS- within early or late trials; markers are shown in black when the Stimulus x Time interaction was significant and in red when the interaction was not significant. On day 1, there was no differentiation between CS+ and CS- in the habituation phase, with significant differentiation emerging during acquisition training. On day 2, differentiation was apparent in early extinction trials and was no longer evident later in extinction (trend-level effect; non-significant Stimulus x Time interaction). On day 3, during the initial recall test, participants exhibited renewed CS+/CS- differentiation, consistent with spontaneous recovery. During initial reacquisition, there were again differential responses to the CS+ and CS-, which appeared reduced in reextinction and the unexpected US phase. Mean values are shown with error bars representing the standard error of the mean. CS: Conditioned stimulus; US: Unconditioned stimulus; SCR: Skin conductance response; μS: microsiemens.

**Figure 3:**
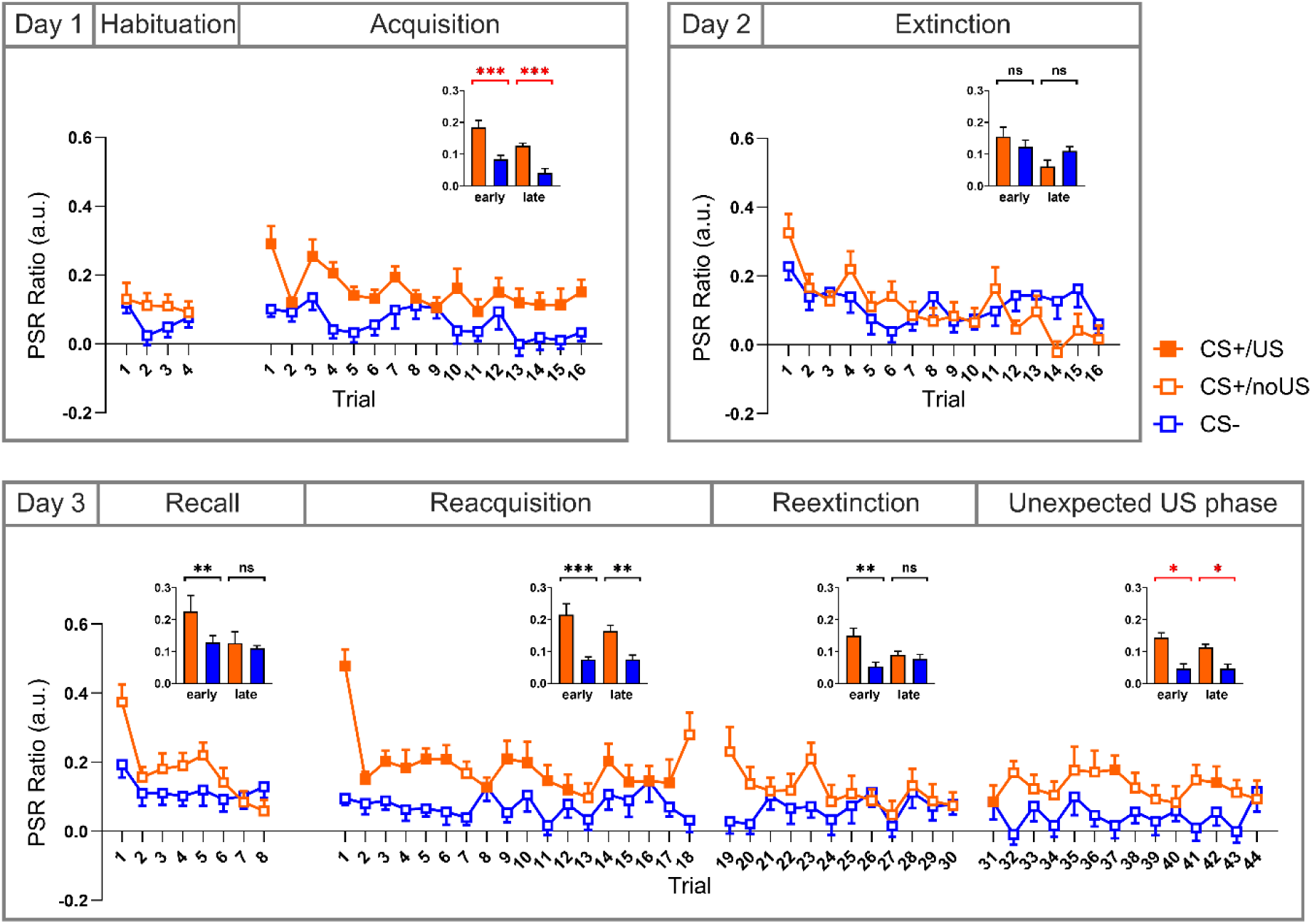
Pupil size responses (PSRs) for each trial, with CS+ (shown in orange) and CS- (shown in blue) responses paired in blocks. Reinforcement of the CS+ by a US (CS+/US) is indicated by filled squares. The CS- is never reinforced. Bar plots on the top right show mean PSRs for early and late trials, defined as the first and second halves of each phase, respectively. Significance markers indicate post hoc comparisons between CS+ and CS- within early or late trials; markers are shown in black when the Stimulus x Time interaction was significant and in red when the interaction was not significant. On day 1, there was no differentiation between CS+ and CS- in the habituation phase, with significant differentiation emerging during acquisition training. On day 2, a significant Stimulus x Time interaction was observed during extinction, reflecting a decrease from early to late extinction trials for the CS+ but not for the CS-, despite the absence of significant post hoc CS+/CS- differences. On day 3, during the initial recall test, participants exhibited renewed CS+/CS- differentiation, consistent with spontaneous recovery. During initial reacquisition, there were again differential responses to the CS+ and CS-, which appeared reduced in reextinction and the unexpected US phase. Mean values are shown with error bars representing the standard error of the mean. CS: Conditioned stimulus; US: Unconditioned stimulus; PSR: Pupil size response.

### Skin conductance responses (SCRs)

During habituation (day 1), SCRs did not differ between CS+ and CS- (Figure 2, top row). SCRs for both stimuli decreased significantly across habituation trials. Non-parametric ANOVA-type statistics revealed a significant effect for Time (early vs. late; *F_1_* = 26.49; *p* < 0.001) but not for Stimulus (CS+ vs. CS-; *F_1_* = 1.19; *p* = 0.276) or the Stimulus x Time interaction (*F_1_* = 0.06; *p* = 0.802). During fear acquisition training (day 1), participants underwent a fully reinforced, semi-instructed differential fear conditioning protocol. Participants quickly learned the associations, and their responses decreased as expected due to habituation effects. SCRs for the CS+ were significantly higher than for the CS-, and decreased in late trials. Non-parametric ANOVA-type statistics revealed a significant effect of Stimulus (CS+ vs. CS-; *F_1_* = 20.79; *p* < 0.001) and Time (early vs. late; *F_1_* = 28.75; *p* < 0.001). No significant Stimulus x Time interaction was observed (*F_1_* = 3.80; *p* = 0.051).

During extinction training (day 2), SCRs were higher for the CS+ than for the CS- only in initial extinction trials. Non-parametric ANOVA-type statistics revealed a significant effect of Stimulus (CS+ vs. CS-; *F_1_* = 7.71; *p* = 0.006) and Time (early vs. late; *F_1_* = 21.10; *p* < 0.001), whereas the Stimulus x Time interaction was not significant (*F_1_* = 3.74; *p* = 0.053). Exploratory post hoc comparisons indicated higher SCRs to the CS+ than to the CS- during early extinction trials (*p* = 0.017), but no differentiation during late extinction (*p* = 0.798). These post hoc results are reported descriptively, as the Stimulus x Time interaction did not reach statistical significance.

During the recall test (day 3), SCRs for CS+ were significantly higher than for CS- (Figure 2, bottom row). In late recall, no difference between SCRs for CS+ and CS- was observed. Non-parametric ANOVA-type statistics revealed a significant main effect of Stimulus (CS+ vs. CS-; *F_1_* = 4.99; *p* = 0.026), Time (early vs. late; *F_1_* = 36.74; *p* < 0.001) and a Stimulus x Time interaction (*F_1_* = 5.53; *p* = 0.019). Post hoc tests revealed that the CS+ had a significantly higher SCR than the CS- in early trials (*p* < 0.001) but not in late trials (*p* = 1.000). SCRs for both stimuli significantly decreased in late trials compared to early trials (both: *p* < 0.001).

During reacquisition on day 3, SCRs for the CS+ were significantly higher than for the CS-, and decreased for both stimuli in late reacquisition. Non-parametric ANOVA-type statistics revealed a significant effect of Stimulus (CS+ vs. CS-; *F_1_* = 27.20, *p* < 0.001), Time (early vs. late; *F_1_* = 35.44, *p* < 0.001) and a Stimulus x Time interaction (*F_1_* = 6.80, *p* = 0.009). Post hoc tests showed that both stimuli were higher in early vs late reacquisition training (CS- early vs. CS-late; *p* = 0.007; CS+ early vs. CS+ late; *p* < 0.001), and a significant difference between stimulus types during early trials (*p* < 0.001), but no significant difference in late trials (*p* = 0.094).

During reextinction on day 3, SCRs related to the CS+ were significantly higher than for the CS-although the difference was small. Non-parametric ANOVA-type statistics revealed a significant main effect of Stimulus (CS+ vs. CS-; *F_1_* = 4.47; *p* = 0.035), but neither for Time (early vs. late; *F_1_* = 0.08; *p* = 0.780) nor for the Stimulus x Time interaction (*F_1_* = 0.95; *p* = 0.329).

In the last phase on day 3, three CS+ trials were unexpectedly reinforced (“unexpected US phase”). SCRs for the CS+ were significantly higher than for the CS-. Non-parametric ANOVA-type statistics revealed a significant main effect of Stimulus (CS+ vs. CS-; *F_1_* = 5.43; *p* = 0.020), but neither for Time (early vs. late; *F_1_* = 0.29; *p* = 0.593) nor for the Stimulus x Time interaction (*F*_1_ = 2.34; *p* = 0.126).

### Pupil size responses (PSRs)

During habituation on day 1, PSRs related to the CS+ and CS- did not differ significantly (non-parametric ANOVA-type statistics; all *p* > 0.199).

During fear acquisition training on day 1, PSRs were significantly higher in CS+ trials than in CS- trials, and PSR was more pronounced in early compared to late trials (Figure 3, top row). Non-parametric ANOVA-type statistics revealed a significant main effect of Stimulus (CS+ vs. CS-; *F_1_* = 39.51; *p* < 0.001) and Time (early vs. late; *F_1_* = 24.31; *p* < 0.001). No significant Stimulus x Time interaction was observed (*F_1_* = 0.58; *p* = 0.446).

During extinction training on day 2, PSRs for the CS+ decreased from early to late extinction, but there were no differences between early and late trials for the CS-. Non-parametric ANOVA-type statistics revealed a significant main effect of Time (early vs. late; *F_1_* = 18.21; *p* < 0.001) and a significant Stimulus x Time interaction (*F_1_* = 4.28; *p* = 0.039). No significant main effect of Stimulus was observed (CS+ vs. CS-; *F_1_* = 1.12; *p* = 0.290). Post hoc tests showed that PSRs related to the CS+ were significantly lower during late extinction compared to early (*p* < 0.001). This result was not observed for PSRs related to the CS- (*p* = 0.643).

During the recall test on day 3, PSRs for the CS+ were significantly higher than for the CS-during early, but not late, recall trials (Figure 3, bottom row). Non-parametric ANOVA-type statistics revealed a significant effect of Stimulus (CS+ vs. CS-; *F_1_* = 6.59; *p* = 0.010) and Time (early vs. late; *F_1_* = 6.41; *p* = 0.011) and a significant Stimulus x Time interaction (*F_1_* = 4.87; *p* = 0.027). Post hoc tests showed higher PSRs for the CS+ than for the CS- during early recall (*p* = 0.017), but not during late recall (*p* = 0.862), and that the PSRs related to the late CS+ were significantly lower than early (*p* = 0.003)

During reacquisition on day 3, PSRs for the CS+ decreased from early to late trials, whereas no change over time was observed for the CS-. PSRs differed between CS+ and CS- in both early and late trials. Non-parametric ANOVA-type statistics revealed significant effects of Stimulus (CS+ vs. CS-; *F₁* = 47.87, *p* < 0.001) and Time (early vs. late; *F₁* = 4.54, *p* = 0.033), as well as a significant Stimulus x Time interaction (*F₁* = 7.70, *p* = 0.006). Post hoc tests indicated that PSRs for the CS+ were higher early than late (*p* = 0.004), whereas no such change was observed for the CS- (*p* = 0.974).

During reextinction on day 3, PSRs for CS+ were significantly higher than for CS- early, but the difference became insignificant in late trials. Non-parametric ANOVA-type statistics revealed a significant main effect of Stimulus (CS+ vs. CS-; *F_1_* = 9.10; *p* = 0.003) and a Stimulus x Time interaction (*F_1_* = 4.43; *p* = 0.035), but not a significant main effect of Time (early vs. late; *F_1_* = 0.83, *p* = 0.364). Post hoc tests showed a significant difference between the CS+ and CS- early but not late (early: *p* = 0.007; late: *p* = 0.775).

During the unexpected US phase on day 3, PSRs for the CS+ were higher than for the CS-. Non-parametric ANOVA-type analysis revealed a significant main effect of Stimulus (CS+ vs. CS-; *F_1_* = 17.35; *p* < 0.001), but not of Time (early vs. late; *F_1_* = 0.83; *p* = 0.363) and the Stimulus x Time interaction (*F_1_* = 0.03; *p* = 0.872)

### Self-reports

Valence, arousal, fear and US expectancy ratings of the CS+ did not differ significantly from the CS- before acquisition training (Table S5 and S6). Following acquisition training, the CS+ was rated as more arousing, fearful and unpleasant compared to the CS-, and participants expected the US to follow the CS+ but not the CS- (all *p* < 0.001). Although less pronounced, the CS+/CS- differentiation persisted after extinction training and recall test (all *p* < 0.004). However, extinction and recall CS+ ratings were significantly reduced compared to acquisition (all *p* ≤ 0.001), whereas CS- ratings remained unchanged (all *p* > 0.532), reflecting extinction learning. The difference between CS+ and CS- increased at the end of day 3, when CS+/US pairings were reintroduced.

Non-parametric ANOVA-type statistics showed a significant main effect of Stimulus, Time and a Stimulus x Time interaction (main effects and interaction; all *p* < 0.001; Table S6). Post hoc tests revealed significant differences between stimulus type (CS+, CS-) following acquisition training, extinction training, recall test and at the end of day 3 but not after habituation.

## fMRI results

Analysis of BOLD fMRI responses revealed significant activations related to the prediction, the presentation and the unexpected omission of the US (Figure 4-7). Our primary focus was on activations and connectivity related to the unexpected omission of the US, which could be detected across four distinct phases: extinction training, recall test, reacquisition, and reextinction (Figure 6-8). Table S7 reports VOI-level summary statistics (mean contrast estimates, 95% confidence intervals, and Cohen’s d) for a cerebellar cortex VOI derived from the conjunction of unexpected US omission contrasts, as well as for the deep cerebellar nuclei (DCN), and VTA across all fMRI contrasts, while Table S8-S12 report cluster-level activation and connectivity results.

**Figure 4:**
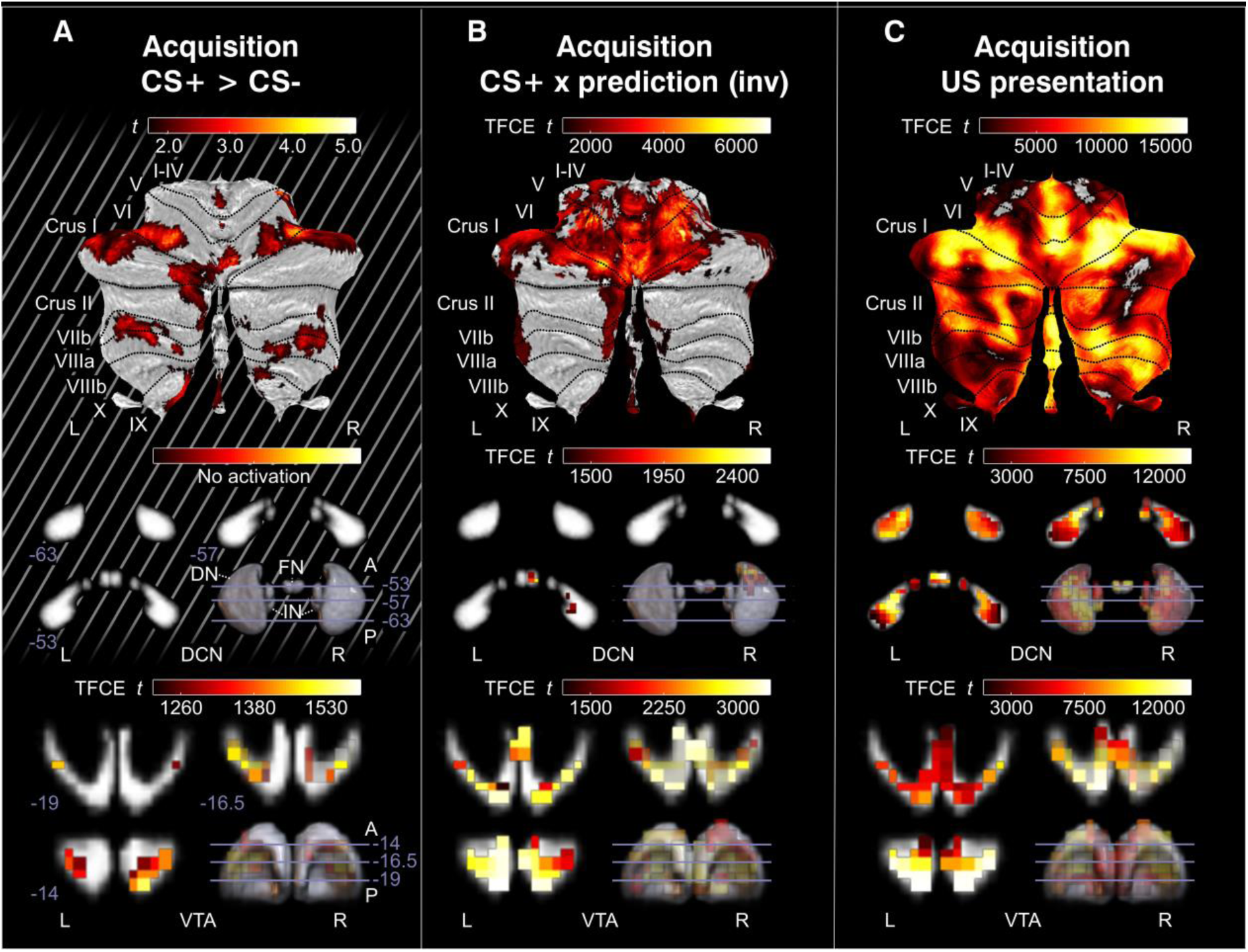
Cerebellar cortex, DCN, and VTA fMRI activations related to prediction and presentation of the US during acquisition training. Cerebellar cortex activations are shown on cerebellar flatmaps (SUIT^22^, while VTA and DCN activations are displayed on coronal slices progressing from posterior to anterior, with MNI y-coordinates indicated at the lower left of each slice. A 3D rendering in the bottom right shows slice locations within the probabilistic DCN and VTA atlases (see Methods for atlas generation). Trend-level results are indicated by gray diagonal lines in the background of the panels. **A:** The event-based CS+ > CS- prediction contrast showed trend-level activations in posterolateral cerebellar lobules and the VTA; no DCN activations were observed. **B:** Inverse activations were observed in the parametric modulation analysis of CS+ × prediction, with activations mainly in the anterior cerebellum, paravermis, lobule VI and Crus I, and the VTA, with trend-level activations in the DCN. **C:** Presentation of the US elicited widespread activations in the cerebellar cortex, DCN, and VTA. VTA: ventral tegmental area; DCN: deep cerebellar nuclei; DN: dentate nucleus; IN: interposed nucleus; FN: fastigial nucleus; CS: conditioned stimulus; US: unconditioned stimulus; MNI: Montreal Neurological Institute standard brain; SUIT: spatially unbiased atlas template of the cerebellum; TFCE t: threshold-free cluster-enhanced test statistic.

**Figure 5:**
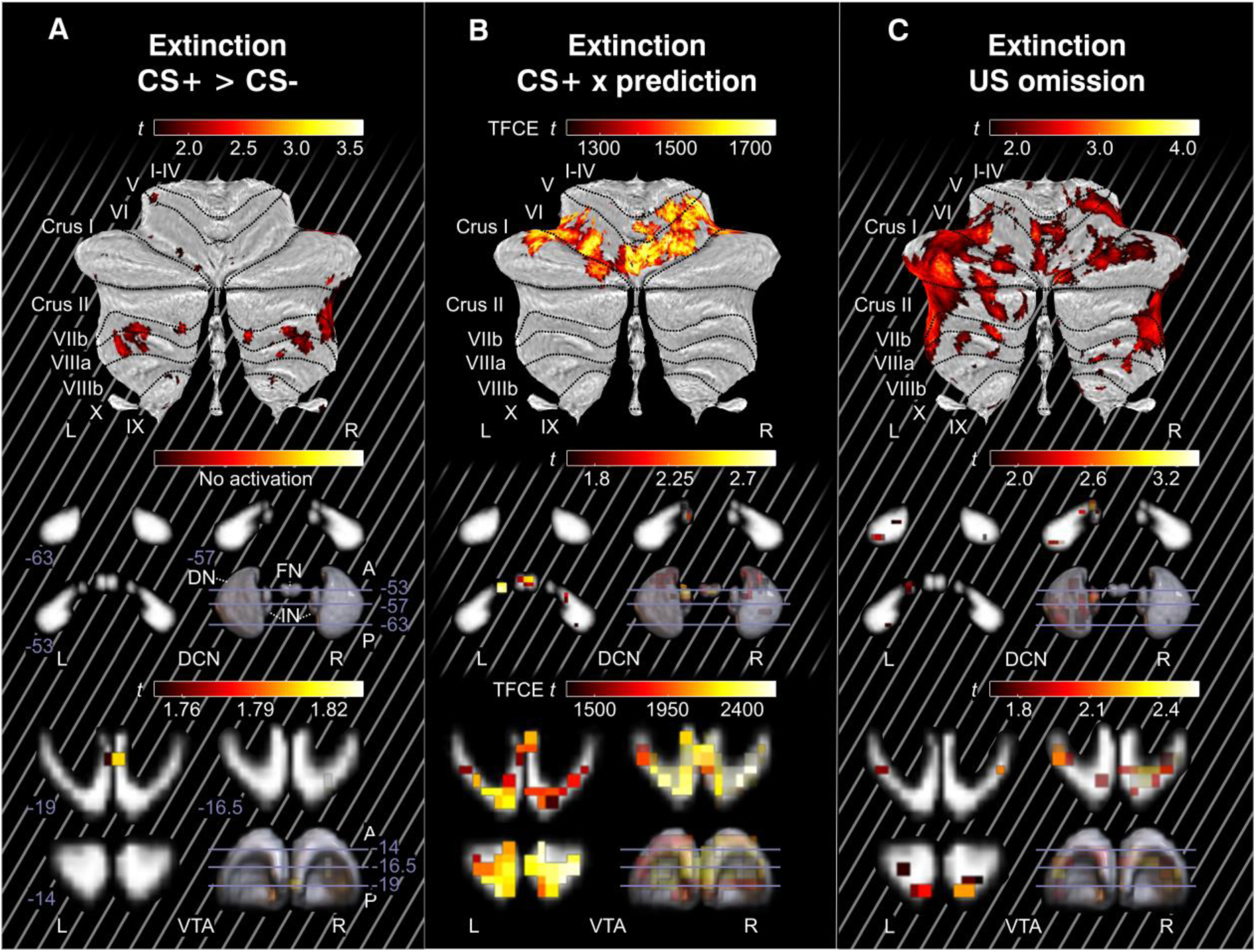
Cerebellar cortex, DCN and VTA fMRI activations related to the prediction and presentation of the US. Cerebellar cortex activations are shown on cerebellar flatmaps (SUIT^22^), while VTA and DCN activations are displayed on coronal slices progressing from posterior to anterior with MNI y-coordinates indicated on the left-bottom for each slice. A 3D rendering in the bottom right shows the slice locations within the probabilistic DCN and VTA atlases (see methods for atlas generation). Trend-level results are indicated by gray diagonal lines in the background of the panels. **A:** The event-based CS+ > CS- contrast showed trend-level activations in lobules VIIb, VIIIa and VIIIb and the central VTA, with no DCN activations. **B:** The parametric modulation contrast CS+ x prediction showed activations in lobule VI and Crus I, the vermis, and the VTA, as well as trend-level activations in the DCN. **C:** Trend-level activations related to the omission of the US were observed in the cerebellar cortex, DCN and VTA. VTA: ventral tegmental area; DCN: deep cerebellar nuclei; DN: dentate nucleus; IN: interposed nucleus; FN: fastigial nucleus; CS: conditioned stimulus; US: unconditioned stimulus; MNI: Montreal Neurological Institute standard brain; SUIT: spatially unbiased atlas template of the cerebellum; TFCE t: threshold-free cluster-enhanced test statistic.

**Figure 6:**
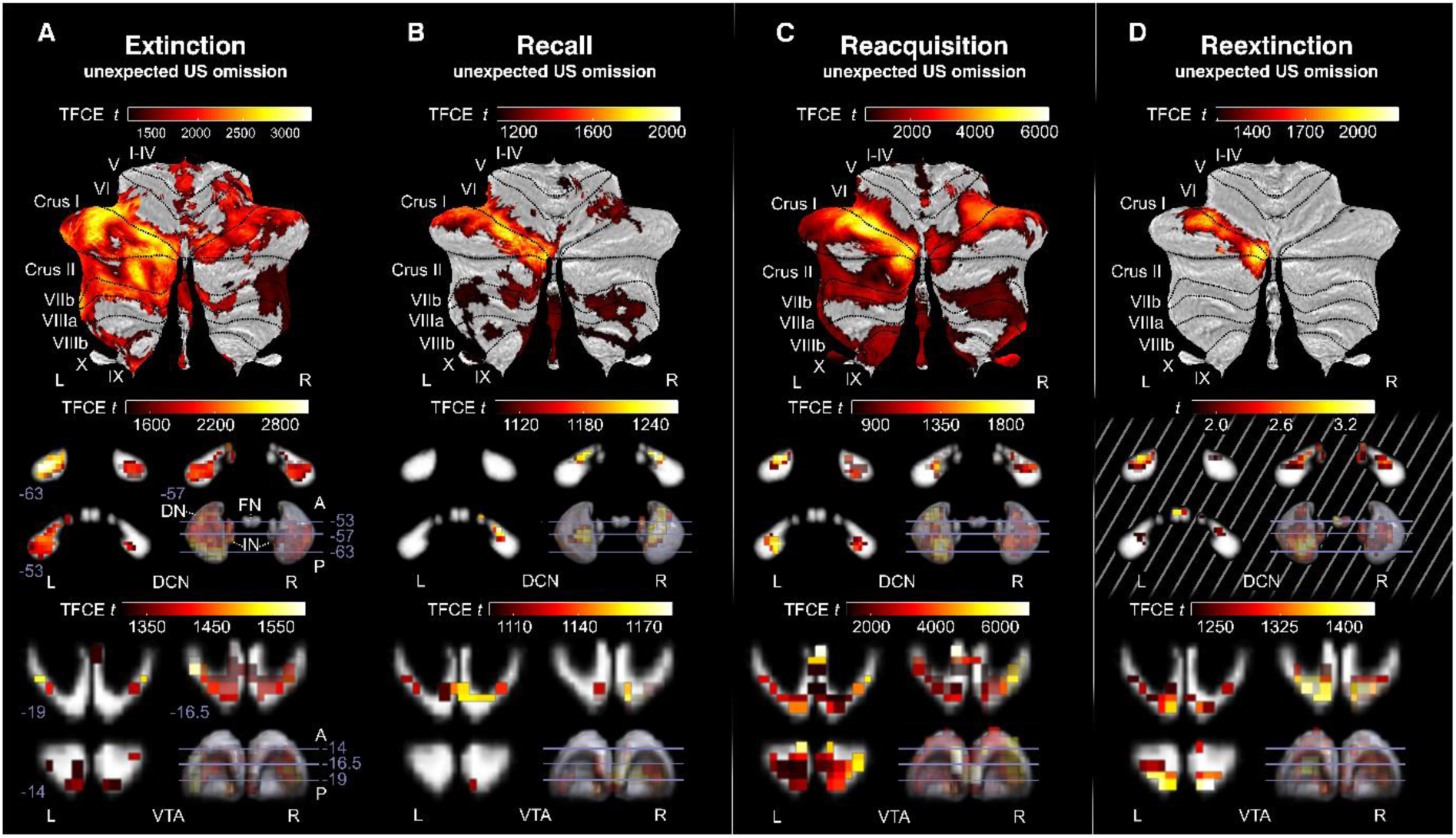
Cerebellar cortex, DCN and VTA fMRI activations to the unexpected omission of the US (event-based analysis). Cerebellar cortex activations are shown on cerebellar flatmaps (SUIT^22^), while VTA and DCN activations are displayed in coronal slices progressing from posterior to anterior with MNI y-coordinates indicated on the left-bottom for each slice. A 3D rendering in the bottom right shows the slice locations within the probabilistic DCN and VTA atlas (see Methods for atlas generation). Trend-level results are indicated by gray diagonal lines in the background of the panels. Event-based contrasts of the first three unexpected omissions showed activations in the cerebellar cortex, DCN and VTA for all four phases. Activations in the DCN during reextinction were trend-level **(D)**. Activations were most consistent in left lobule VI and Crus I, with DCN activation mainly in the left dentate, while VTA activations were bilateral. VTA: ventral tegmental area; DCN: deep cerebellar nuclei; DN: dentate nucleus; IN: interposed nucleus; FN: fastigial nucleus; CS: conditioned stimulus; US: unconditioned stimulus; MNI: Montreal Neurological Institute standard brain; SUIT: spatially unbiased atlas template of the cerebellum; TFCE t: threshold-free cluster-enhanced test statistic.

**Figure 7:**
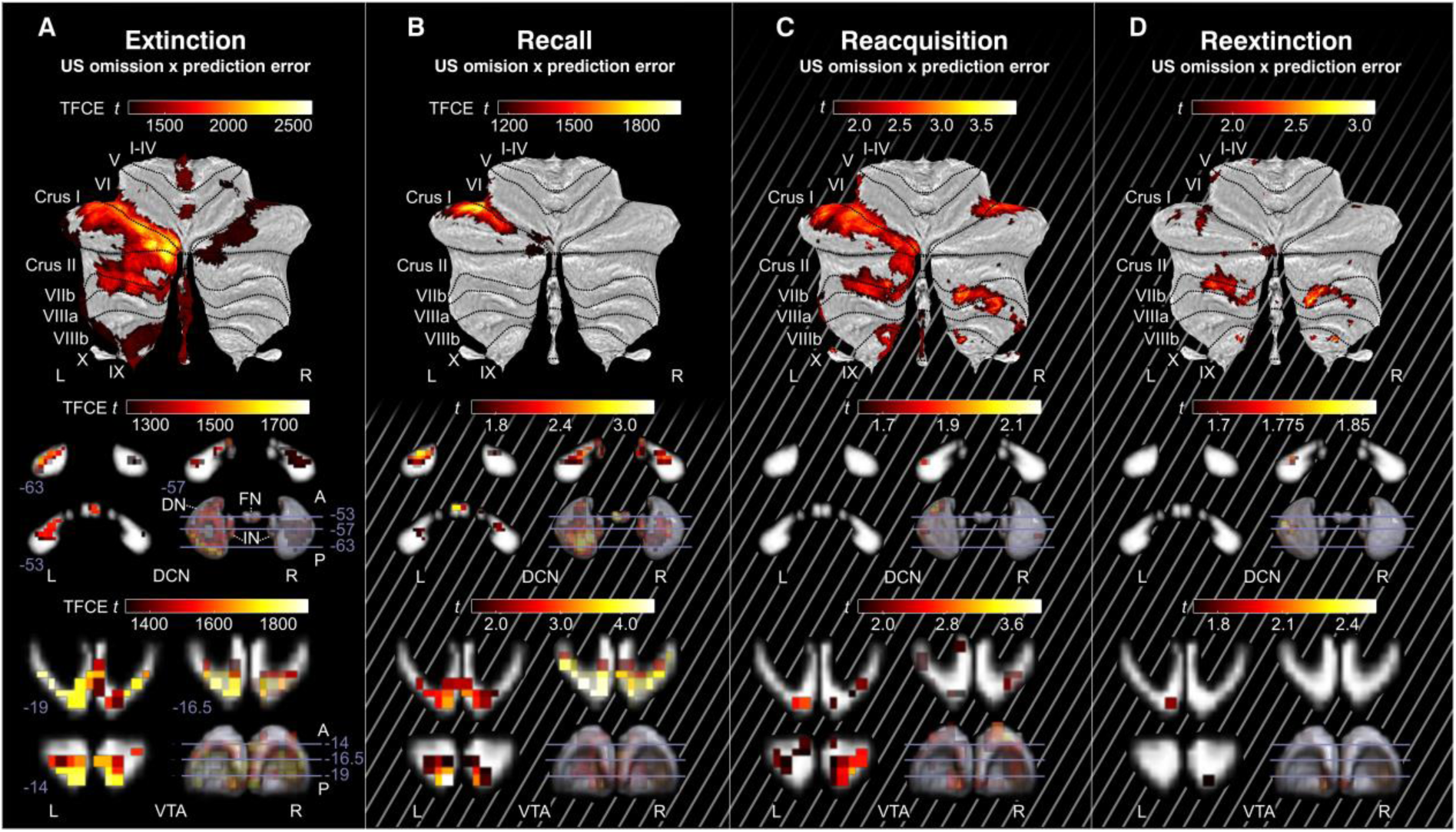
Cerebellar cortex, DCN and VTA fMRI activations to the unexpected omission of the US. Cerebellar cortex activations are shown on cerebellar flatmaps (SUIT^22^), while VTA and DCN activations are displayed in coronal slices progressing from posterior to anterior with MNI y-coordinates indicated on the left-bottom for each slice. A 3D rendering in the bottom right shows the slice locations within the probabilistic DCN and VTA atlas (see methods for atlas generation). Trend-level results are indicated by gray diagonal lines in the background of the panels. Parametric modulation contrasts related to prediction errors from unexpected US omissions showed activations in all four phases in the cerebellum, DCN and VTA. Activations during the recall test in the DCN and VTA were trend-level (B). Activations in reacquisition and reextinction were trend-level (C, D). Activations were most consistently found in lobule VI and Crus I. In addition, activations were found in the vermis, Crus II and lobule VIIb. VTA: ventral tegmental area; DCN: deep cerebellar nuclei; DN: dentate nucleus; IN: interposed nucleus; FN: fastigial nucleus; CS: conditioned stimulus; US: unconditioned stimulus; MNI: Montreal Neurological Institute standard brain; SUIT: Spatially Unbiased Atlas Template of the cerebellum; TFCE t: threshold-free cluster-enhanced test-statistic.

**Figure 8:**
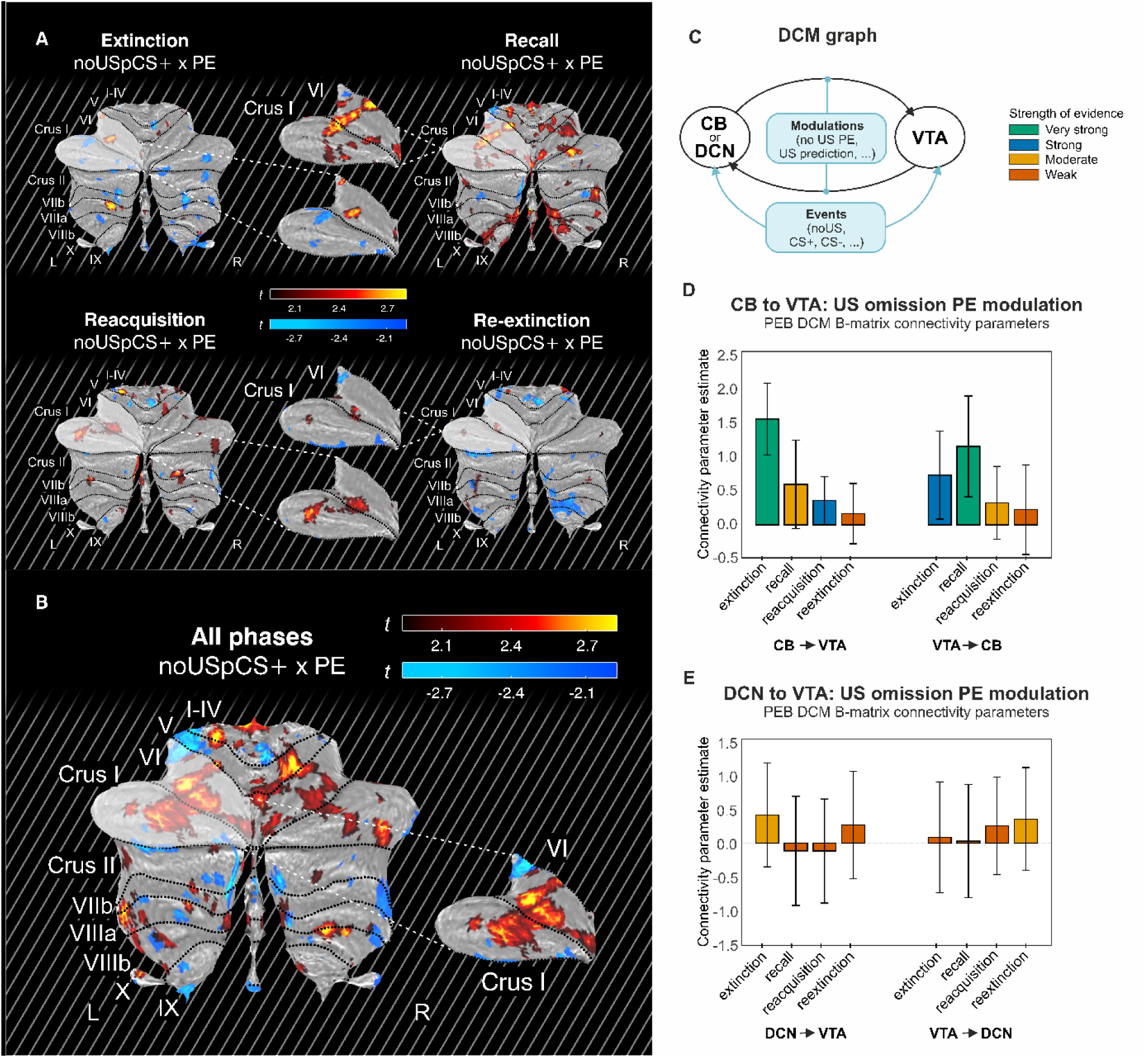
Functional connectivity analysis using PPI and DCM. **A,B:** Trend-level activations in the cerebellum found by PPI analysis using the VTA as a seed region, shown on cerebellar flatmaps^22^. To highlight activations, a zoomed cutout of lobule VI and Crus I is also shown. Trend-level results are indicated by gray diagonal lines in the background of the panels. **A:** Trend-level connectivity with VTA is found in lobule VI and Crus I for extinction training, the recall test, reacquisition and reextinction using PPI of the unexpected US omission contrasts from the parametric modulation analysis with a VTA seed. **B:** Summary of trend-level PPI results by summing all contrasts shown in panel A. **C-E:** DCM analysis showed significant modulation of cerebellar cortex (CB) to VTA connections with prediction errors during unexpected omission events. **C:** DCM model is shown, modulations act on both the CB/DCN to VTA as well as the VTA to CB/DCN connections. Events, i.e., US post CS+, no US post CS+, no US post CS-, CS+ and CS- for each phase, are provided as inputs to both nodes. Colors of bars are determined by the posterior probability calculated in the PEB analysis (very strong: P > 0.99; strong: 0.95 < P < 0.99; moderate: 0.75<P < 0.95; weak: P < 0.75). Very strong and strong results are considered significant, moderate results are considered trend-level. **D:** Significant modulations during extinction training, the recall test and reacquisition. The strongest result was found during extinction training in the connection from the cerebellum to VTA. **E:** Trend-level results showed connectivity from the DCN to VTA in extinction, and from the VTA to DCN in reextinction. PPI: Psychophysiological interaction; DCM: Dynamic causal modeling; PEB: Parametric Empirical Bayes; PE: Prediction error; P: Posterior probability; VTA: Ventral tegmental area; CB: Cerebellar region in lobule VI and Crus I consistently active during unexpected US omissions; US: unconditioned stimulus; L: left; R: right; SUIT: spatially unbiased atlas template of the cerebellum; t: test-statistic.

### Activations related to US prediction (CS+ > CS- and CS+ x prediction)

In the event-based analysis, there was no significantly higher activation in the cerebellum during the CS+ compared to the CS- (CS+ > CS-) during acquisition or extinction training. In the same contrast, the lateral VTA exerted significant differential activations during acquisition (Figure 4A; threshold free cluster enhancement (TFCE) and family-wise error (FWE) corrected p < 0.05), but not during extinction training (Figure 5A). Trend-level results showed differential cerebellar activation in posterolateral regions (lobule VI, Crus I, Crus II, lobule VIIb and VIIa; Figure 4A and Figure 4D; uncorrected p < 0.05) during acquisition training, which match results obtained by Ernst and colleagues (2019).

In the parametric modulation analysis, there were significant activations in the cerebellum (lobule and vermis I-VI), DCN and VTA negatively associated with CS+ prediction value modulation during acquisition training (CS+ x prediction; Figure 4B), whereas there were significant positively associated activations during extinction training (Figure 5B; lobule VI and Crus I bilaterally, right lobule V and VTA). Parametric modulation results indicate that early CS+ presentations are paired with high cerebellar activations in both acquisition and extinction training, with prediction values being low during early and high during late acquisition, whereas prediction values are high during early and low during late extinction (Figure 9). In other words, the activations were negative due to high activations of early CS+ compared to late CS+ events with prediction values taking the opposite course (i.e., low during early acquisition and high during late acquisition).

**Figure 9:**
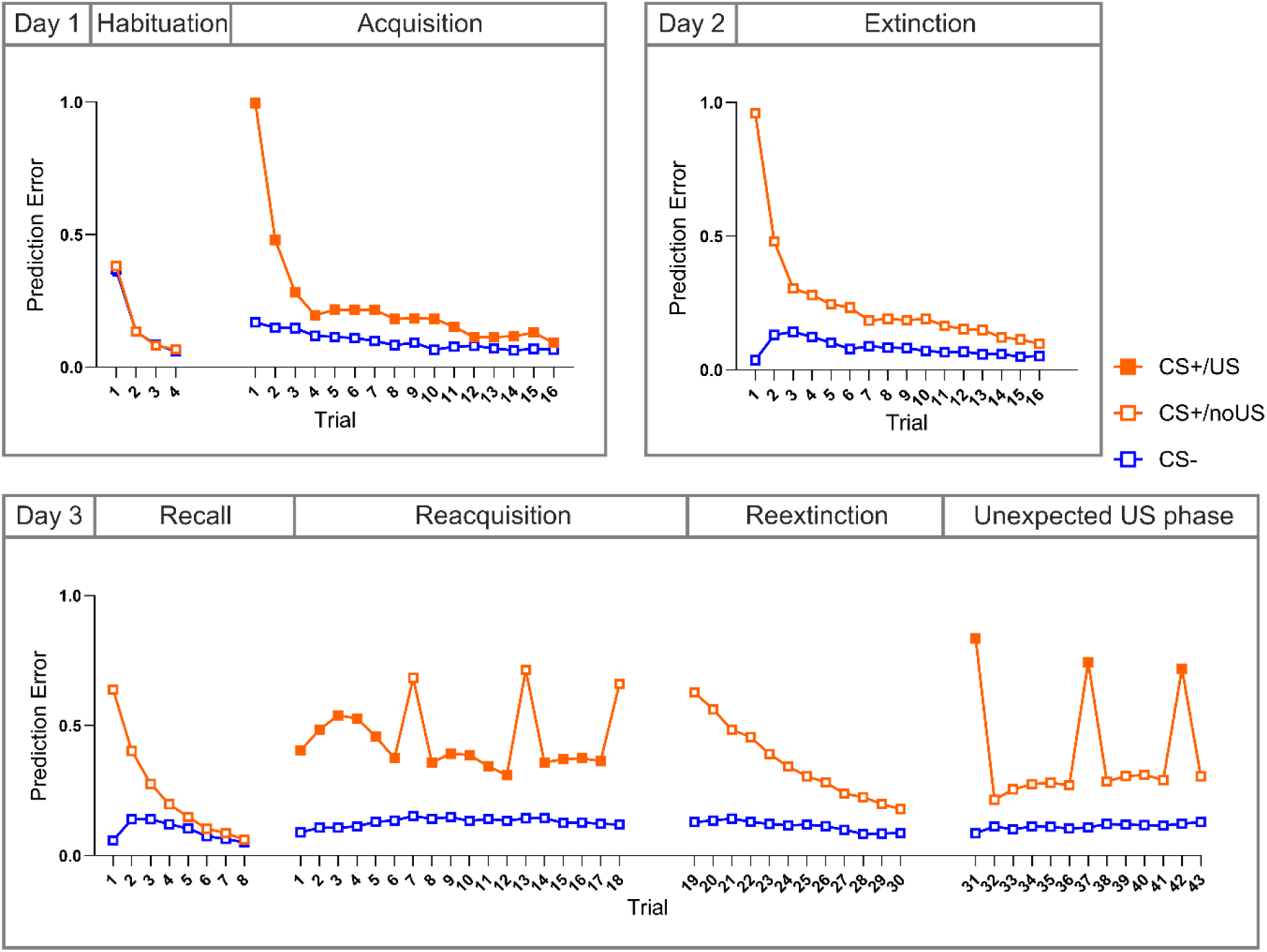
Mean absolute prediction error values for each trial estimated from SCRs, with CS+ and CS- responses paired in blocks. Reinforcement of the CS+ by a US (CS+/US) is indicated by filled squares. Prediction and prediction error values were estimated over 200 iterations. CS: conditioned stimulus; US: unconditioned stimulus.

### Activations related to US presentation (US post CS+ > no US post CS-)

During acquisition training, significant activations related to the presentation of the US (US post CS+ > no US post CS-) were widespread and included the whole cerebellum and VTA (Figure 4C). Cerebellar activations extended to the DCN. Peak activations in the cerebellum were found ipsilateral to the US (electrical stimulation applied to right hand).

### Activations related to the unexpected omission of the US (first 3 no US post CS+ > no US post CS- and no US post CS+ x prediction error)

Considering the whole extinction training phase, trend-level results showed activations in posterolateral lobules and the VTA related to the omission of the US (no US post CS+ > no US post CS-; Figure 5C). When focusing specifically on the early trials in extinction training to quantify activations related to the unexpected omission of the US, we found significant activations in both event-based and parametric modulation analyses.

In the event-based analysis, the cerebellar cortex, DCN and VTA showed significant activations related to unexpected omissions of the US in initial trials for all four phases (first 3 no US post CS+ > no US post CS-; extinction training, recall test, reacquisition and reextinction; Figure 6). The location of activations was consistent across phases, with highest activations in left posterolateral lobule VI and Crus I. Activations in the left posterolateral cerebellar cortex were paired with high activations in the left dentate nucleus. The VTA was bilaterally activated.

In the parametric modulation analysis, significant activations were found in the cerebellum for unexpected omissions during extinction training and the recall test (Figure 7A and Figure 7B), and showed trend-level activations during reacquisition and reextinction (Figure 7C and Figure 7D). The highest activations were again found in left lobule VI and Crus I. In the DCN, significant activations were found during extinction training with peak activations in the left dentate, with trend-level results in the recall test. In the VTA, results were significant during extinction training (Figure 7A), with trend-level activations during the recall test, reacquisition and reextinction.

### PPI: Functional connectivity during unexpected US omissions between the cerebellum and VTA

Next, we aimed to assess functional interactions between the cerebellum and VTA using psychophysiological interaction (PPI), using the VTA as a seed region. In the parametric modulation analysis, we observed trend-level positive functional connectivity in left lobule VI and Crus I for extinction training, recall test, reacquisition and reextinction (Figure 8A). When combining the contrasts by summing, the combination of all four phases showed similar positive functional connectivity between the cerebellar cortex and the VTA in lobule VI and Crus I, but only on a trend-level (Figure 8B). No activations were seen in the DCN.

To assess whether similar connectivity patterns were observed when using an alternative seed, we performed an additional PPI analysis using a cerebellar region of interest defined from the conjunction analysis of unexpected US omissions (see Volumes of interest (VOI) definition in Methods and Figure S4) as the seed. In this analysis, connectivity effects in the VTA were very limited and restricted to a small number of voxels at an uncorrected threshold (Figure S3).

### DCM: Functional connectivity during unexpected US omissions between the cerebellum and VTA

In the dynamic causal modeling (DCM) parametric modulation analysis, unexpected omissions significantly positively modulated both the connection from cerebellar cortex to VTA and VTA to cerebellar cortex during extinction training (Figure 8D; very strong evidence, posterior probability *P* > 0.99, strong evidence: 0.95 < *P* < 0.99). In the recall test, the VTA to cerebellar cortex connection was positively modulated. In reacquisition, the cerebellar cortex to VTA connection was positively modulated. Results were less strong for the other unexpected omission modulations, with trend-level moderate evidence for the cerebellar cortex (CB) to VTA connection in the recall test and the VTA to CB connections in reacquisition, and weak evidence for both CB to VTA and VTA to CB connections in reextinction (Figure 8D; moderate 0.75 < *P* < 0.95; weak *P* < 0.75). In the model incorporating the DCN and VTA, there was a trend-level modulation from the DCN to the VTA during extinction training, as well as a trend-level modulation from the VTA to the DCN during reextinction (Figure 8E; moderate significance, 0.75 < *P* < 0.95).

## Discussion

The cerebellum and VTA were active during unexpected omissions of aversive unconditioned stimuli in the initial extinction training trials and in other learning phases. Notably, this activation was accompanied by increased functional connectivity between the cerebellum and VTA. Given that the unexpected omission of an aversive stimulus can be considered rewarding, our data suggest that interactions between the cerebellum and the VTA contribute to reward-like prediction error processing during fear extinction learning in humans. Such interactions may represent one mechanism by which the cerebellum contributes to non-motor functions, in this case the control of emotions. Our findings build on previous research in rodents showing that the cerebellum modulates addictive, social and depression-like behavior via direct projection to VTA^16,23^.

Cerebellar activations were most prominent in trials with maximum prediction errors, i.e., during the initial extinction training phase, but were also seen in early recall, reacquisition and early reextinction. The activations in lobule VI and Crus I are consistent with previous findings by Ernst et al.^18^ during unexpected omissions in a partially reinforced acquisition phase.

Additionally, other studies have reported activations in these regions during the prediction of the US in extinction training^24–27^ and during unexpected omissions of aversive stimuli^28–31^. More broadly, activations in lobule VI and Crus I have been observed during US prediction^32^, particularly in the early phases of acquisition training. Conversely, unexpected rewards lead to activation in Crus I^33^, and a meta-analysis implicated left lobule VI and Crus I in both the anticipation and processing of rewarding outcomes^34^. Anticipation-related activations were more hemispheric, whereas rewarding outcome activations were more paravermal.

Activated areas in more lateral parts of lobule VI and Crus I overlap with multi-demand regions^35^, whereas activations neighboring the vermis overlap with emotional regions^36^. The main components of the multi-demand (or executive) neural network are working memory, attention and inhibition^37^. Thus, attention-related processes involved in fear conditioning may also contribute to cerebellar activations. Attention-related processes have been related to the salience of the stimulus^38^, but are also thought to change when a cue is followed by an unexpected outcome with more attention being paid in the next trial^39,40^. Attention related processes may therefore contribute to activations in the more lateral cerebellar parts of lobule VI/Crus I. However, unexpected omission of the US should lead to increased attention to the CS in the next trial and therefore increased activation towards the CS (which was not the case) and not at the time the US was presented and did not occur.

Our findings of increased activation in the VTA related to the unexpected omission of the aversive US align well with rodent data^6,7^. In rodents, it has been shown that dopamine neurons in the VTA are active in initial extinction trials, and that lack of this dopamine signal prevents extinction learning^5,6,41^. In a more recent study, Salinas-Hernandez et al.^10^ observed that these dopamine neurons project to extinction-related neurons of the nucleus accumbens, and receive input from the dorsal raphe. Our data show that the VTA is also involved in extinction learning in humans, and provide first evidence that the VTA is functionally coupled with the cerebellum during extinction learning. Esser et al.^42^ found that the administration of L-Dopa in healthy human participants enhanced activation of the nucleus accumbens which was associated with VTA activity at the time of unexpected omission of the US in early extinction learning, which is in good agreement with our findings.

The unexpected omission of the aversive US is thought to be rewarding and fMRI signals in the cerebellum and VTA may be related to reward-like prediction errors. Whereas the VTA has long been known to process rewards, and fMRI data in humans indicate that the VTA is active related to the unexpected presentation of rewards^43,44^, the cerebellum has only recently been found to process reward signals in rodents^11,13,45^. fMRI signals are known to be driven by synaptic input^46–48^. In the cerebellum this is mainly related to mossy fiber input to the cerebellar cortex^49^. There is evidence, however, that granule cell activity also modulates vasodilatation and likely contributes to fMRI BOLD signals in the cerebellar cortex^50,51^. Likewise, recent evidence suggests that activation of dopamine neurons in the VTA leads to increased BOLD signals using DREADD-fMRI in rodents^52^. Human fMRI-PET studies have shown that VTA BOLD activation in response to rewards correlates with increases in dopamine in the striatum, suggesting that the fMRI signal in the SN/VTA is related to dopaminergic neuron activity^44,53^. fMRI signals in the cerebellar cortex and VTA may therefore reflect incoming reward-related signals, prediction error processing or, maybe most likely, both.

The origin of the afferent reward signals to the cerebellum remains unclear; while an early study showed afferent connections from the VTA to the cerebellum^54^, this finding has not been replicated in more recent research^55^. However, preliminary evidence from Guarque-Chabrera et al.^56^ suggests the existence of a connection from the VTA to the cerebellum, although its strength and functional relevance remain uncertain. Given this limited evidence, the VTA-to-cerebellum interactions detected with DCM may not be driven primarily by a direct connection. Instead, they may reflect indirect network-level interactions within the mesolimbic and mesocortical systems.

In rodents, a subset of VTA dopamine neurons has been found to be active when the US is expected and does not occur, and are therefore considered to represent (reward-like) prediction error signals driving extinction learning^6,10^. The VTA, however, also contains neurons which are active during the presentation and prediction of threat^57,58^, and likewise we observed fMRI activation of the VTA related to threat prediction and presentation. Cerebellar activations and their interactions are likely not limited to reward-like prediction errors; they may also be related to unexpected presentation of aversive stimuli. Because aversive stimulus presentation results in pronounced cerebellar activations and the response to the US habituates over time, we were unable to separate cerebellar activation related to the unexpected (initial acquisition trials) and the expected (late acquisition trials) presentation of the US.

Although the present findings are consistent with the interpretation of prediction error related activations in the VTA (and cerebellum), based on the present fMRI data, we cannot decide whether activations reflect surprise signals (i.e., unsigned prediction error related signals) or signed prediction error signals^39,59–61^. Rodent data show that cerebellar neurons in the cortex and nuclei increase activations both related to the unexpected presentation and omission of reward signals^62^, which would indicate that the cerebellum primarily contributes to unsigned (reward) prediction errors. Likewise, a meta-analysis of human prediction error showed that activations in the cerebellum were more consistently related to unsigned prediction errors^63^.

Results showed significant functional interaction between the cerebellum and VTA during unexpected US omission in extinction training, and interactions were also observed during spontaneous recovery and reacquisition. This suggests that cerebellum-VTA coupling consistently emerges in conditions where prediction errors occur. Predictive signals of the cerebellum may be linked to the timing of the unconditioned response and therefore its omission and may interact with VTA-related prediction error processing.

Although interpretation must remain cautious given BOLD and DCM limitations outlined in more detail below, we detected phase-specific differences in cerebellum-VTA connectivity. Extinction learning showed bidirectional coupling, spontaneous recovery showed primarily VTA-to-cerebellum influences, and reacquisition showed predominantly cerebellum-to-VTA influences. A tentative interpretation is that a bidirectional loop may support the formation of an extinction memory during active learning, whereas in spontaneous recovery the extinction memory is mainly retrieved, possibly reducing the need for cerebellar output. In reacquisition, reactivation of the fear memory may again engage cerebellar outputs that modulate omission-related processing in the VTA, while the reactivated fear memory during reinforced trials may reduce the detectability of VTA-to-cerebellum influences.

Notably, the unexpected US omission area in the cerebellum was more prominent on the left, while we applied the US to the right. In our previous work it was also the left cerebellum which was mainly activated, however, stimulation was always done on the left^18,64^. As outlined above, we believe that cerebellar activation is reward related. In a recent meta-analysis of fMRI data in monetary reward tasks, reward outcome was also related to the left cerebellum (i.e., lobule VI in the cerebellar hemisphere and vermis)^34^. There is good evidence that emotional information is most prominently processed in the right cerebral hemisphere^65,66^. The present data suggest that emotional processing may also be lateralized in the cerebellum, with the left cerebellar hemisphere projecting to the right cerebral cortex and vice versa^67^. Lateralization of positive emotions to left cerebellar lobule VI has also been reported by Liu et al.^68^. Furthermore, emotion-related tasks resulted in increased activity of the left cerebellar hemisphere in patients with anorexia nervosa compared to controls^69^. Meta-analyses of fMRI studies of emotional tasks, however, report mostly bilateral cerebellar activations with no clear lateralization^70–72^. These studies primarily focused on emotion recognition tasks. Future research should explore the lateralization of emotional processing in the cerebellum using a variety of emotional tasks. In addition to emotions, there is also a preference of the right cerebral hemisphere for the ventral fronto-parietal attention network, which has been related to the detection of potentially harmful events^73^. Thus, as outlined above, attention-related processes involved in fear conditioning may also contribute to (lateralized) cerebellar activations.

In the following, we discuss potential limitations of our study. Firstly, while we demonstrated functional connectivity in the DCM analysis, the results were weaker with PPI. PPI is known to frequently lack power and to have a high proportion of false negatives^74,75^, particularly for event-based designs. Additionally, the low number of unexpected events in our experiment, due to the 100% reinforcement/non-reinforcement rates in acquisition training (which was designed to induce maximum prediction error in the initial extinction trials), may have made detecting activation challenging. Furthermore, as the PPI regressor is demeaned, any persistent activation (e.g., due to an anatomical connection) will decrease detected connectivity. Because cerebellar and VTA activation co-occur not only in response to unexpected US omissions, but also to US presentation and prediction, this lack of specificity could have weakened our results. The same reasons apply for the lack of observation in functional connectivity between the DCN and the VTA.

Secondly, in the DCM analysis, we observed strong modulation related to unexpected US omissions when using the cerebellar cortex and VTA as volumes of interest (VOIs), but observed only trend-level results when substituting the cerebellar cortex with the DCN. This is likely explained due to the high iron content in the DCN causing susceptibility artefacts and reducing signal^76^. Although we opted for a whole-brain sequence with a conventional echo-time in order to generate a dataset of general value, future studies could opt for lower echo times and optimize for detection of signal in the DCN^77^.

Thirdly, the extent of the VTA varies depending on the MRI atlas used. The VTA in the atlas from Pauli et al.^78^ is about 4 times smaller than the VTA in the atlas by Trutti et al.^79^ due to different VTA definitions used. Our average volumes are in good accordance with Trutti et al.^79^, which in turn is in good accordance with histological data. The VTA and its boundaries, particularly laterally, are not clearly visible on MRI scans and were identified by using landmarks such as the substantia nigra, red nucleus, interpeduncular fossa and the cerebral aqueduct. We have approximated the VTA region as closely as we could, but cannot rule out imperfect masks, especially at both lateral ends of the VTA.

In conclusion, our study provides evidence that interactions between the cerebellum and VTA contribute to fear extinction learning. These interactions may support reward-like prediction error processing, a mechanism thought to drive fear extinction learning. Previous studies found that the cerebellum modulates addictive and social behavior via direct projection to VTA in rodents^16,23^. In humans, our findings extend this framework by highlighting cerebellum-VTA interactions involved in emotional processing. Incorporating these interactions into models of the fear extinction network may provide new ways of improving exposure therapy by targeting the dopaminergic system or stimulating the cerebellum specifically during unexpected learning events.

## Methods

### Preregistration

This study was preregistered on OSF on 16/11/2023^80^. The preregistration document can be viewed at https://doi.org/10.17605/OSF.IO/2PXWE.

The preregistration outlines the methods explained here, and includes the main hypotheses. Adjustments were made to specific parts of the methods, which include the optimized localization of the VTA and DCN by manual drawing by an expert annotator, and the inclusion as volumes of interest (VOIs) in the functional connectivity analyses.

### Deviations from preregistration

A primary deviation from the preregistration involved the event-based fMRI analysis of unexpected US omissions. The preregistered analysis plan specified event-based contrasts using early versus late halves of extinction, recall, reacquisition, reextinction and unexpected US phases. In addition, the preregistration described an exploratory analysis focusing on the initial three unexpected omission trials within these phases. In the final analyses, this preregistered exploratory first-three-trial contrast was used as the main event-based fMRI analysis, replacing the early/late block contrasts. This decision was supported by the rapid decay of behavioral differentiation and model-derived prediction error estimates observed under the full (non-)reinforcement structure of the paradigm (100% reinforcement during acquisition and 0% during extinction and recall), which indicated that prediction errors were largely confined to the initial trials (see Figure 2 for SCRs, Figure 3 for PSRs, and Figure 9 for prediction error estimates).

Furthermore, several preregistered exploratory analyses were not performed. In particular, a preregistered exploratory multi-node DCM of the broader fear extinction network was omitted as it was out of scope for the current VOI-specific manuscript centered on cerebellum-VTA interactions. Other preregistered exploratory analyses were likewise not performed, as they were not necessary to address the primary hypotheses of the present study.

In addition, the preregistration contains a reporting error in the sample size rationale: the alpha level was listed as *p* < 0.05, whereas the underlying power analysis conducted during grant preparation used *p* < 0.005. This affects the estimated minimum sample size but not the target sample size, which was 50 participants in both the preregistration and the original manuscript. The correct parameters are reported below.

### Participants

#### Experiment power estimate and number of participants

Experiment power was estimated based on previous fear conditioning data acquired at 7T fMRI^18^ using the FMRIPower toolbox (fmripower.org)^81^. Considering first level unexpected US omission contrasts during a partially reinforced acquisition training and aiming for a power of 80% at *p* < 0.005 for a one-sided hypothesis test, group sizes were estimated at 41 participants for Crus I ipsilateral to US presentation (effect size 0.56 in units of standard deviations). To account for potential dropouts and outliers, 50 participants were initially recruited. However, only 44 successfully completed the 3-day study, with reasons for non-completion including Covid-related issues (1 participant), problems with the Digitimer equipment (3 participants), and failure to attend on day 3 (2 participants). Additionally, one participant dropped out due to a shimming failure, resulting in a final sample size of 43 participants.

#### Exclusion criteria and demographic information

Participants were between 20 and 35 years old (23 men, 20 women, mean age: 24.7 (SD = 3.4) years), fluent in German, non-smokers, reported no intake of medication or illicit drugs affecting the central nervous system, had no history of neurological or mental disorders for themselves or their first-degree relatives, and had no previous participation in similar learning experiments. Additionally, female participants were not pregnant, breastfeeding, or using hormonal contraceptives. Before the experiment, participants completed the Depression Anxiety Stress Scale (DASS21G)^82^. Participants who scored higher than 20 for depression, 14 for anxiety or 25 for stress components were excluded from the study. All participants were right-handed based on the Edinburgh handedness inventory^83^. Participants were asked to refrain from alcohol consumption the night before the experiment. Informed consent was obtained from all participants. The study was approved by the local ethics committee and conducted in accordance with the Declaration of Helsinki.

#### Fear conditioning paradigm

Participants underwent a three-day differential fear conditioning paradigm. The paradigm presentation was controlled by a computer running Presentation (version 21.0, Neurobehavioral System Inc, Berkeley, CA). Participants were shown two visual stimuli (square and diamond) as CS (CS+ and CS-) and informed that electrical stimulations would be applied during the experiment. Three types of trials were used: reinforced CS+ (CS+ paired with the US; CS+/US), unreinforced CS+ (no US) and unreinforced CS-. The CS- was never reinforced.

The paradigm consisted of seven phases separated over three days (Figure 1A). On day 1, the experiment started with a habituation phase consisting of 8 unreinforced CS- and CS+ trials (4 CS+ trials, 4 CS- trials, presented in semi-randomized order). This phase is intended to decrease novelty effects and enable participants to acclimate to the scanner environment and presentation of visual stimuli. The habituation phase was followed by acquisition training consisting of 32 trials with fully reinforced CS+ (16 CS+/US, 16 CS- trials), during which the fear association was learned. On day 2, extinction training was applied and consisted of 32 unreinforced trials (16 CS+, 16 CS-). On day 3, the recall test, reacquisition, reextinction and the unexpected US phase were applied. The recall test consisted of 16 unreinforced trials (8 CS+, 8 CS-). This phase tested the recall of extinguished fear responses and showed spontaneous recovery (see Results). Reacquisition, reextinction, and the unexpected US phases were done in a single continuous fMRI run without pauses between sequences. The partially reinforced reacquisition phases consisted of 36 trials (15 CS+/US trials, 3 CS+, 18 CS-) with 3 unexpected omissions of the US, reextinction consisted of 24 trials (12 CS+, 12 CS-) and the unexpected US phase consisted of 28 trials (3 CS+/US trials, 11 CS+, 14 CS-) with 3 unexpected reinforced CS+ trials (Figure 1A). Learned fear responses were measured by skin conductance responses (SCRs) and pupil size reactions (PSRs) in each trial. The reacquisition, reextinction and unexpected US phases were included to increase the number of trials with high prediction errors (i.e., discrepancy between US expectation and actual US presentation).

We used the model from Batsikadze et al.^30^ to optimize our experimental design. Values for US prediction and US prediction error produced by the model were recorded during simulation. Simulations were repeated for different paradigm CS sequences, and the sequence which required a short number of trials and still yielded high prediction errors was selected.

The CS+ and CS- were presented in equal numbers during the first and second half of each phase, regardless of whether the CS+ was followed by a US or not. Counterbalancing was implemented for the visual CS (CS+ square and CS- diamond, or CS+ diamond and CS- square) as well as for the order of presentation of the CS during the unreinforced habituation, extinction training, and recall test phases (i.e., CS+ presented first or CS- presented first). The reinforced acquisition training and reacquisition phases always started with a CS+. The two orders of CS presentation were pseudorandomly generated, similar to Ernst et al.^18^. Participants were instructed to pay attention to any possible connection between the CS and US presentations. They were also informed that the patterns of CS and US presentations would stay the same across all phases. However, they were not told about the CS/US contingencies or the timing and occurrence of the US.

In the analysis, the focus was on trials containing unexpected omissions of the US (CS+/no US) with high prediction error values. These trials occurred during early extinction, early recall, reacquisition and the reextinction phase.

#### Visual conditioned stimuli (CS)

Visual stimuli were displayed on an fMRI monitor (BOLD-screen 32, Cambridge Research Systems Ltd., Rochester, UK) placed at the end of the scanner bore which was projected to the participant using a mirror system (Figure 1B). Two pictures of black geometric figures (a square and a diamond shape) on a gray background were used as CS+ and CS-. Both CS were presented for 6 s. The period of the CS was shortened from 8 s used in our previous study^18^ to 6 s to decrease the length of the experiment, while still being able to assess SCRs^4^. In the periods between visual stimuli a fixation cross was shown (time of ITI: varies between 9 s and 13.5 s).

#### Aversive unconditioned stimulus (US)

In reinforced CS+ trials, the visual stimulus co-terminated with the presentation of the aversive US, an electrical stimulation (100 ms) applied to the first dorsal interosseous muscle area of the right hand via a 6.5 mm concentric surface electrode (WASP electrode, Specialty Developments, Bexley, UK). The US was produced by a high voltage DC stimulator (DS7A, Digitimer Ltd., London, UK) and consisted of four consecutive 500 µs current pulses (maximum output voltage: 400 V) with an interpulse interval of 33 ms. The location of the US electrode was kept constant across all days of measurement by marking the electrode position with a permanent marker on day 1 and 2. Before the first experimental phase began, the individual electrical stimulation intensity threshold was calibrated to be perceived as ‘very unpleasant but not painful’. To reach the threshold, the current strength was progressively increased and modulated according to each participant’s feedback. The calibrated US intensity was increased by 20 % to compensate for possible habituation to the US, which could result in a weakening of the conditioned responses as successfully done in previous studies of our group^30,84^. The US intensity remained the same for the three days of the experiment (mean US intensity: 3.8 (SD = 2.9) mA, range 0.6-19.2 mA).

#### Physiological data acquisition

During each phase, SCRs and pupil size were recorded. SCRs were acquired with appropriate hardware filters sampling at 2 kHz through an MP160 Data Acquisition Hardware unit (BIOPAC Systems Inc, Goleta, CA). Two SC electrodes were attached to the left-hand hypothenar eminence. Pupil size was measured using an MRI compatible eye tracking system (Avotec Inc, Stuart, FL). Calibration of the eye tracking system was performed prior to each phase to track gaze position on the screen.

#### Self-reports

After each learning phase, participants completed questionnaires using a 4-button fiberoptic response device (Current Designs, Haverford, PA). The Likert scale questionnaires assessed self-reports such as arousal (rated on a 1-9 scale from 1: “very calm” to 9: “very nervous”), valence (rated on a 1-9 scale from 1: “comfortable” to 9: “uncomfortable”), and fear (rated on a 1-9 scale from 1: “not afraid” to 9: “very afraid”) related to the visual stimuli. The questionnaires also included questions about US perception and CS-US contingency. Additionally, participants completed standardized questionnaires before the start of the experiment to assess handedness, depression, anxiety, and stress levels. All participants were screened for contraindications of 7T MRI.

#### Skin conductance analysis

To eliminate high frequency artefacts, skin conductance data was first low-pass filtered at 10 Hz (62^nd-^order Blackman FIR filter) in MATLAB (Release 2022b, RRID:SCR_001622, The MathWorks Inc., Natick, MA). SCRs were defined as the maximum trough-to peak amplitude within a given time interval after CS onset using semi-automated peak detection implemented in a MATLAB-based EDA-Analysis App^85^. A minimum amplitude criterion of 0.01 μS was used as the SCR detection threshold. In each trial, SCRs were evaluated for two distinct time windows: the conditioned response within a time window of 1.0 s to 5.9 s after CS onset and the unconditioned response within a time window of 6 s to 10 s after CS onset (irrespective of whether a US was presented). To account for between-subject variance, the resulting raw SCR amplitudes were increased by 1 μS and normalized through a logarithmic transformation (LN(1+SCR))^86^.

#### Pupil size analysis

Preprocessing of the raw pupil size data was performed to detect and remove blinks^87^. Trials with fewer than 50% of their data points remaining after blink removal were excluded from the analysis. For each trial, the baseline was computed as the mean pupil size during the 500 ms period prior to CS onset. The baseline was subtracted from the corresponding pupil size^88,89^ and the result was divided by the baseline to compute the pupil size response. The mean pupil size response for CS+ and CS- trials were calculated in the time windows from 4 to 5.9 s and 6.0 to 8.0 s for the CS and US pupil size response, respectively. Pupil size responses during possible US presentation were excluded (5.9 to 6 s). The time window for the CS was chosen based on Jentsch et al.^88^ who observed the largest differentiation between CS+ and CS- 2 s prior to the US. The US time window was chosen to match the CS time window. For valid trials, a missing pupillometric data point was considered as ‘not a number value’ to not influence the pupil size mean computation^89^.

#### Behavioral statistical analysis

For physiological data, due to non-normal distribution (SCR and PSR: Shapiro-Wilk test, *p* < 0.001), statistical analysis was performed using a non-parametric ANOVA-type statistic (ATS) implemented via the ANOVAF option in the PROC Mixed procedure in SAS Studio 3.8 (SAS Institute Inc., Cary, NC, USA). This method is suggested for addressing skewed distributions, outliers, or small sample sizes. To ensure more reliable results, an ATS was applied, with the denominator degrees of freedom set to infinity^90–92^ as the use of finite denominator degrees of freedom can result in increased type I errors^93^. The ATS was calculated for each phase (habituation, acquisition training, extinction training, recall test, reacquisition, reextinction, unexpected US phase), with stimulus type (CS+, CS-) and block (early vs. late, defined as the first and second halves of the phase, respectively) as within-subjects factors and SCRs or PSRs as dependent variables.

For analyses of self-reports, an ATS with repeated measures was calculated with stimulus type and phase as within-subjects factors and ratings (valence, arousal, fear and US expectancy) as dependent variables.

For behavioral results, a *p <* 0.05 criterion was used to determine if results were significantly different from those expected if the null hypotheses were correct. Post hoc comparisons were calculated using least square means tests and were adjusted for multiple comparisons using the Tukey-Kramer method.

#### Computational modeling

An artificial agent was trained to predict the likelihood of a US for a given visual input in a virtual version of the experiment as in Batsikadze et al.^30^. The model was based on reinforcement learning^94^ and consisted of a deep neural network (DNN)^95^. The model hyper-parameters were fit to SCRs recorded in the experiment, which served as a proxy for participants’ expectation of an US.

The same simplified visual stimuli for trial stimuli 𝑠_𝑡_ and the same encoding for reinforcement signals 𝑟_𝑡_ were used as in Batsikadze et al.^30^. We employed the same network architecture, which comprised two hidden fully connected layers with 64 units each and an output layer with activation function 𝜑. To account for the counterbalanced order of CS+ and CS-presentations in the experimental design, two trial sequences were modeled separately: In trial sequence 1, habituation, extinction, and recall began with a CS+ trial, whereas in trial sequence 2 these phases began with a CS-- trial. For each of the two trial sequences experienced by the participants, 25 randomly initialized agents were trained at each point of the hyper-parameter grid search. Their outputs were averaged and compared to the group-averaged SCR learning curves, and the final set of hyper-parameters were selected by minimizing a cost function that quantified the difference between model predictions and SCR data. After selecting the best-fitting hyper-parameters, 100 randomly initialized agents were trained anew, and their averaged outputs provided the trial-by-trial prediction 𝑣_𝑡_ and prediction errors 𝛿_𝑡_ used in the fMRI analyses.

On each trial 𝑡, the agent stored an experience tuple 𝑒_𝑡_ = (𝑠_𝑡_, 𝑟_𝑡_, 𝛿_𝑡_) in memory for later replay^96^. The agents were trained using the backpropagation algorithm^97^ on batches of experiences of size b, which were sampled randomly from memory with a probability that was proportional to a priority score 𝑝:

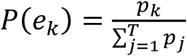

Priority scores depended on the experiences’ recency 𝜆^𝜏^, where 𝜏 is the time passed since the experience and 𝜆 is a decay factor. Optionally, the priority could additionally depend on the magnitude of US prediction error, i.e., 𝑝 = |𝛿|𝜆^𝜏^ ^98^. The parameter RPE ∊ {Yes, No} indicates whether the replay priorities also depended on the magnitude of the US prediction error. The number of replays 𝑖 was varied to control the degree of learning on each trial.

While the previous model could account for ABA renewal^30^, it cannot account for spontaneous recovery in the AAA paradigm, because, in the model, extinction in the same context overwrites the association formed during acquisition. Hence, we extended the model by an additional replay phase, which takes place between habituation, acquisition, extinction, recall and reacquisition phases, and serves to recover the initially acquired association. This replay phase prioritized experiences according to the reinforcement 𝑟_𝑘_ received in a trial and consisted of a total 100 replays of batches of size 128. Reactivation probabilities for these replays were computed as follows:

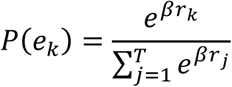

where 𝛽 denotes the inverse temperature and controls the relative difference of reactivation probabilities.

#### Hyper-parameter fitting

We averaged SCRs from CS+ and CS- trials separately for each trial sequence and applied min-max scaling. The averaged SCRs were accordingly defined as 𝑌̅_𝑙_ = (𝑦̅_+,1_, . . ., 𝑦̅_+,𝑁_, 𝑦̅_−,1_, . . ., 𝑦̅_−,𝑁_), where 𝑦̅_+,𝑛_ and 𝑦̅_−,𝑛_ are the averaged SCRs for the n-th CS+ and n-th CS- presentations across all participants who completed a given trial sequence 𝑙, respectively. Analogously, the averaged US predictions of the model were defined as 𝑉̅_𝑙_(𝑏, 𝜆, 𝑖, 𝛽, 𝑅𝑃𝐸, 𝜑) = (𝑣̅_+,1_, . . ., 𝑣̅_+,𝑁_, 𝑣̅_−,1_, . . ., 𝑣̅_−,𝑁_), where 𝑣̅_+,𝑛_ and 𝑣̅_−,𝑛_ are the averaged US predictions for the n-th CS+ and CS- presentations across all model instances who were trained on a given trial sequence 𝑙, respectively. For each trial sequence, 25 randomly initialized agents were trained with the same hyper-parameters, and their outputs were averaged to reduce variability and obtain stable trial-wise estimates of US predictions which are able to show a good fit with the population data and prediction errors. The goodness of fit was defined as:

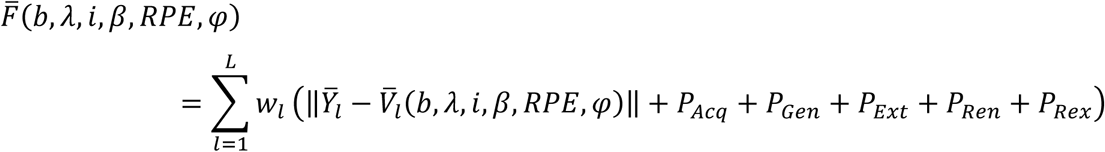

where 𝑤_𝑙_ is the number of participants who experienced trial sequence 𝑙. To ensure that the overall learning curve in the model resembled that of the participants, we added the following penalty terms, if the model failed to

1. learn that CS+ is followed by the US: 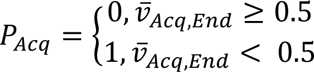 where 𝑣̅_𝐴𝑐𝑞,𝐸𝑛𝑑_ is the average US prediction over the last 2 CS+ presentations of acquisition training.
2. show the CR at the start of extinction training: 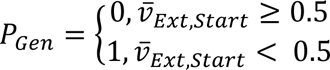 where 𝑣̅_𝐸𝑥𝑡,𝑆𝑡𝑎𝑟𝑡_ is the average US prediction over the first 2 CS+ presentations of extinction training.
3. successfully extinguish the CR during extinction training: 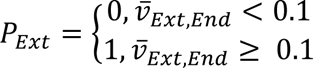 where 𝑣̅_𝐸𝑥𝑡,𝐸𝑛𝑑_ is the average US prediction over the last 2 CS+ presentations of extinction training.
4. show renewal of the CR: 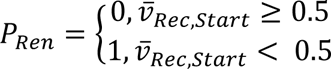 where 𝑣̅_𝑅𝑒𝑐,𝑆𝑡𝑎𝑟𝑡_ is the average US prediction over the first 2 CS+ presentations of recall.
5. successfully extinguish the CR during recall: 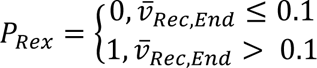 where 𝑣̅_𝑅𝑒𝑐,𝑆𝑡𝑎𝑟𝑡_ is the average US prediction over the last 2 CS+ presentations of recall

A grid search was conducted over the hyper-parameter sets shown in Table 1. The model with the best goodness-of-fit was then chosen to derive trial-by-trial values for US predictions and prediction errors. Prediction errors were used for parametric modulation of fMRI data to test hypotheses related to prediction error processing of the cerebellum and VTA.

**Table 1:**
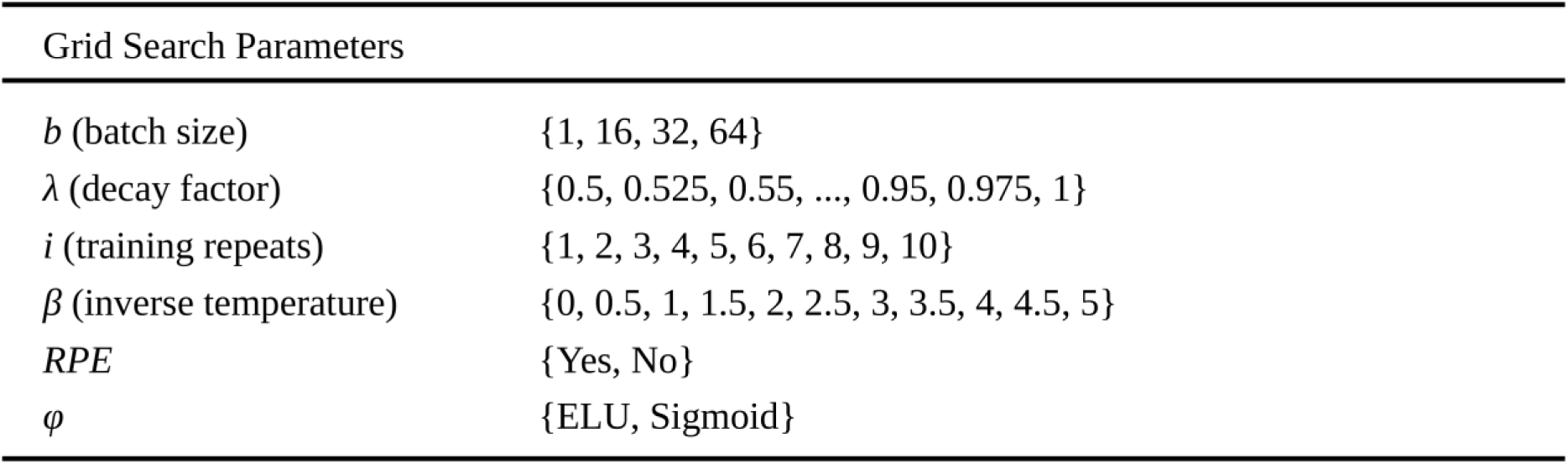
Modeling. Parameters the grid search was run for.

#### MRI acquisition

All MRI imaging data were collected while participants were lying supine in the 7T MRI scanner (MAGNETOM Terra, Siemens Healthineers AG, Forchheim, Germany). A 1-channel transmit/32-channel receive head RF coil (Nova Medical Inc., Wilmington MA, USA), was used.

Two dielectric pads for signal homogenization were placed on either side of each participant’s upper neck^99^. Depending on head size, further cushioning was added to prevent participant head movement and discomfort.

Whole-brain functional MRI acquisition was performed with a 3-dimensional echo planar image sequence^100^ with an isotropic voxel size of 1.5 mm. The sequence was run for each phase (habituation, 98 volumes; acquisition training, 354 volumes; extinction training, 350 volumes; recall test, 182 volumes; reacquisition, reextinction and unexpected US phases; 938 volumes). Imaging parameters were as follows: TR/TE, 1620/20 ms; flip angle, 11°; phase encoding acceleration factor, 2; 3D acceleration factor, 3; slice partial Fourier factor, 7/8; acquisition matrix, 140 × 140; number of slices, 96.

After acquisition training on day 1, a transversal QSM ASPIRE sequence was acquired^101,102^ with an isotropic voxel size of 0.7 mm. Further imaging parameters were as follows: TR/TE1/TE2/TE3/TE4/TE5, 28/5/10/15/20/25 ms; flip angle 15°; phase encoding acceleration factor, 3; slice partial Fourier factor, 6/8; acquisition matrix, 320 × 280; number of slices, 256; TA, 10:52 min.

After the acquisition of the QSM ASPIRE sequence, a sagittal MP2RAGE sequence including fat navigators^103–105^ was acquired with an isotropic voxel size of 0.75 mm. Further imaging parameters were as follows: TR/TE, 6000/1.85 ms; TI1/TI2, 800/2750 ms; flip angles 1/2, 4°/5°; phase encoding acceleration factor, 3; acquisition matrix, 340 × 340; number of slices, 256; TA, 14:58 min.

#### Image processing

Motion correction of MP2RAGE volumes including fat navigators was performed using offline reconstruction with the Retro-MoCo toolbox for MATLAB provided by David Gallichan (version23 0.9.0dev, https://github.com/dgallichan/retroMoCoBox.git). A T1 map was generated from the motion corrected MP2RAGE with the MP2RAGE-utils package implemented in MATLAB (release 1.0, https://github.com/srikash/MP2RAGE-utils). The MP2RAGE was normalized to MNI-space using the CAT12 (CAT12, release 1450)^106^ toolbox in SPM12 (Wellcome Department of Cognitive Neurology, London, UK).

Functional MRI volumes were brain extracted with BET (Brain Extraction Tool)^107^ in FSL (Release 6.0.1, RRID:SCR_002823, Centre for Functional MRI of the Brain, Oxford, UK). Motion and distortion correction was performed for each fMRI run with ANTs (Version 2.3.5, RRID:SCR_004757, University of Pennsylvania, Philadelphia, USA)^108^. A 5-volume fMRI sequence preceding each run with phase encoding in the opposite direction was used for the distortion correction. All volumes were coregistered to a T1 map derived from the acquired MP2RAGE with ANTs. The CAT12 processing of the MP2RAGE was used to normalize fMRI data. A 4.5 mm Gaussian kernel was used for smoothing. Motion nuisance regressors (3 translations, 3 rotations) were derived from ANTS affine transformation matrices output.

#### fMRI analysis

The 1st level analysis was done in MNI space, defining conditions for both the CS and US window (i.e., CS+, CS-, US post CS+, no US post CS+, no US post CS-) separately for each phase. Regressors were calculated from the conditions in SPM12 by convolving a canonical hemodynamic response function with events, which were modelled as delta functions in an event-based design (i.e., the duration of each event is 0 s). Specifically, CS events were time-locked to CS onset (appearance of the visual stimulus), whereas US related events were time-locked to CS offset (6 s after CS onset), which coincided with either delivery or omission of the US. This event-related separation allowed us to test CS related contrasts at CS onset, including CS differentiation (CS+ > CS-). US-related contrasts were tested at CS offset and included contrasts reflecting unexpected omission of the US (e.g. first 3 no US post CS+ > no US post CS-) and US presentation (US post CS+ > no US post CS-). Beta weights were fitted for each regressor by a general linear model for each voxel fMRI time series. Contrasts were computed as linear combinations of beta weights. Values in contrast maps were converted to t-statistic values in first level analysis. Contrast maps are combined and used in a one-sample t-test for second level analysis in SPM12. Tests for significance (p < 0.05) were done after threshold-free cluster enhancement (TFCE toolbox in SPM12, R174, http://dbm.neuro.uni-jena.de/tfce/) and family-wise error (FWE) corrections. Any trend-level results refer to significance tests without TFCE and FWE corrections (p < 0.05). For the cerebellar cortex, results were visualized on cerebellar flatmaps using the SUIT toolbox in SPM12 ^22^. For the cerebellar nuclei and VTA, results were visualized in coronal slices and 3D renderings generated in MRIcroGL^109^ using custom probabilistic atlases (described in “Volume of interest definition”). To acquire cerebellar anatomical region labels, activation maps were projected onto the SUIT atlas volume (Cerebellum-SUIT.nii, Diedrichsen, 2006).

#### Event-based analysis

In addition to main effects of conditions (e.g., during acquisition training), events were additionally separated over time for each experimental phase. For fear acquisition training, extinction training, recall test, reacquisition, reextinction and the unexpected US phase, events were grouped in two equal-size blocks representing early and late halves of those phases (e.g., the 8 first CS+ trials of fear acquisition training correspond to early acquisition and the 8 last CS+ trials of fear acquisition training correspond to late acquisition), with equal numbers of CS+ and CS− trials in each block. Trials with unexpected events—such as unexpected US presentations or omissions—were treated as separate single trial regressors. These included the initial three trials of acquisition training, extinction training, recall test, reacquisition and reextinction and specific events such as the three unexpected US omissions in reacquisition. When CS+ trials were modeled as single trials in the GLM, CS- trials were modeled in the same way to maintain a consistent model structure. This modeling choice did not translate into paired or numerically matched CS+ and CS− trials being used in the event-based contrasts. Rather, event-based contrasts for unexpected US omissions contrasted the first three unexpected omissions following CS+ against all “expected” omissions following CS-(e.g., [first 3 no US post CS+ > no US post CS-]).

#### Parametric modulation analysis

Parametric modulation of fMRI data was performed with prediction and prediction error values in all learning phases derived from our computational model. CS+ and CS- events were modulated with prediction values, while US omission and presentation events were modulated with prediction error values (Figure 9). Events were not further divided within phases (e.g., no early or late halves or separation of the first three trials). Modulations were done separately for each phase, except for reacquisition, reextinction and the unexpected US phase which were modulated together as they were part of the same fMRI run.

#### Volumes of interest (VOI) definition

A global conjunction between unexpected US omission parametric modulation contrasts during extinction, the recall test and reacquisition was multiplied with a cerebellum mask and showed a region in lobule VI and Crus I. This region was used as a cerebellar cortex (lobule VI and Crus I; Figure S4) volume of interest (VOI). Deep cerebellar nuclei (DCN) and VTA VOIs were drawn for each participant. Drawing of the DCN (i.e., the left and right dentate, globose, emboliform and fastigial nuclei) was done by an expert annotator using both MP2RAGE and QSM information in ITK-SNAP (version 3.8.0)^111^. The DCN were combined into a single bihemispheric VOI. Drawing of the VTA was done by adjusting estimated masks with both MP2RAGE and QSM information, using the red nuclei, substantia nigra, the cerebral aqueduct and interpeduncular fossa as landmarks. Initial masks were estimated for the VTA by inversely transforming the probabilistic atlas by Trutti et al.^79^ thresholded at a probability of p > 0.4. All drawn masks were transformed into MNI space. After transformation to MNI space, a probabilistic atlas was generated by averaging binary masks for each structure across participants. The probabilistic atlas was thresholded so that the volume of each structure matched the average volume across participants. For drawing of globose, emboliform and fastigial nuclei, the thresholded atlas was inversely transformed from MNI to T1 space in order to add missing nuclei for each participant with lower quality QSM data. The VTA VOI was used as a seed region in the PPI analysis. In addition, the cerebellar cortex (CB), DCN and VTA VOIs were used as nodes in the 2-node dynamic causal modelling (DCM) networks. The DCN and VTA VOIs were used as masks for displaying fMRI activations in coronal slices and 3D renderings.

#### Functional connectivity: PPI

For the PPI analysis, the VTA VOI was used to extract time series with SPM12 in MNI space. Time series were used as physiological regressors, and psychophysiological regressors were added for each condition using the gPPI toolbox^112^. Both first- and second-level analyses were performed with the same contrasts as defined in the parametric modulation (e.g., no US x prediction error during extinction) analysis. To summarize connectivity in the parametric modulation analysis, all contrasts related to unexpected US omissions were combined in a single contrast within the gPPI toolbox.

#### Functional connectivity: DCM

To calculate dynamic causal models (DCMs), the cerebellar cortex (CB) and VTA VOIs were used to extract time series in a concatenated version of the event-based and parametric modulation models using SPM12^113–115^. Each condition was defined as input to the CB and VTA nodes (i.e., CS+, CS-, US post CS+, no US post CS+, no US post CS-), while each prediction or prediction error related event-based contrast (i.e., CS+ > CS-, first 3 no US post CS+ > no US post CS-) or parametric modulation (i.e., no US prediction error) could modulate the reciprocal intrinsic connections. Recurrent connections were active but were not modulated. Parametric Empirical Bayes (PEB) was used for analysis on the second level^116^. All DCM and PEB analyses were done within SPM12.

## Acknowledgements

We would like to thank our technician Beate Brol for the drawing and adjustment of DCN and VTA masks. Additionally, we would like to thank Greta Wippich for providing helpful illustrations. The MAGNETOM Terra 7T MRI system used in the study was funded by the Deutsche Forschungsgemeinschaft (DFG, German Research Foundation), grant number 432647511. This project has received funding from the DFG (project number 316803389 - SFB1280), the European Union’s Horizon 2020 research and innovation program under the Marie Skłodowska-Curie grant agreement No 956414 and an individual scholarship for Patrick Pais Pereira from the Hans-Böckler-Stiftung.

## Supplementary information

### Skin conductance responses

#### Non-parametric ANOVA SCR results

**Table S1:** Non-parametric ANOVA-type statistics for skin conductance responses (SCRs). Results are shown separately for habituation, fear acquisition training, extinction training, recall, reacquisition, reextinction, and the unexpected US phase. Factors included Stimulus (CS+ vs. CS-), Time (early vs. late halves of each phase), and the Stimulus x Time interaction. Reported statistics include numerator degrees of freedom, F-values, and p-values. Significance levels are indicated as * p < 0.05; ** p < 0.01; *** p < 0.001.

| Factor | Numerator Df | F | p |
| --- | --- | --- | --- |
| *Habituation* |  |  |  |
| Stimulus | 1 | 1.19 | 0.276 |
| Time | 1 | 26.49 | **<0.001\*\*\*** |
| Stimulus x Time | 1 | 0.06 | 0.802 |
| *Fear acquisition training* |  |  |  |
| Stimulus | 1 | 20.79 | **<0.001\*\*\*** |
| Time | 1 | 28.75 | **<0.001\*\*\*** |
| Stimulus x Time | 1 | 3.80 | 0.051 |
| *Extinction training* |  |  |  |
| Stimulus | 1 | 7.71 | **0.006\*\*** |
| Time | 1 | 25.05 | **<0.001\*\*\*** |
| Stimulus x Time | 1 | 3.74 | 0.053 |
| *Recall* |  |  |  |
| Stimulus | 1 | 4.99 | **0.026\*** |
| Time | 1 | 36.74 | **<0.001\*\*\*** |
| Stimulus x Time | 1 | 5.53 | **0.019\*** |
| *Reacquisition* |  |  |  |
| Stimulus | 1 | 27.20 | **<0.001\*\*\*** |
| Time | 1 | 35.44 | **<0.001\*\*\*** |
| Stimulus x Time | 1 | 6.80 | **0.009\*\*** |
| *Reextinction* |  |  |  |
| Stimulus | 1 | 4.47 | **0.035\*** |
| Time | 1 | 0.08 | 0.780 |
| Stimulus x Time | 1 | 0.95 | 0.329 |
| *Unexpected US phase* |  |  |  |
| Stimulus | 1 | 5.43 | **0.020\*** |
| Time | 1 | 0.29 | 0.593 |
| Stimulus x Time | 1 | 2.34 | 0.126 |

#### SCRs for first-three and last-three trial analysis

To assess the robustness of behavioral results when restricting analyses to a small number of trials, we additionally re-binned trials into blocks comprising the first three and last three trials of each phase, instead of halves of each phase. Overall, the results were largely consistent with the primary analysis. The corresponding statistical results are summarized in Table S2.

**Figure S1:**
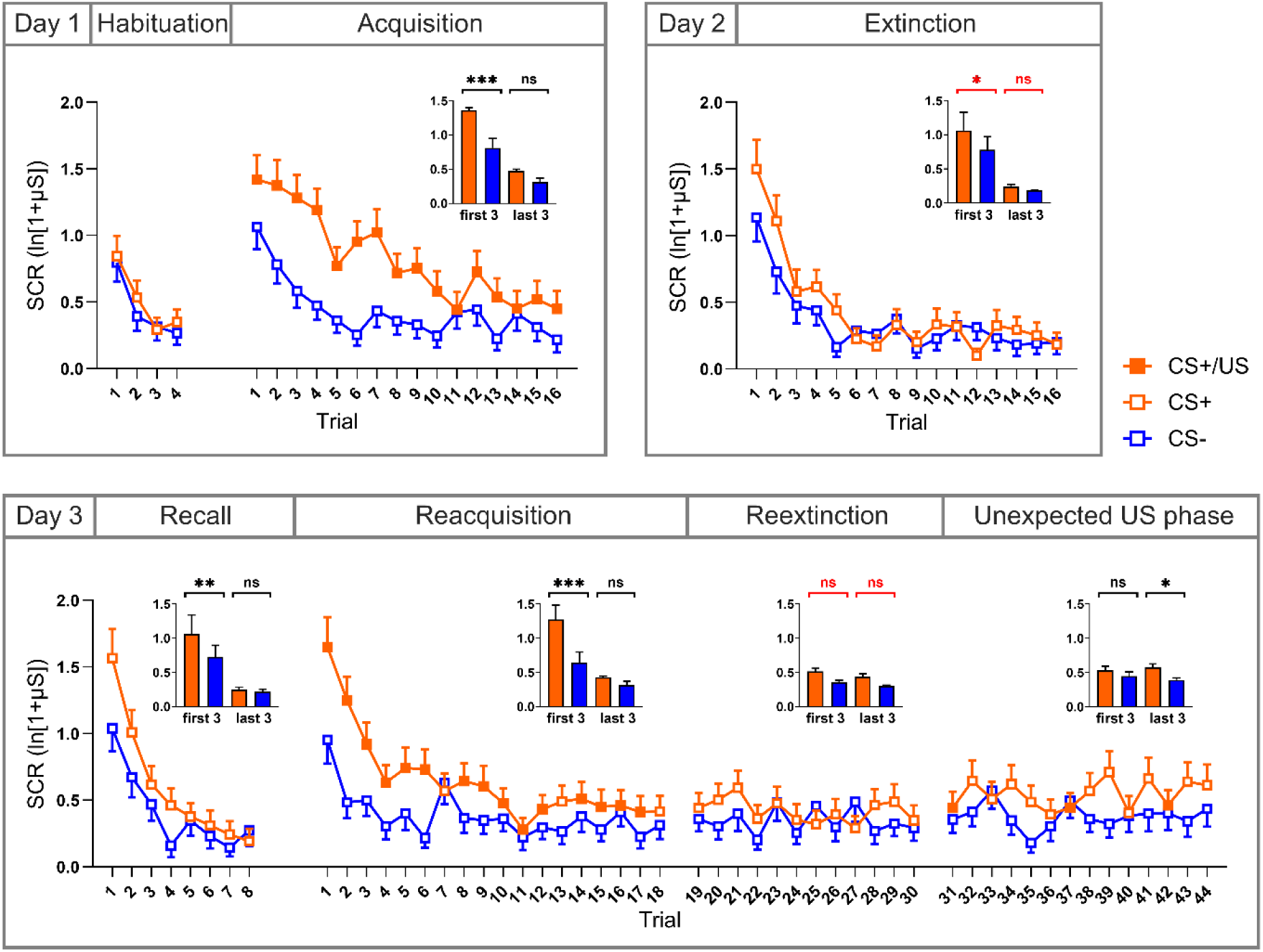
Skin conductance responses (SCRs) for each trial, with CS+ (shown in orange) and CS- (shown in blue) responses paired in blocks. Reinforcement of the CS+ by a US (CS+/US) is indicated by filled squares. The CS- is never reinforced. Bar plots on the top right show mean responses for the first three and last three trials for both CS+ and CS-. On day 1, there was no differentiation between CS+ and CS- in the habituation phase, with significant differentiation emerging during acquisition training. On day 2, CS+/CS- differentiation was confined to the first three extinction trials and was no longer present in the last three extinction trials (trend-level effect; non-significant Stimulus x Time interaction). On day 3, during the initial recall test, participants exhibited spontaneous recovery, i.e., a return of differential responses after extinction training. During initial reacquisition, there were again differential responses to the CS+ and CS-, which decreased in reextinction and the unexpected US phase. Mean values are shown with error bars representing the standard error of the mean. CS: conditioned stimulus; US: unconditioned stimulus; SCR: skin conductance response; μS: microsiemens.

### Non-parametric ANOVA SCR results (first-three and last-three trial analysis)

**Table S2:** Non-parametric ANOVA-type statistics for skin conductance responses (SCRs) based on the first-three and last-three trial analysis. Results are shown separately for habituation, fear acquisition training, extinction training, recall, reacquisition, reextinction, and the unexpected US phase. Factors included Stimulus (CS+ vs. CS-), Time (first three vs. last three trials), and the Stimulus x Time interaction. Significance levels are indicated as * p < 0.05; ** p < 0.01; *** p < 0.001.

| Factor | Numerator Df | F | p |
| --- | --- | --- | --- |
| *Habituation* |  |  |  |
| Stimulus | 1 | 1.19 | 0.276 |
| Time | 1 | 26.49 | **<0.001\*\*\*** |
| Stimulus x Time | 1 | 0.06 | 0.802 |
| *Fear acquisition training* |  |  |  |
| Stimulus | 1 | 8.22 | **0.004\*\*** |
| Time | 1 | 40.06 | **<0.001\*\*\*** |
| Stimulus x Time | 1 | 4.05 | **0.044\*** |
| *Extinction training* |  |  |  |
| Stimulus | 1 | 4.99 | **0.026\*** |
| Time | 1 | 34.62 | **<0.001\*\*\*** |
| Stimulus x Time | 1 | 3.36 | 0.067 |
| *Recall* |  |  |  |
| Stimulus | 1 | 4.24 | **0.039\*** |
| Time | 1 | 47.42 | **<0.001\*\*\*** |
| Stimulus x Time | 1 | 6.35 | **0.012\*** |
| *Reacquisition* |  |  |  |
| Stimulus | 1 | 20.47 | **<0.001\*\*\*** |
| Time | 1 | 50.62 | **<0.001\*\*\*** |
| Stimulus x Time | 1 | 8.70 | **0.003\*\*** |
| *Reextinction* |  |  |  |
| Stimulus | 1 | 9.36 | **0.002\*\*** |
| Time | 1 | 0.02 | 0.895 |
| Stimulus x Time | 1 | 0.47 | 0.494 |
| *Unexpected US phase* |  |  |  |
| Stimulus | 1 | 4.02 | **0.045\*** |
| Time | 1 | 0.83 | 0.363 |
| Stimulus x Time | 1 | 4.67 | **0.031\*** |

### Pupil size responses

#### Non-parametric ANOVA PSR results

**Table S3:** Non-parametric ANOVA-type statistics for pupil size responses (PSRs). Results are shown separately for habituation, fear acquisition training, extinction training, recall, reacquisition, reextinction, and the unexpected US phase. Factors included Stimulus (CS+ vs. CS-), Time (early vs. late halves of each phase), and the Stimulus x Time interaction. Significance levels are indicated as * p < 0.05; ** p < 0.01; *** p < 0.001.

| Factor | Numerator Df | F | p |
| --- | --- | --- | --- |
| *Habituation* |  |  |  |
| Stimulus | 1 | 1.65 | 0.200 |
| Time | 1 | 0.26 | 0.609 |
| Stimulus x Time | 1 | 0.04 | 0.849 |
| *Fear acquisition training* |  |  |  |
| Stimulus | 1 | 39.51 | **<0.001\*\*\*** |
| Time | 1 | 24.31 | **<0.001\*\*\*** |
| Stimulus x Time | 1 | 0.58 | 0.446 |
| *Extinction training* |  |  |  |
| Stimulus | 1 | 1.12 | 0.290 |
| Time | 1 | 18.21 | **<0.001\*\*\*** |
| Stimulus x Time | 1 | 4.28 | **0.038\*** |
| *Recall* |  |  |  |
| Stimulus | 1 | 6.59 | **0.010\*** |
| Time | 1 | 6.41 | **0.011\*** |
| Stimulus x Time | 1 | 4.87 | **0.027\*** |
| *Reacquisition* |  |  |  |
| Stimulus | 1 | 47.87 | **<0.001\*\*\*** |
| Time | 1 | 4.54 | **0.033\*** |
| Stimulus x Time | 1 | 7.70 | **0.006\*** |
| *Reextinction* |  |  |  |
| Stimulus | 1 | 9.10 | **0.003\*** |
| Time | 1 | 0.83 | 0.364 |
| Stimulus x Time | 1 | 4.43 | **0.035\*** |
| *Unexpected US phase* |  |  |  |
| Stimulus | 1 | 17.35 | **<0.001\*\*\*** |
| Time | 1 | 0.83 | 0.363 |
| Stimulus x Time | 1 | 0.03 | 0.872 |

#### PSRs for first-three and last-three trial analysis

To assess the robustness of behavioral results when restricting analyses to a small number of trials, we additionally re-binned trials into blocks comprising the first three and last three trials of each phase, instead of halves of each phase. Overall, the results were largely consistent with the primary analysis. The corresponding statistical results are summarized in Table S4.

**Figure S2:**
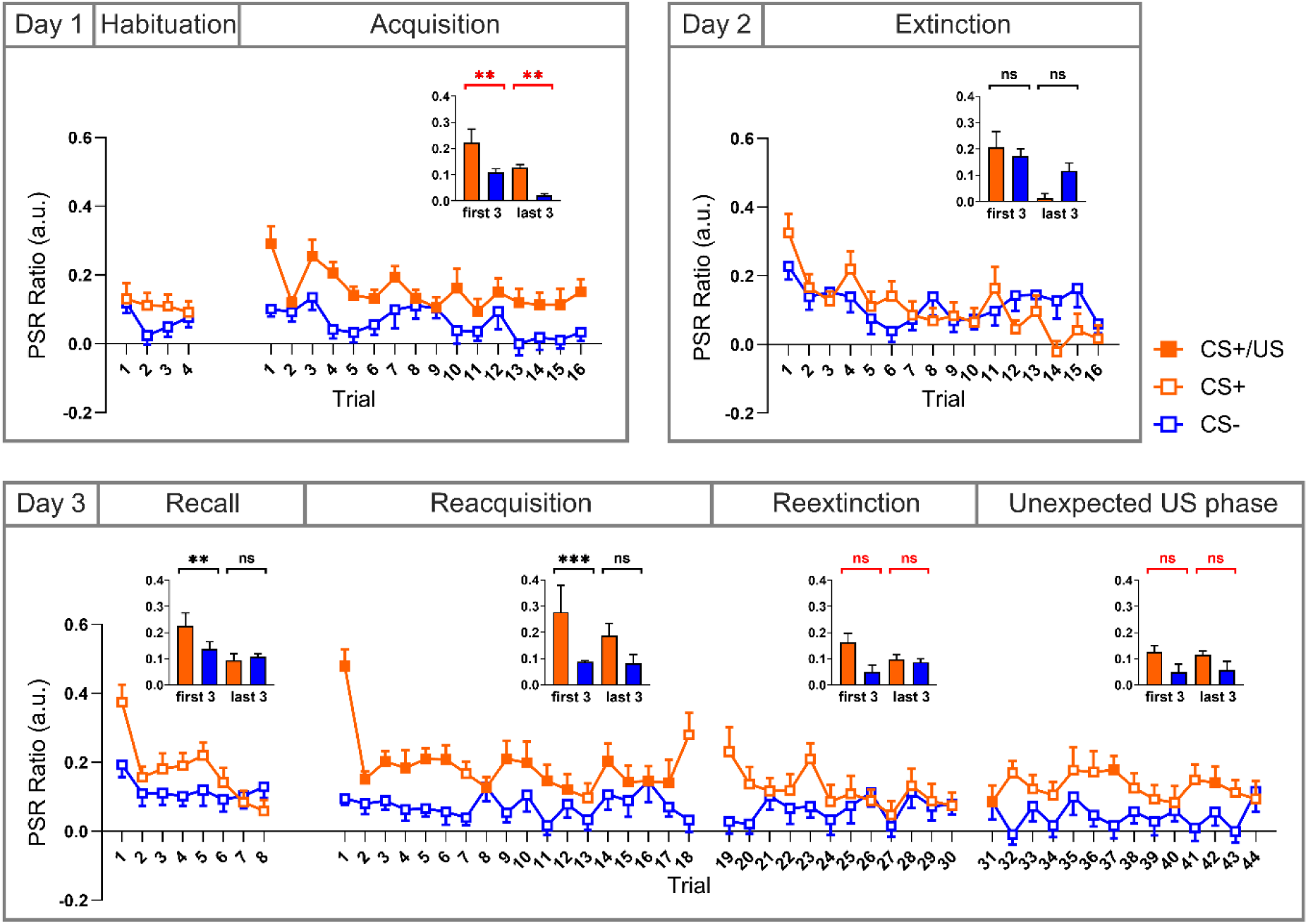
Pupil size responses (PSRs) for each trial, with CS+ (shown in orange) and CS- (shown in blue) responses paired in blocks. Reinforcement of the CS+ by a US (CS+/US) is indicated by filled squares. The CS- is never reinforced. Bar plots on the top right show mean responses for the first three and last three trials for both CS+ and CS-. Significance markers indicate post hoc comparisons between CS+ and CS- within the first three or last three trials; markers are shown in black when the Stimulus x Time interaction was significant and in red when the interaction was not significant. On day 1, there was no differentiation between CS+ and CS- in the habituation phase, with significant differentiation emerging during acquisition training. On day 2, CS+/CS- differentiation decreased across extinction training. Although post hoc CS+/CS- comparisons were not significant during extinction in PSRs, a significant Stimulus x Time interaction was observed, reflecting a decrease from the first three to the last three extinction trials for the CS+ but not for the CS-. On day 3, during initial recall, participants exhibited spontaneous recovery, i.e., a return of differential responses after extinction training. During initial reacquisition, there were again differential responses to the CS+ and CS-, which decreased in reextinction and the unexpected US phase. Mean values are shown with error bars representing the standard error of the mean. CS: conditioned stimulus; US: unconditioned stimulus; PSR: pupil size response.

#### Non-parametric ANOVA PSR results (first-three and last-three trial analysis)

**Table S4:** Non-parametric ANOVA-type statistics for pupil size responses (PSRs) based on the first three and last three trial analysis. Results are shown separately for habituation, fear acquisition training, extinction training, recall, reacquisition, reextinction, and the unexpected US phase. Factors included Stimulus (CS+ vs. CS-), Time (first three vs. last three trials), and the Stimulus x Time interaction. Significance levels are indicated as * p < 0.05; ** p < 0.01; *** p < 0.001.

| Factor | Numerator Df | F | p |
| --- | --- | --- | --- |
| *Habituation* |  |  |  |
| Stimulus | 1 | 1.65 | 0.200 |
| Time | 1 | 0.26 | 0.609 |
| Stimulus x Time | 1 | 0.04 | 0.849 |
| *Fear acquisition training* |  |  |  |
| Stimulus | 1 | 29.02 | **<0.001\*\*\*** |
| Time | 1 | 30.93 | **<0.001\*\*\*** |
| Stimulus x Time | 1 | 0.35 | 0.554 |
| *Extinction training* |  |  |  |
| Stimulus | 1 | 0.93 | 0.334 |
| Time | 1 | 41.57 | **<0.001\*\*\*** |
| Stimulus x Time | 1 | 7.72 | **0.006\*\*** |
| *Recall* |  |  |  |
| Stimulus | 1 | 4.72 | **0.030\*** |
| Time | 1 | 12.23 | **<0.001\*\*\*** |
| Stimulus x Time | 1 | 7.13 | **0.008\*\*** |
| *Reacquisition* |  |  |  |
| Stimulus | 1 | 39.99 | **<0.001\*\*\*** |
| Time | 1 | 9.40 | **0.002\*\*** |
| Stimulus x Time | 1 | 8.22 | **0.004\*\*** |
| *Reextinction* |  |  |  |
| Stimulus | 1 | 4.44 | **0.035\*** |
| Time | 1 | 0.36 | 0.547 |
| Stimulus x Time | 1 | 2.63 | 0.105 |
| *Unexpected US phase* |  |  |  |
| Stimulus | 1 | 4.36 | **0.037\*** |
| Time | 1 | 0.16 | 0.689 |
| Stimulus x Time | 1 | 0.14 | 0.705 |

### Self-reports

#### Self-reports results summary

**Table S5:** Summary of self-report ratings. Median values and interquartile ranges (in parentheses) are shown for arousal, fear, US expectancy, and valence ratings for CS+ and CS- assessed after habituation, acquisition training, extinction training, recall test, and at the end of day 3. Rating scales ranged from 1 to 9, with anchors as indicated for each measure. Interquartile ranges are shown in parentheses. Statistically significant differences between CS+ and CS- are indicated in bold (least squares means tests; p < 0.01).

| Stimulus | Time of assessment |  |  |  |  |
| --- | --- | --- | --- | --- | --- |
|  | Post habituation | Post acquisition | Post extinction | Post recall | End of day 3 |
| *Arousal ratings (1 - very calm, 9 - very excited)* |  |  |  |  |  |
| CS+ | 2 (1-3) | **6 (5-7)** | **3 (1-6)** | **3 (1-6)** | **7 (4-7)** |
| CS- | 2 (1-4) | **2 (1-3)** | **1 (1-3)** | **1 (1-3)** | **1 (1-2)** |
| *Fear ratings (1 - not afraid, 9 - very afraid)* |  |  |  |  |  |
| CS+ | 1 (1-2) | **6 (3-6)** | **2 (1-5)** | **2 (1-4)** | **5 (3-7)** |
| CS- | 1 (1-2) | **1 (1-2)** | **1 (1-2)** | **1 (1-2)** | **1 (1-2)** |
| *US expectancy (1 - US not expected, 5 - unsure, 9 - US expected)* |  |  |  |  |  |
| CS+ | 1 (1-2) | **9 (9-9)** | **3 (1-5)** | **3 (1-5)** | **6 (5-7)** |
| CS- | 1 (1-2) | **1 (1-2)** | **1 (1-1)** | **1 (1-2)** | **1 (1-2)** |
| *Valence ratings (1 - unpleasant, 9 - very pleasant)* |  |  |  |  |  |
| CS+ | 7 (5-8) | **3 (2-4)** | **5 (4-7)** | **6 (5-8)** | **3 (3-5)** |
| CS- | 7 (5-9) | **8 (6-9)** | **7 (5-9)** | **8 (6-9)** | **8 (6-9)** |

#### Non-parametric ANOVA results for self-reports

**Table S6:** Non-parametric ANOVA-type statistics for self-report measures. Results are shown for arousal, fear, valence, and US expectancy ratings, with Stimulus (CS+ vs. CS-) and Time of assessment as within-subject factors, as well as the Stimulus x Time interaction. Degrees of freedom, F values, and p values are reported for each effect. Significance levels are indicated as * p < 0.05; ** p < 0.01; *** p < 0.001.

| Factor | Df | F | p |
| --- | --- | --- | --- |
| *Arousal* |  |  |  |
| Stimulus | 1 | 128.71 | <0.001*** |
| Time | 3.51 | 16.31 | <0.001*** |
| Stimulus x Time | 3.29 | 35.06 | <0.001*** |
| *Fear* |  |  |  |
| Stimulus | 1 | 105.13 | <0.001*** |
| Time | 3.32 | 18.60 | <0.001*** |
| Stimulus x Time | 3.4 | 34.47 | <0.001*** |
| *Valence* |  |  |  |
| Stimulus | 1 | 124.79 | <0.001*** |
| Time | 3.39 | 19.80 | <0.001*** |
| Stimulus x Time | 3.11 | 51.42 | <0.001*** |
| *US expectancy* |  |  |  |
| Stimulus | 1 | 451.62 | <0.001*** |
| Time | 2.4 | 62.78 | <0.001*** |
| Stimulus x Time | 2.8 | 187.33 | <0.001*** |

### fMRI results

#### fMRI PPI results: VTA connectivity with a cerebellar (CB) seed

**Figure S3:**
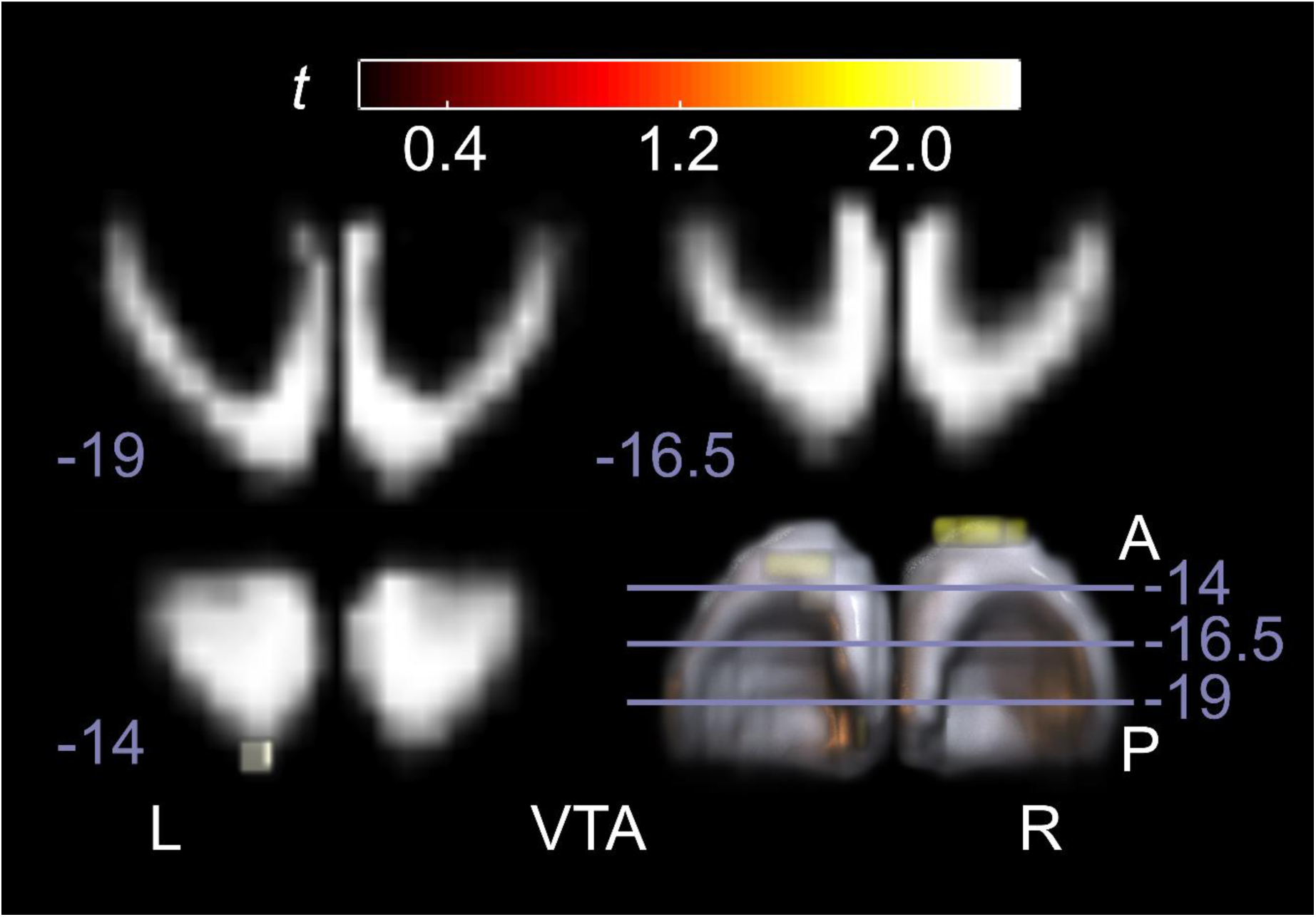
Trend-level psychophysiological interaction (PPI) results using a cerebellar cortex seed (CB; VOI defined from the conjunction analysis; see VOI definition in Methods) during unexpected US omission contrasts. Cerebellar activations are displayed on cerebellar flatmaps (SUIT), while midbrain activations are shown on coronal slices progressing from posterior to anterior, with MNI y-coordinates indicated for each slice. The color scale corresponds to uncorrected voxelwise t-values (p < 0.05, uncorrected). Effects in the ventral tegmental area (VTA) are weak and spatially limited, reflecting the reduced sensitivity of cerebellar-seed PPI analyses in the present dataset. This analysis is included for completeness and illustrates connectivity patterns when reversing the seed direction relative to the main VTA-seed-PPI analyses shown in the main figures.

#### Summary statistics for VOI contrast estimates across fMRI contrasts

**Table S7:**
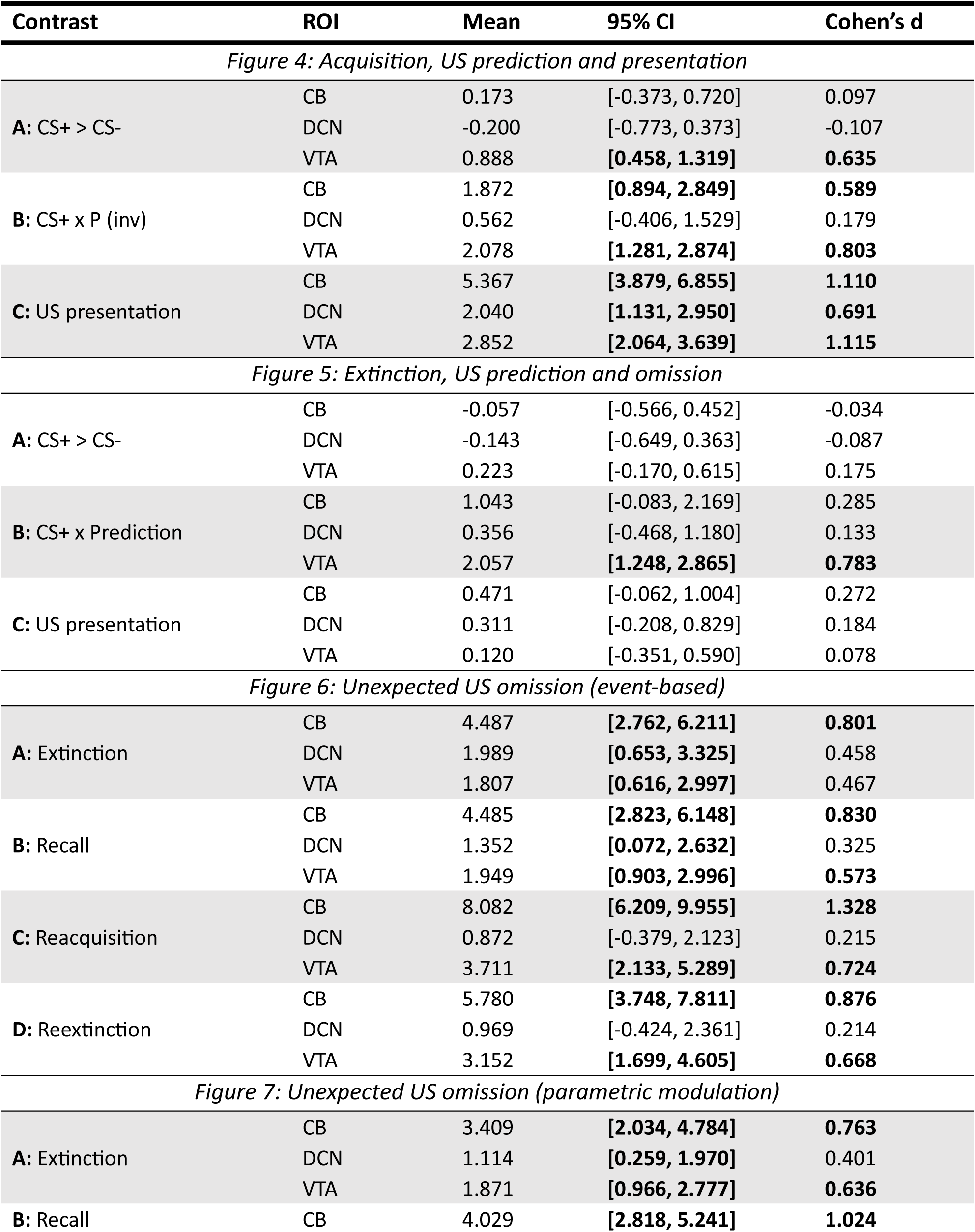
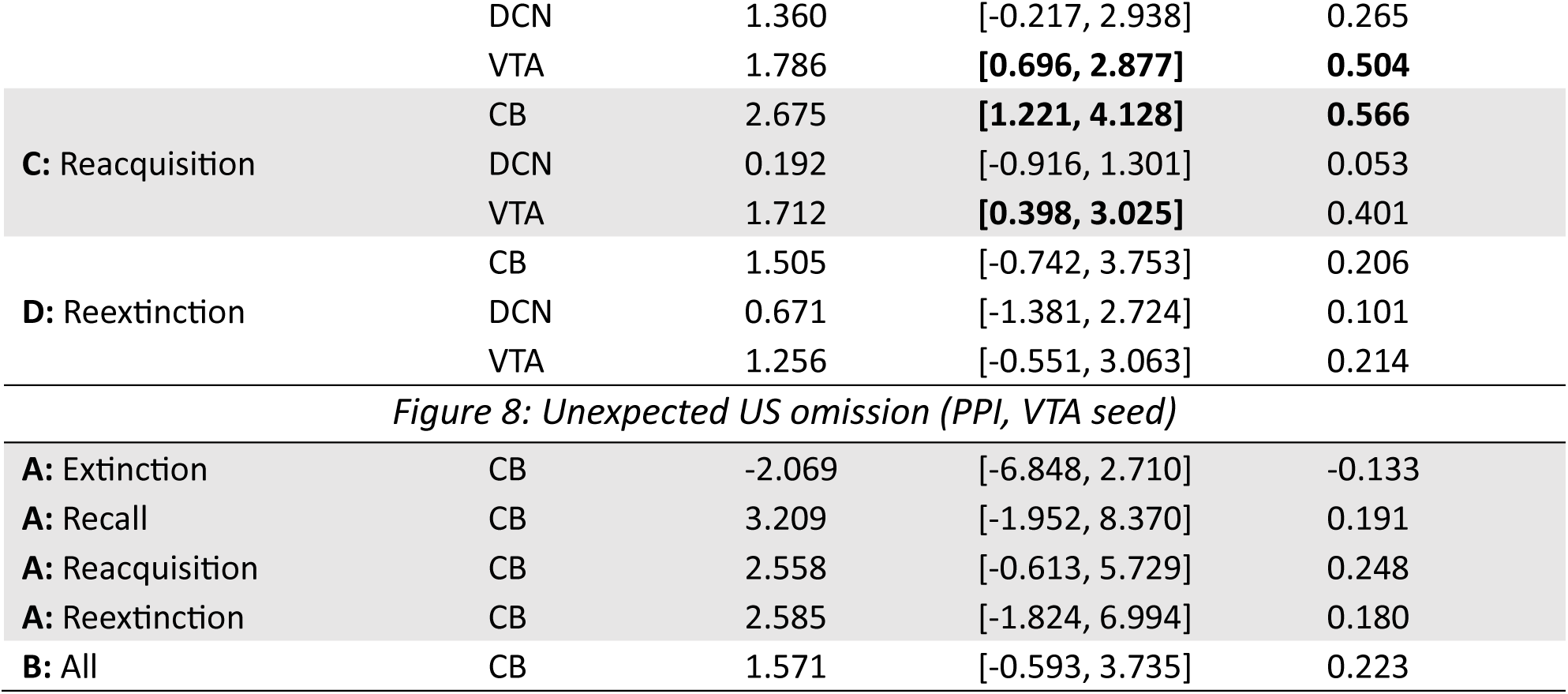
Summary statistics for subject-level VOI contrast estimates across fMRI contrasts. For each contrast and region of interest (CB: cerebellar cortex; DCN: deep cerebellar nuclei; VTA: ventral tegmental area), the table reports mean contrast estimates, 95% confidence intervals (CI), and Cohen’s d (one-sample effect size relative to zero). Effect sizes with Cohen’s d > 0.5 and 95% confidence intervals entirely above zero are highlighted in bold to facilitate interpretation of effect magnitude and consistency across participants. Event-based contrasts show comparatively consistent effects despite being based on a small number of trials (e.g., first three unexpected US omissions), whereas parametric modulation and psychophysiological interaction (PPI) analyses incorporate a larger number of observations and show greater inter-individual variability.

| Contrast | ROI | Mean | 95% CI | Cohen's *d* |
| --- | --- | --- | --- | --- |
| *Figure 4: Acquisition, US prediction and presentation* |  |  |  |  |
| **A: CS+ > CS-** | CB | 0.173 | [-0.373, 0.720] | 0.097 |
|  | DCN | -0.200 | [-0.773, 0.373] | -0.107 |
|  | VTA | 0.888 | **[0.458, 1.319]** | **0.635** |
| **B: CS+ x P (inv)** | CB | 1.872 | **[0.894, 2.849]** | **0.589** |
|  | DCN | 0.562 | [-0.406, 1.529] | 0.179 |
|  | VTA | 2.078 | **[1.281, 2.874]** | **0.803** |
| **C: US presentation** | CB | 5.367 | **[3.879, 6.855]** | **1.110** |
|  | DCN | 2.040 | **[1.131, 2.950]** | **0.691** |
|  | VTA | 2.852 | **[2.064, 3.639]** | **1.115** |
| *Figure 5: Extinction, US prediction and omission* |  |  |  |  |
| **A: CS+ > CS-** | CB | -0.057 | [-0.566, 0.452] | -0.034 |
|  | DCN | -0.143 | [-0.649, 0.363] | -0.087 |
|  | VTA | 0.223 | [-0.170, 0.615] | 0.175 |
| **B: CS+ x Prediction** | CB | 1.043 | [-0.083, 2.169] | 0.285 |
|  | DCN | 0.356 | [-0.468, 1.180] | 0.133 |
|  | VTA | 2.057 | **[1.248, 2.865]** | **0.783** |
| **C: US presentation** | CB | 0.471 | [-0.062, 1.004] | 0.272 |
|  | DCN | 0.311 | [-0.208, 0.829] | 0.184 |
|  | VTA | 0.120 | [-0.351, 0.590] | 0.078 |
| *Figure 6: Unexpected US omission (event-based)* |  |  |  |  |
| **A: Extinction** | CB | 4.487 | **[2.762, 6.211]** | **0.801** |
|  | DCN | 1.989 | **[0.653, 3.325]** | 0.458 |
|  | VTA | 1.807 | **[0.616, 2.997]** | 0.467 |
| **B: Recall** | CB | 4.485 | **[2.823, 6.148]** | **0.830** |
|  | DCN | 1.352 | **[0.072, 2.632]** | 0.325 |
|  | VTA | 1.949 | **[0.903, 2.996]** | **0.573** |
| **C: Reacquisition** | CB | 8.082 | **[6.209, 9.955]** | **1.328** |
|  | DCN | 0.872 | [-0.379, 2.123] | 0.215 |
|  | VTA | 3.711 | **[2.133, 5.289]** | **0.724** |
| **D: Reextinction** | CB | 5.780 | **[3.748, 7.811]** | **0.876** |
|  | DCN | 0.969 | [-0.424, 2.361] | 0.214 |
|  | VTA | 3.152 | **[1.699, 4.605]** | **0.668** |
| *Figure 7: Unexpected US omission (parametric modulation)* |  |  |  |  |
| **A: Extinction** | CB | 3.409 | **[2.034, 4.784]** | **0.763** |
|  | DCN | 1.114 | **[0.259, 1.970]** | 0.401 |
|  | VTA | 1.871 | **[0.966, 2.777]** | **0.636** |
| **B: Recall** | CB | 4.029 | **[2.818, 5.241]** | **1.024** |
|  | DCN | 1.360 | [-0.217, 2.938] | 0.265 |
|  | VTA | 1.786 | **[0.696, 2.877]** | **0.504** |
| **C: Reacquisition** | CB | 2.675 | **[1.221, 4.128]** | **0.566** |
|  | DCN | 0.192 | [-0.916, 1.301] | 0.053 |
|  | VTA | 1.712 | **[0.398, 3.025]** | 0.401 |
| **D: Reextinction** | CB | 1.505 | [-0.742, 3.753] | 0.206 |
|  | DCN | 0.671 | [-1.381, 2.724] | 0.101 |
|  | VTA | 1.256 | [-0.551, 3.063] | 0.214 |
| **A: Extinction** | CB | -2.069 | [-6.848, 2.710] | -0.133 |
| **A: Recall** | CB | 3.209 | [-1.952, 8.370] | 0.191 |
| **A: Reacquisition** | CB | 2.558 | [-0.613, 5.729] | 0.248 |
| **A: Reextinction** | CB | 2.585 | [-1.824, 6.994] | 0.180 |
| **B: All** | CB | 1.571 | [-0.593, 3.735] | 0.223 |

#### Cerebellar VOI used for connectivity analyses on a SUIT flatmap

**Figure S4:**
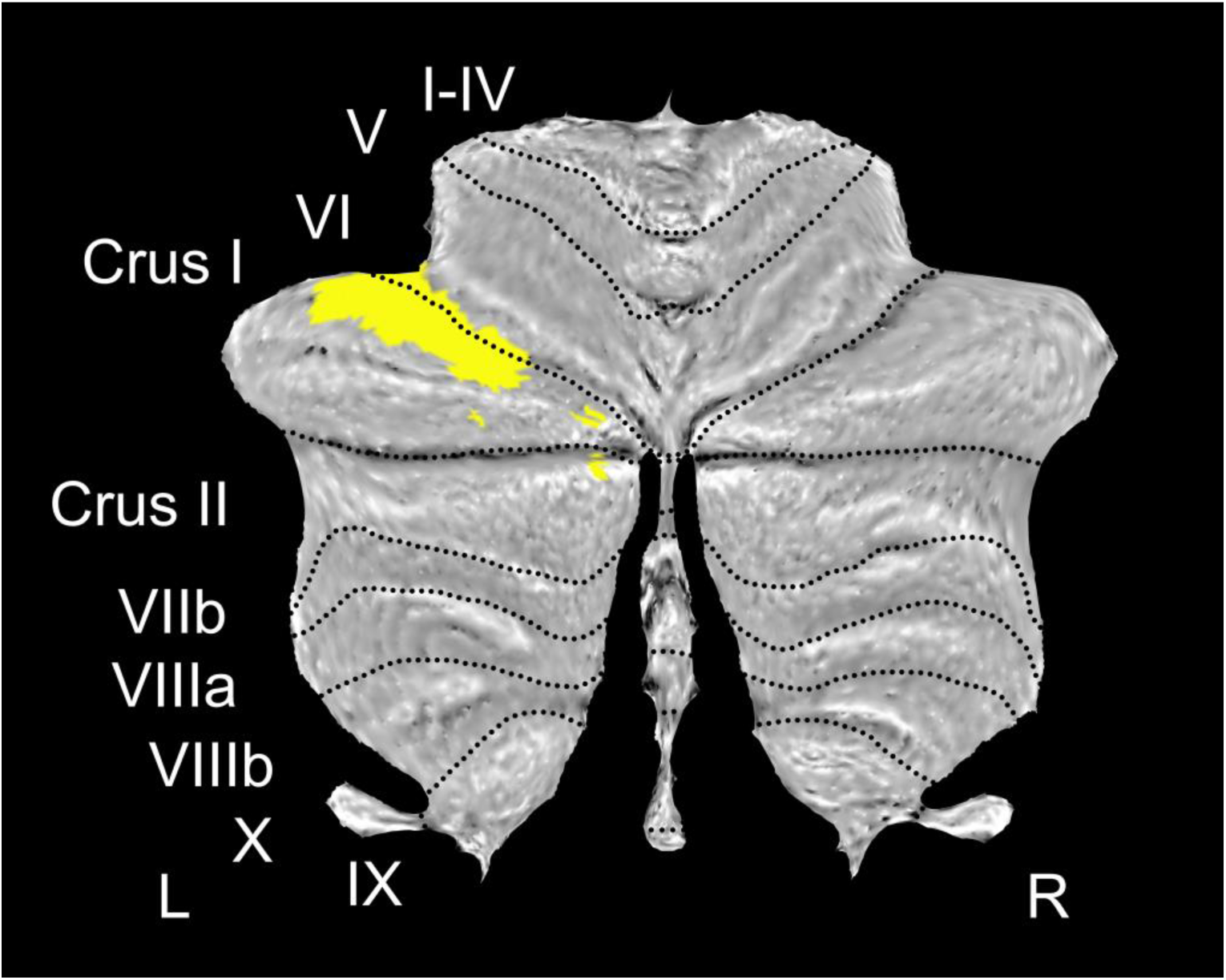
SUIT cerebellar flatmap^22^ showing the cerebellar volume of interest (VOI; CB) used for the DCM and PPI analyses. The VOI (yellow) is restricted to the cerebellar cortex and was defined as the conjunction of unexpected US omission contrasts (see Methods: Volumes of interest (VOI) definition). This VOI was used as the cerebellar node for the DCM (Figure 8) and as the seed region for the PPI analysis (Figure S3).

### fMRI activation cluster tables

#### fMRI activations related to the prediction and presentation of the US. TFCE and FWE corrected

**Table S8:**
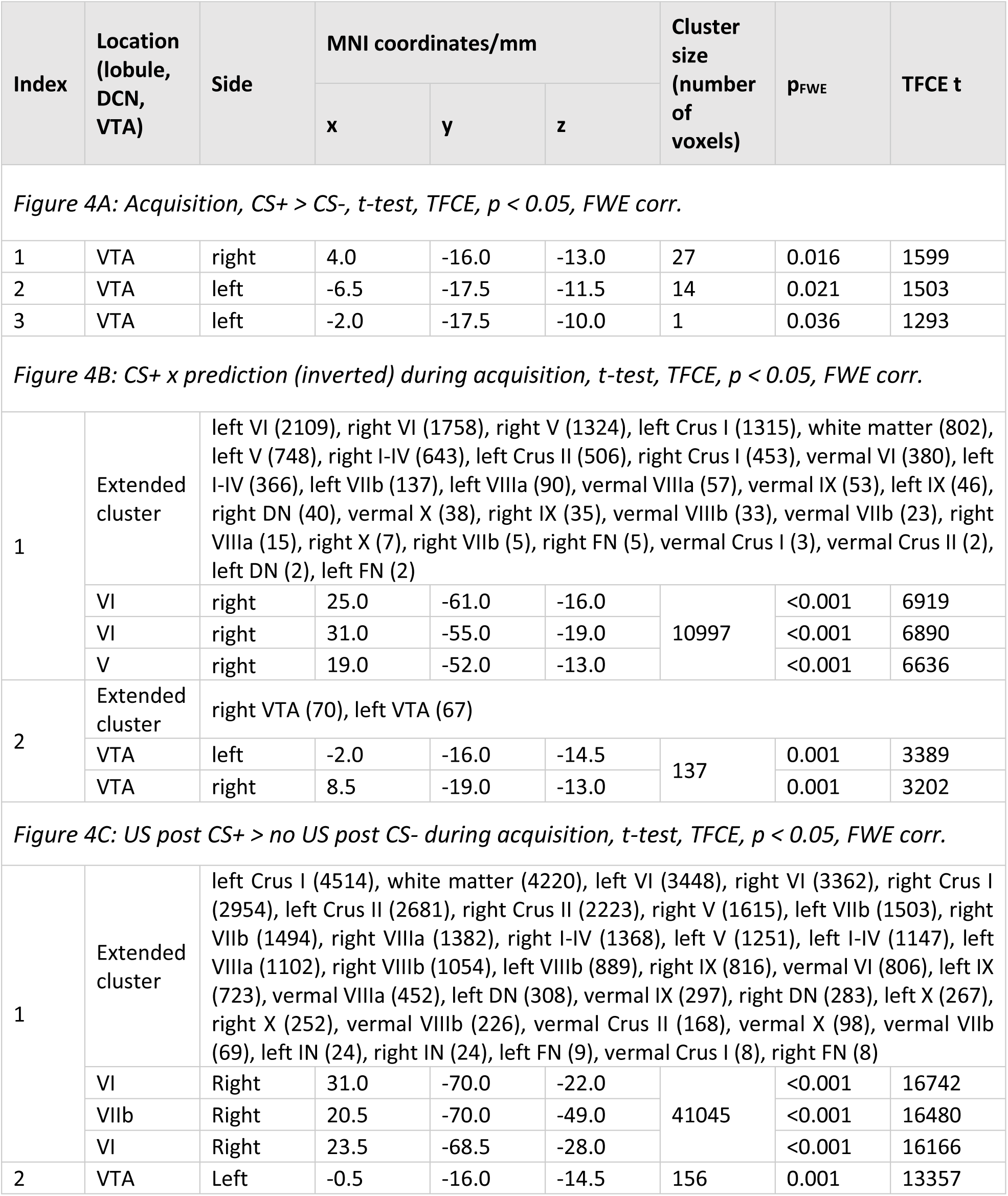
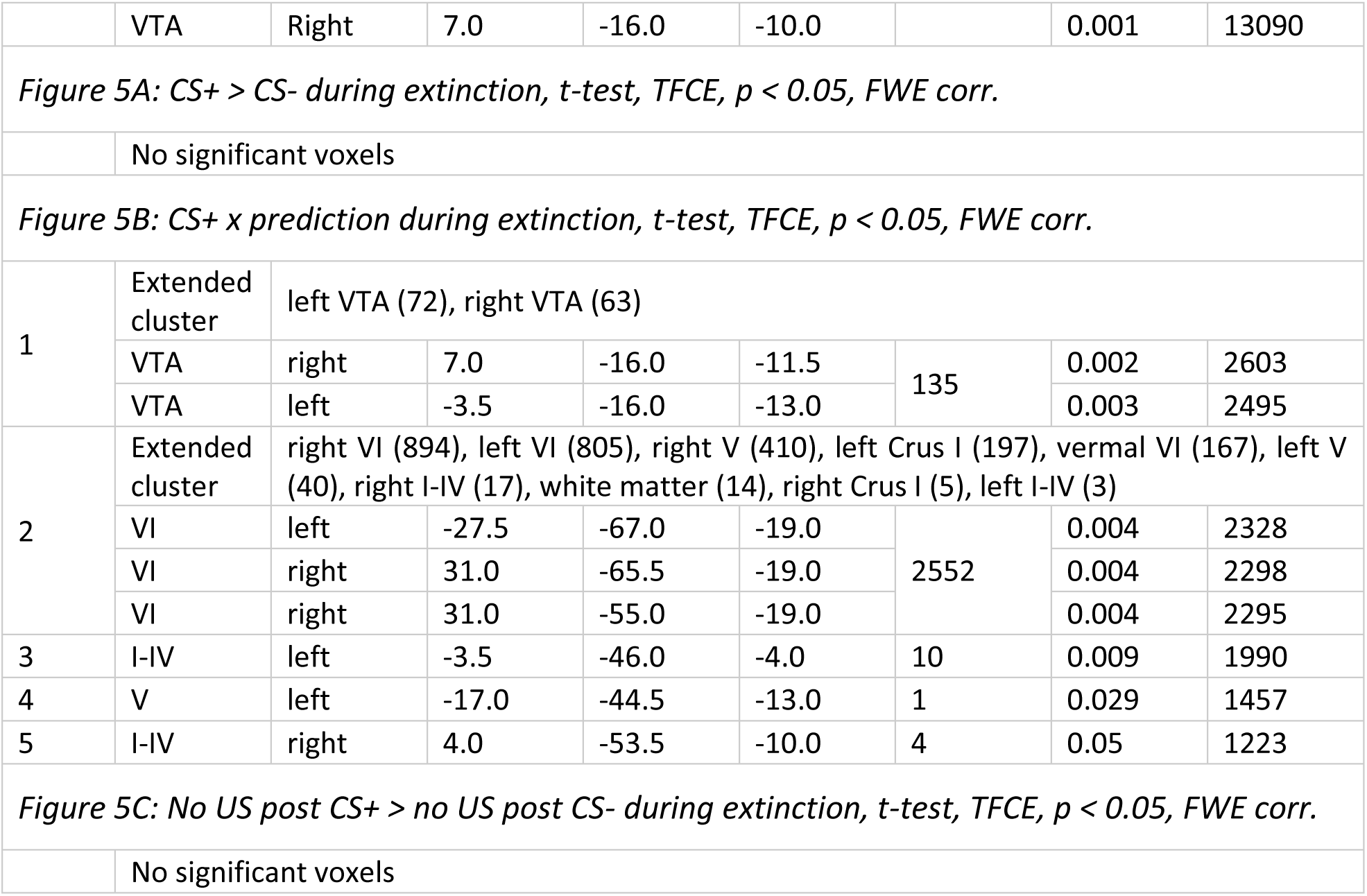
fMRI activation clusters related to the prediction, presentation and omission of the unconditioned stimulus (US) during acquisition and extinction training (Figure 4 and 5). Clusters were identified in the cerebellar cortex, deep cerebellar nuclei (DCN), and ventral tegmental area (VTA) using threshold-free cluster enhancement (TFCE) with family-wise error (FWE) correction (p < 0.05). Up to three local maxima per cluster are reported, separated by at least 8 mm. Coordinates are given in MNI space (x, y, z). Cluster size is reported as number of voxels (voxel volume = 3.375 mm³). US: unconditioned stimulus; CS: conditioned stimulus; VTA: ventral tegmental area; DCN: deep cerebellar nuclei; DN: dentate nucleus; IN: interposed nucleus; FN: fastigial nucleus; MNI: Montreal Neurological Institute standard brain; TFCE t: threshold-free cluster-enhanced t-statistic; pFWE: family-wise error-corrected p-value.

#### fMRI activations related to the prediction and presentation of the US. Uncorrected

**Table S9:**
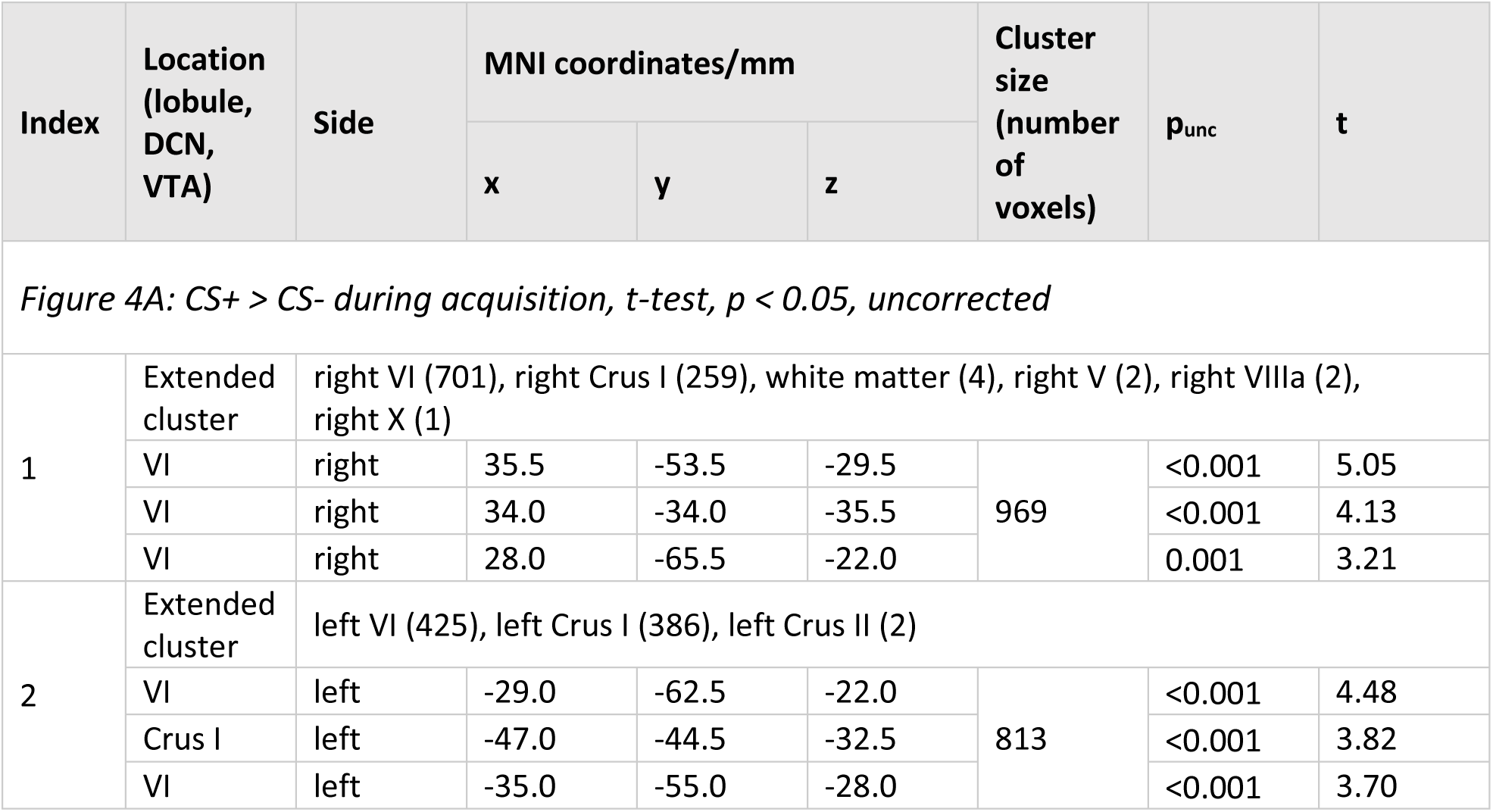
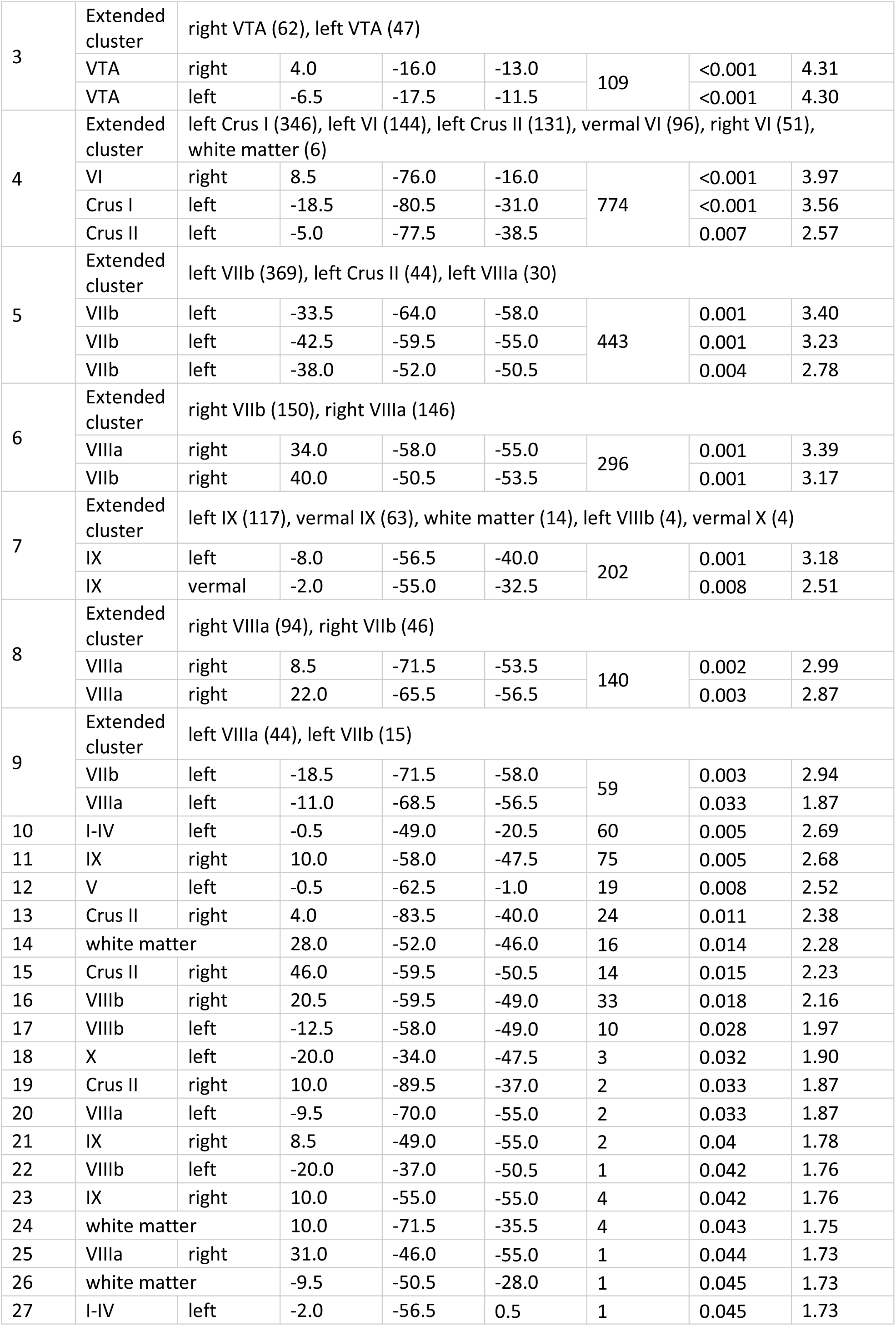
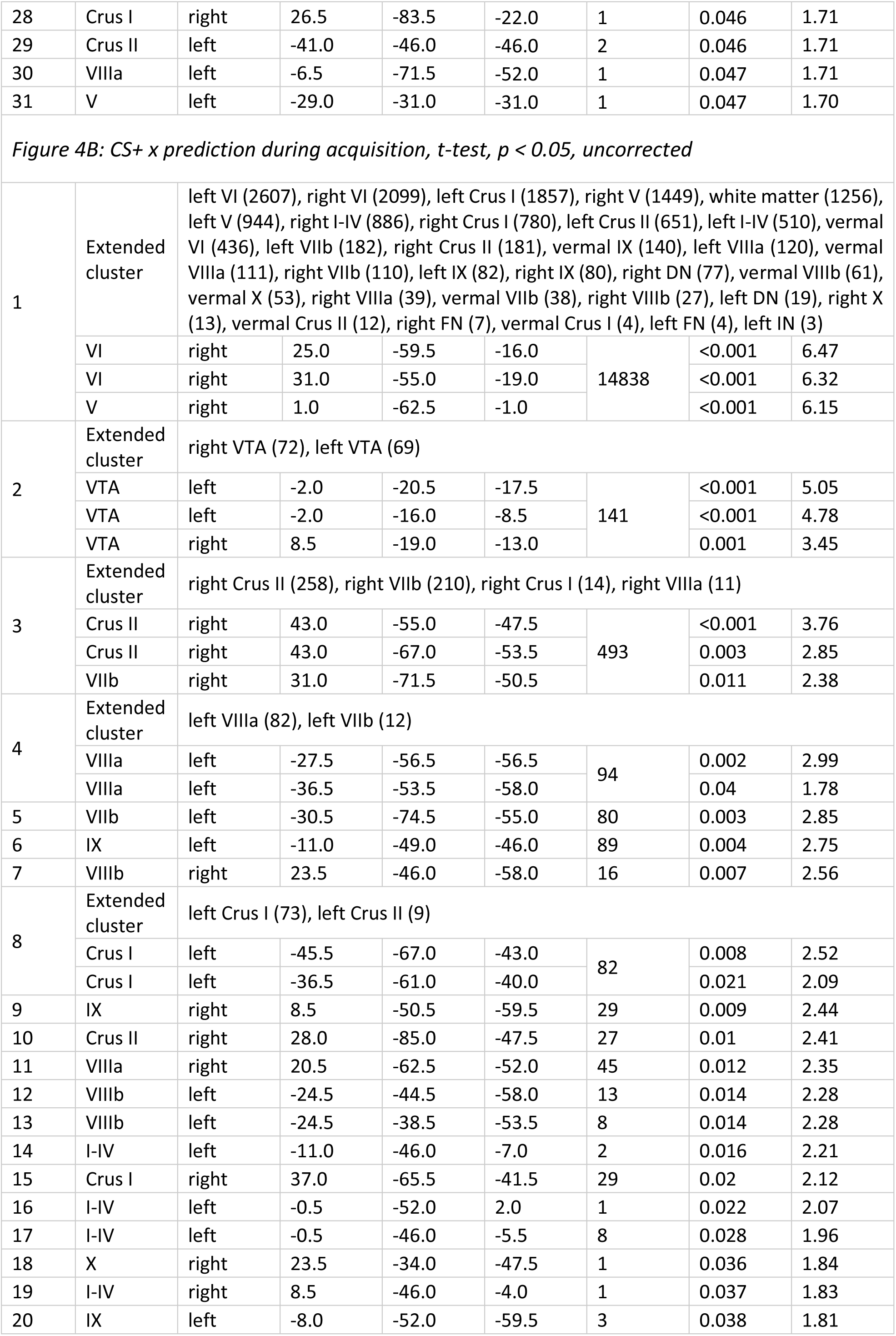
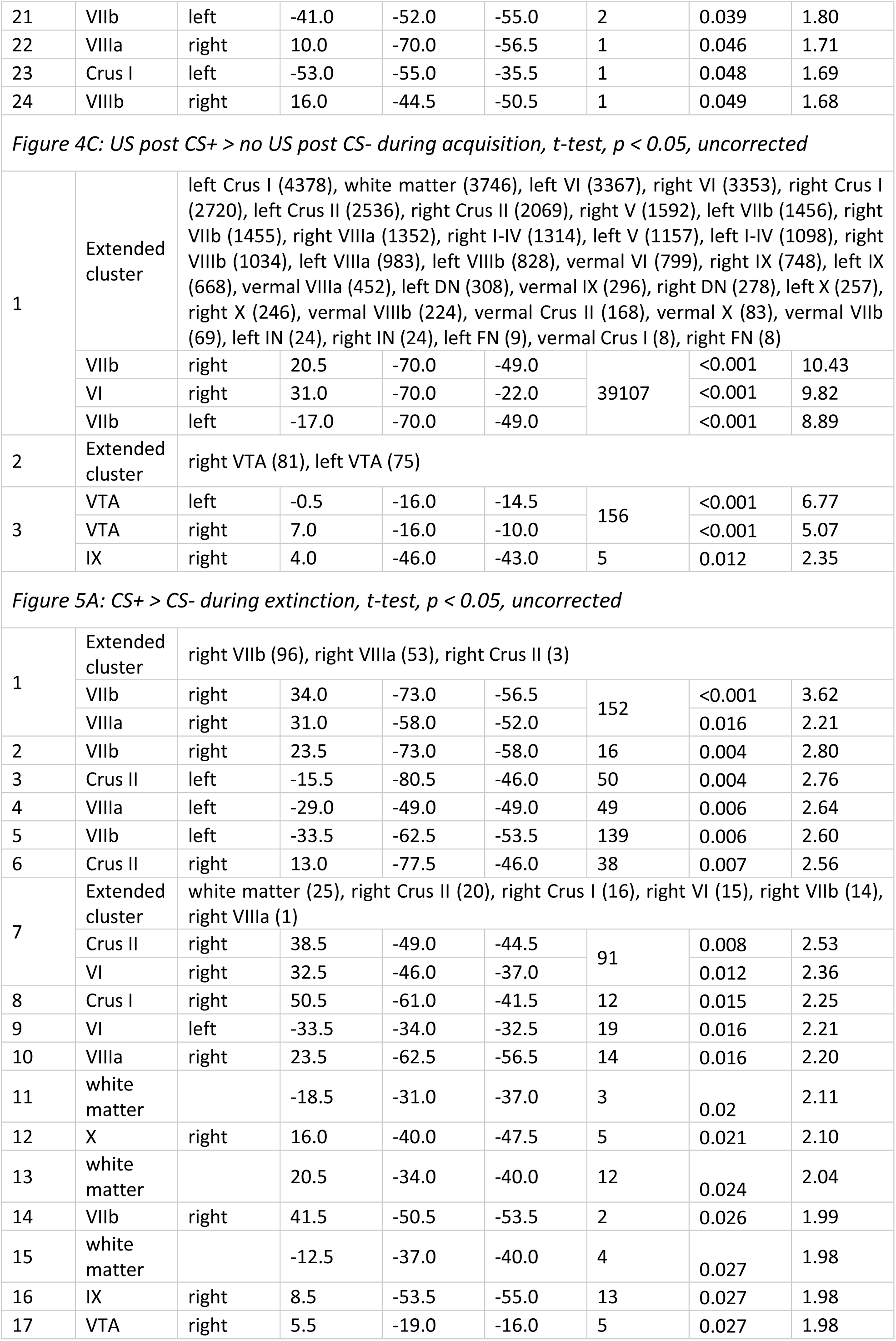
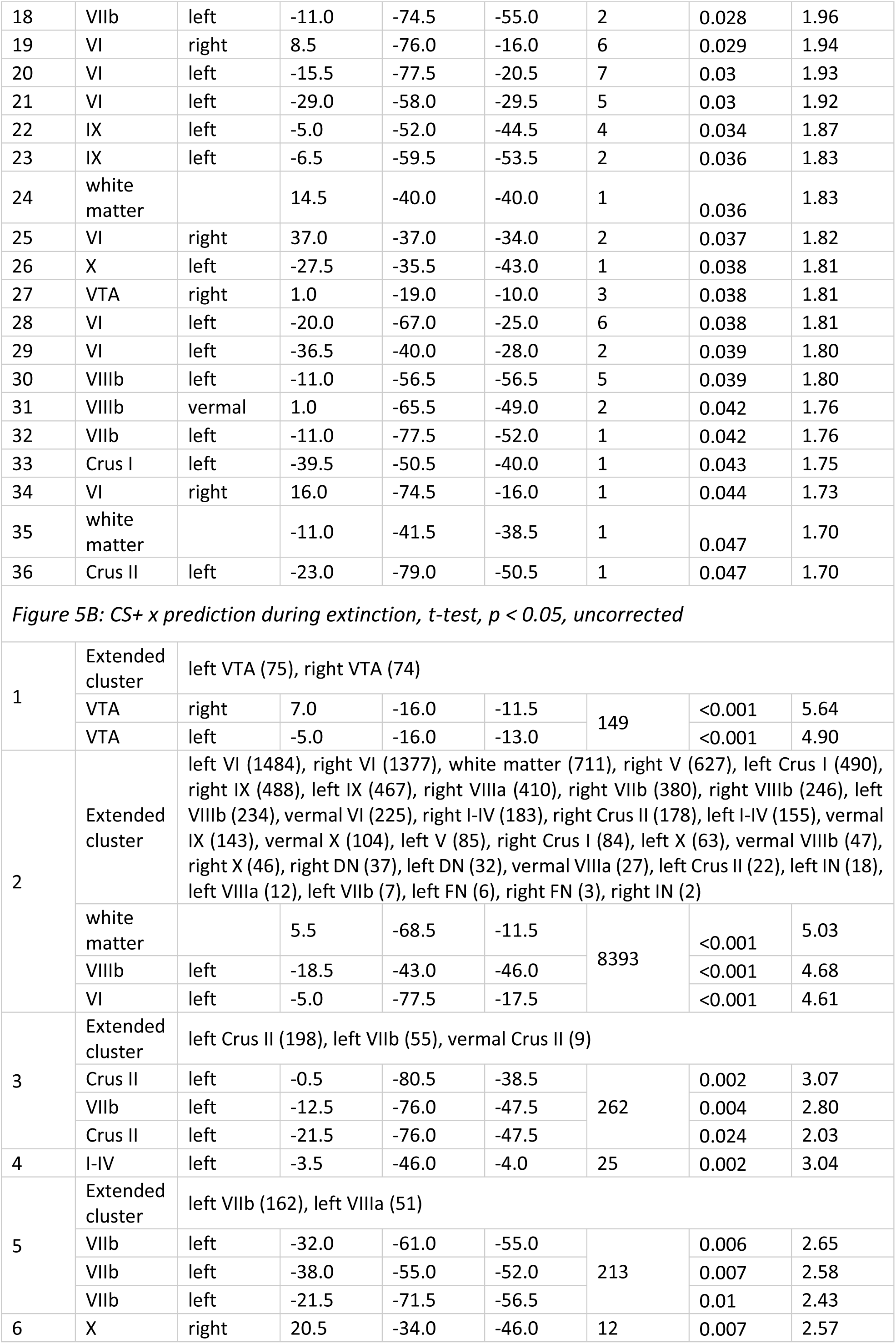
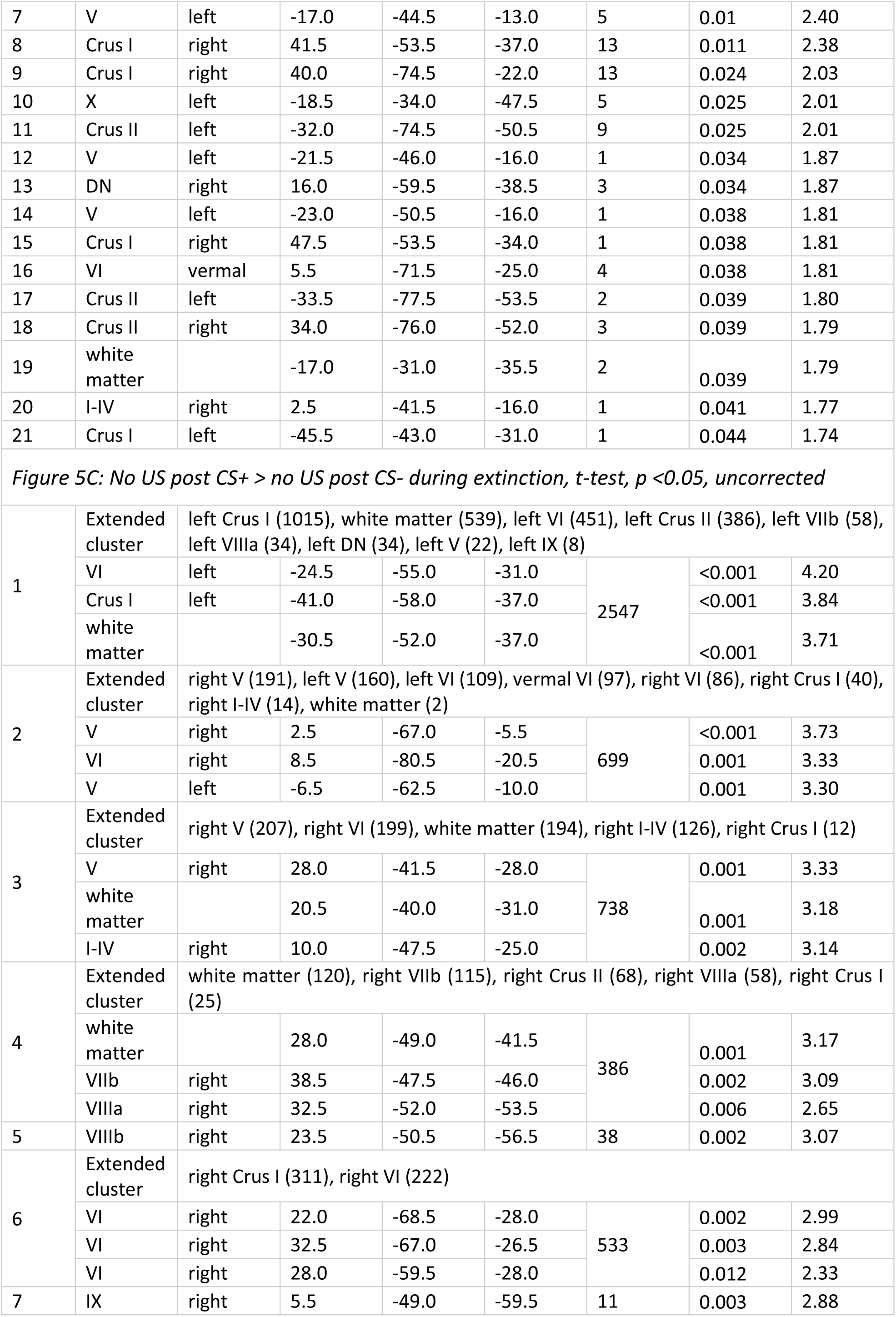
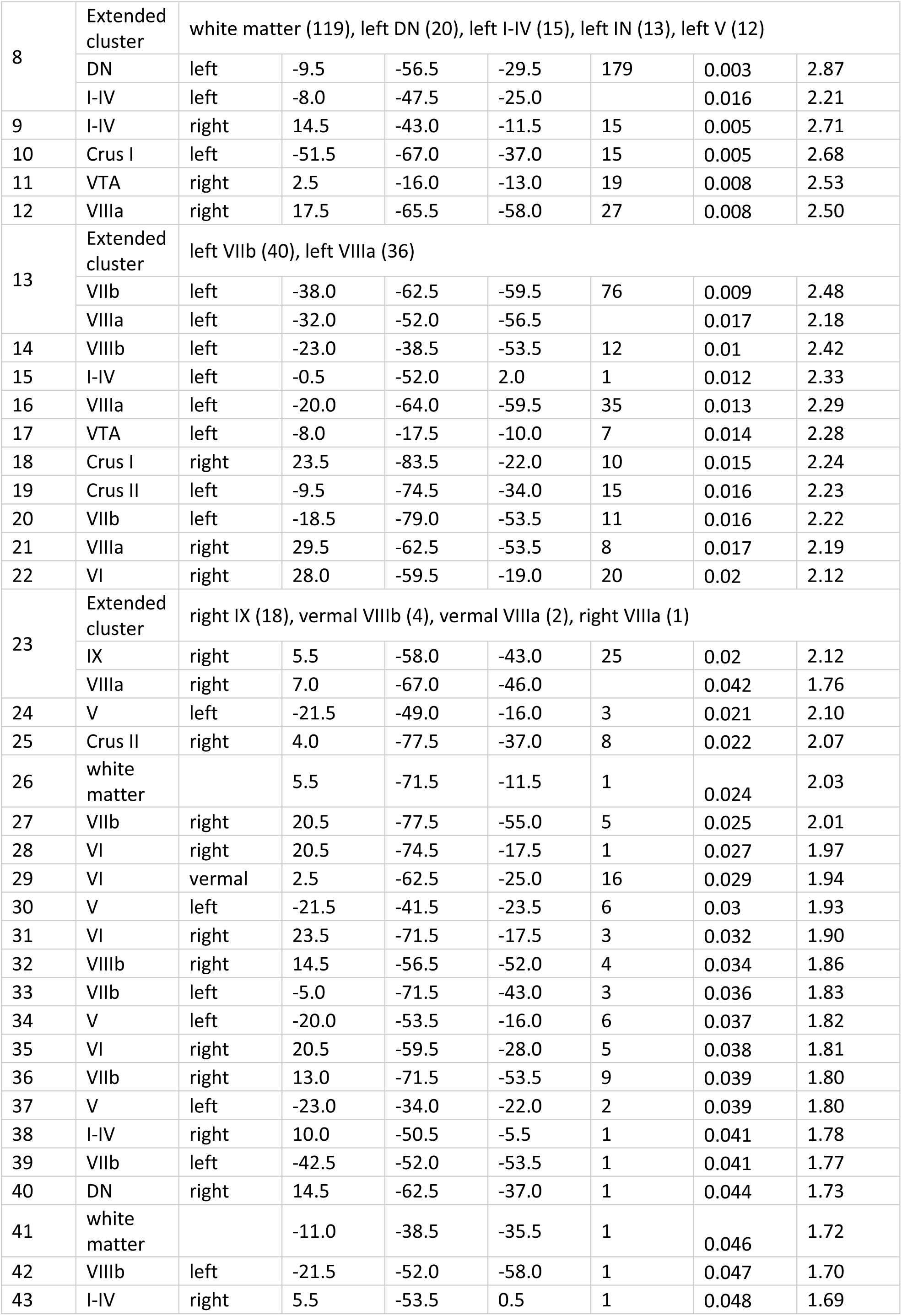
fMRI activation clusters (p < 0.05, uncorrected) related to prediction, presentation and omission of the unconditioned stimulus (US) during acquisition and extinction training (Figure 4 and 5). Clusters were identified in the cerebellar cortex, deep cerebellar nuclei (DCN), and ventral tegmental area (VTA). Up to three local maxima per cluster are reported, separated by at least 8 mm. Coordinates are given in MNI space (x, y, z). Cluster size is reported as number of voxels (voxel volume = 3.375 mm³). US: unconditioned stimulus; CS: conditioned stimulus; VTA: ventral tegmental area; DCN: deep cerebellar nuclei; DN: dentate nucleus; IN: interposed nucleus; FN: fastigial nucleus; MNI: Montreal Neurological Institute standard brain; t: t-statistic; punc: uncorrected p-value.

#### fMRI activations related to the unexpected omission of the US. TFCE and FWE corrected

**Table S10:**
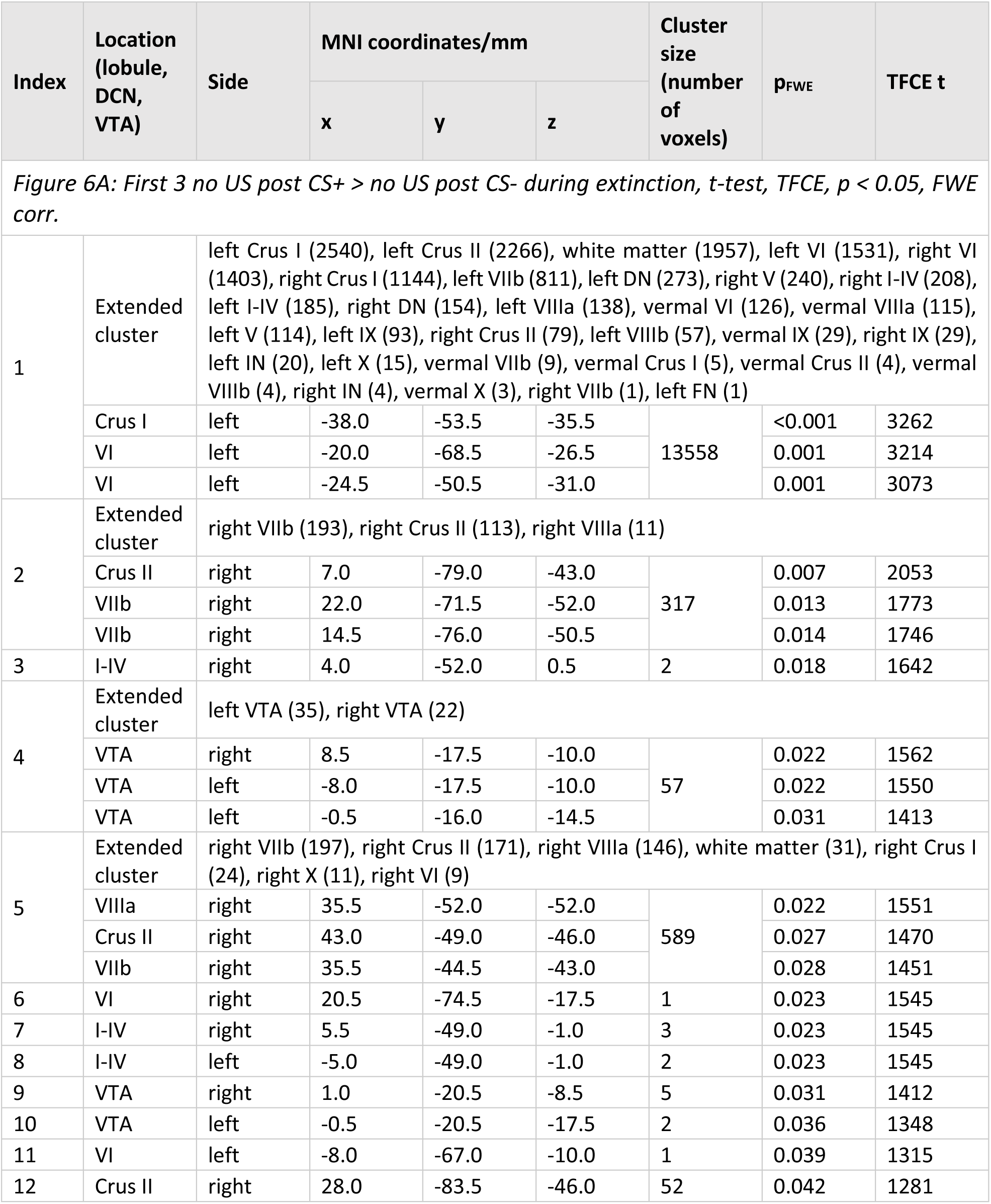
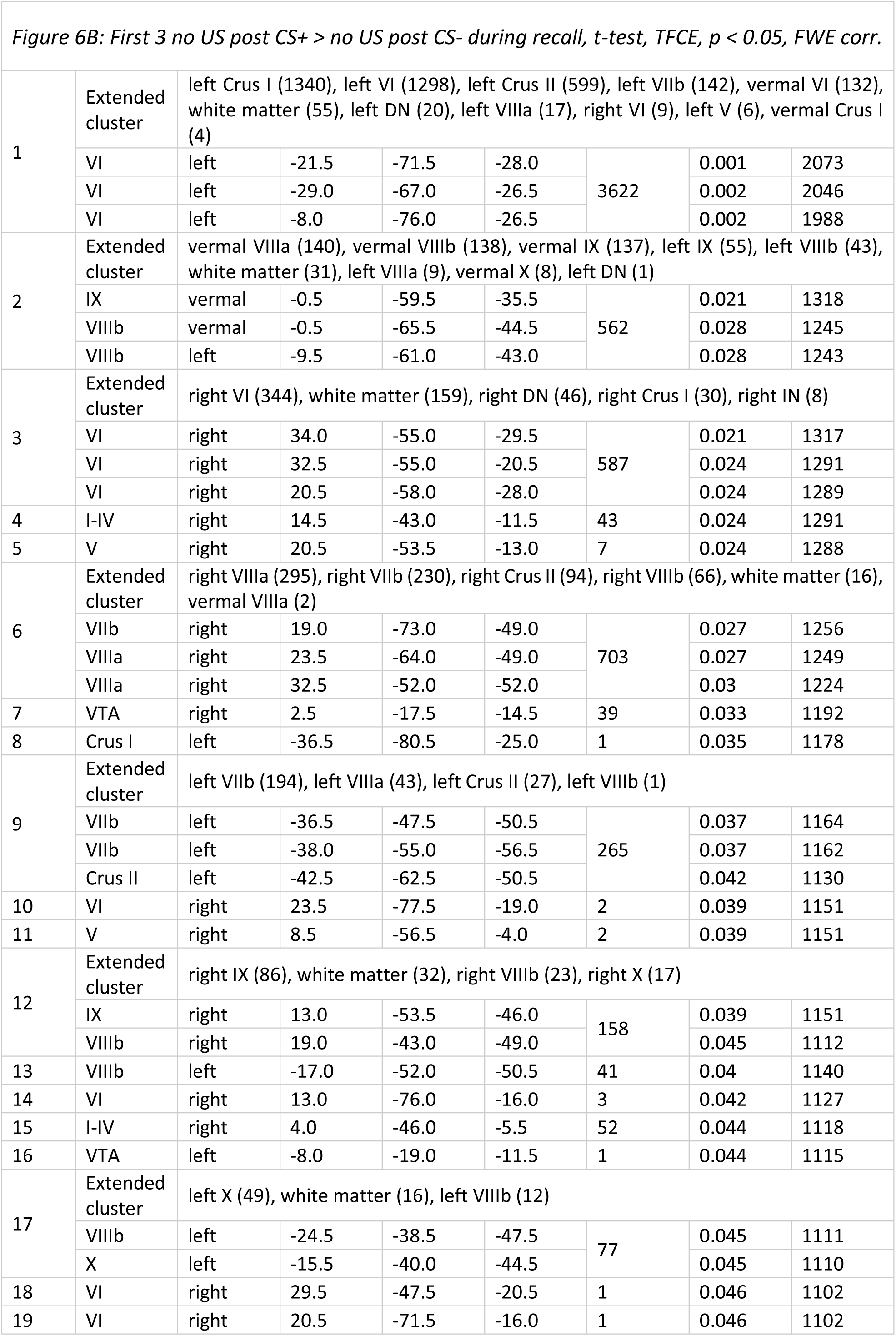
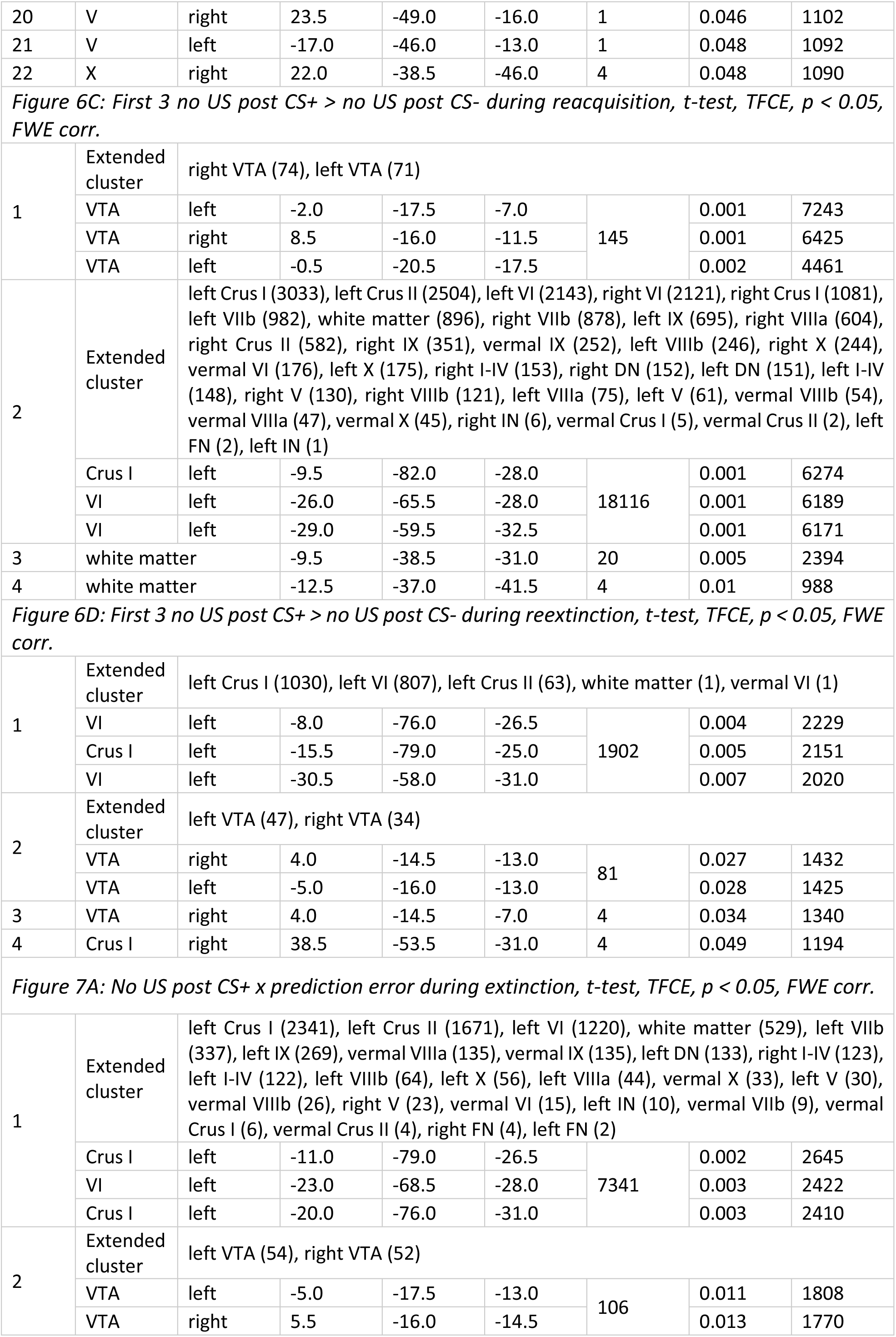
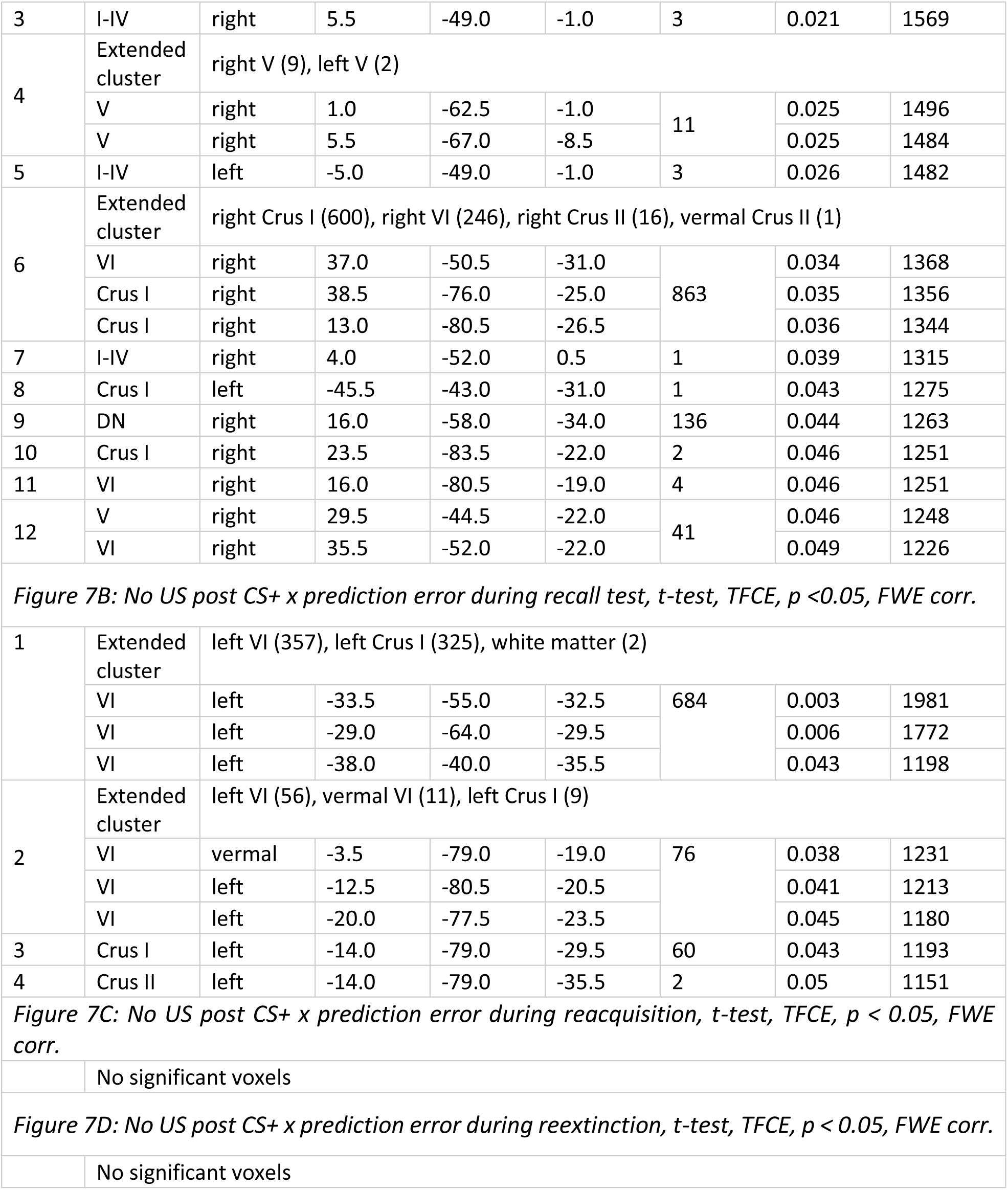
fMRI activation clusters related to the unexpected omission of the unconditioned stimulus (US) during extinction training (Figure 6 and 7). Clusters were identified in the cerebellar cortex, deep cerebellar nuclei (DCN), and ventral tegmental area (VTA) using threshold-free cluster enhancement (TFCE) with family-wise error (FWE) correction (p < 0.05). Up to three local maxima per cluster are reported, separated by at least 8 mm. Coordinates are given in MNI space (x, y, z). Cluster size is reported as number of voxels (voxel volume = 3.375 mm³). US: unconditioned stimulus; CS: conditioned stimulus; VTA: ventral tegmental area; DCN: deep cerebellar nuclei; DN: dentate nucleus; IN: interposed nucleus; FN: fastigial nucleus; MNI: Montreal Neurological Institute standard brain; TFCE t: threshold-free cluster-enhanced t-statistic; pFWE: family-wise error-corrected p-value.

#### fMRI activations related to the unexpected omission of the US. Uncorrected

**Table S11:**
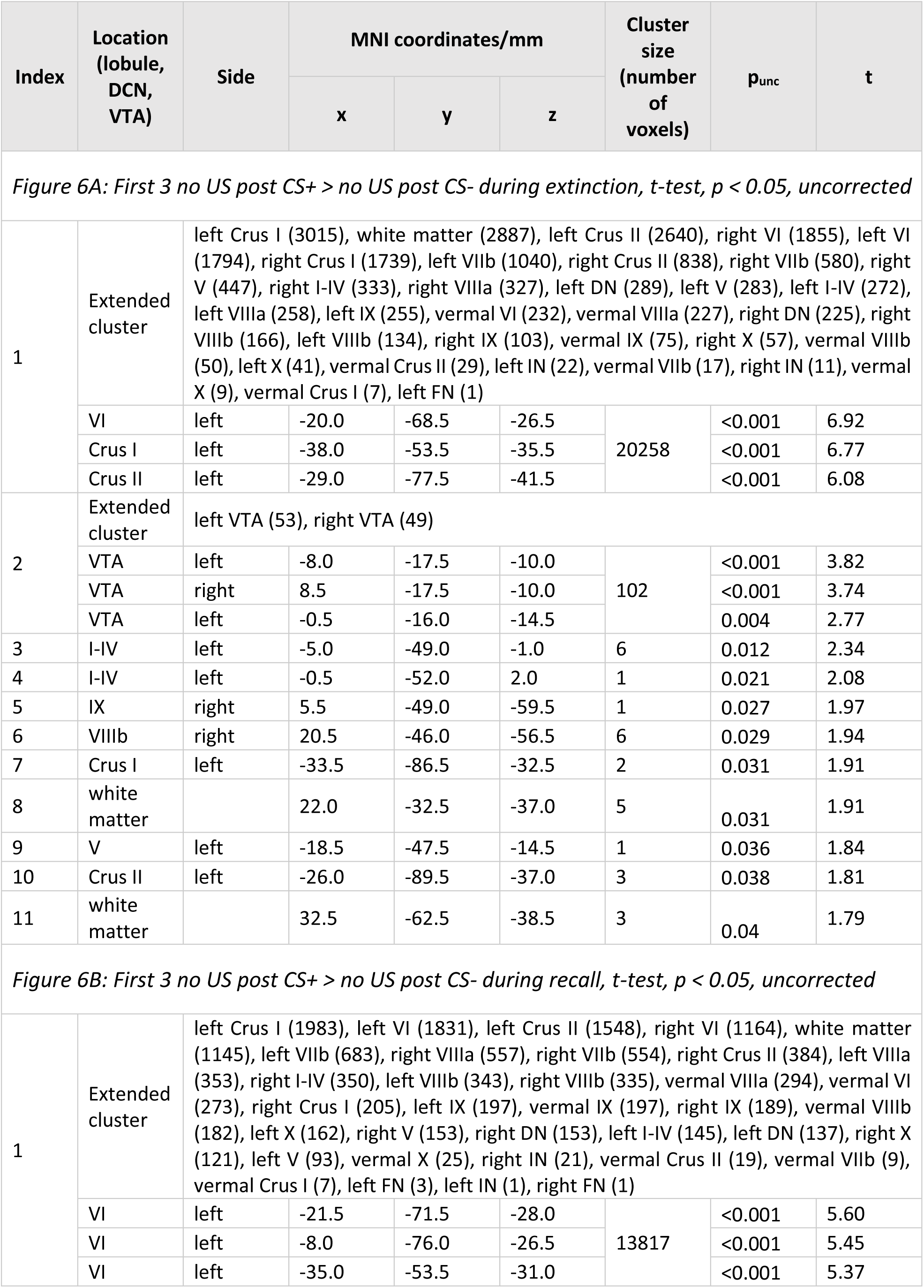
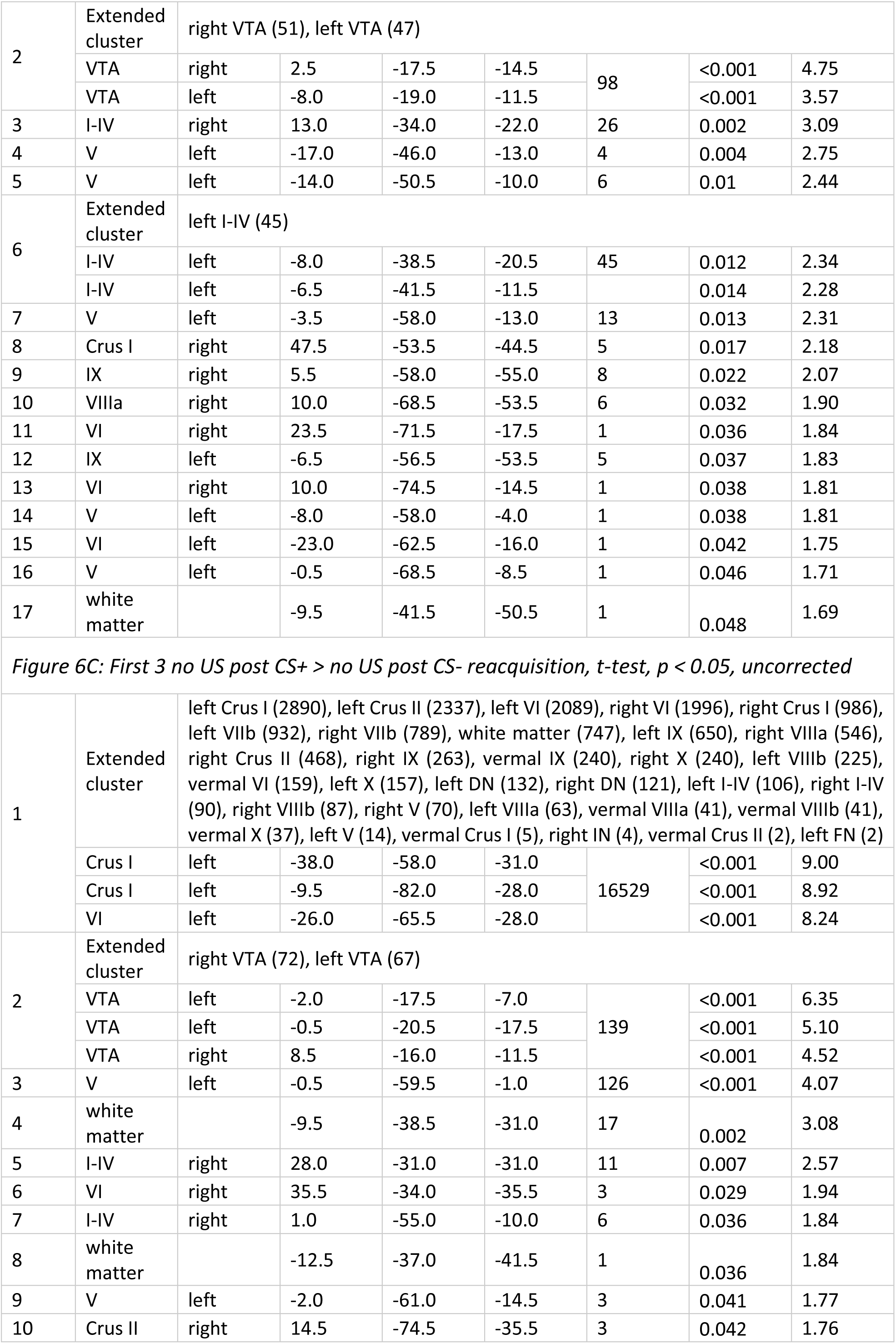
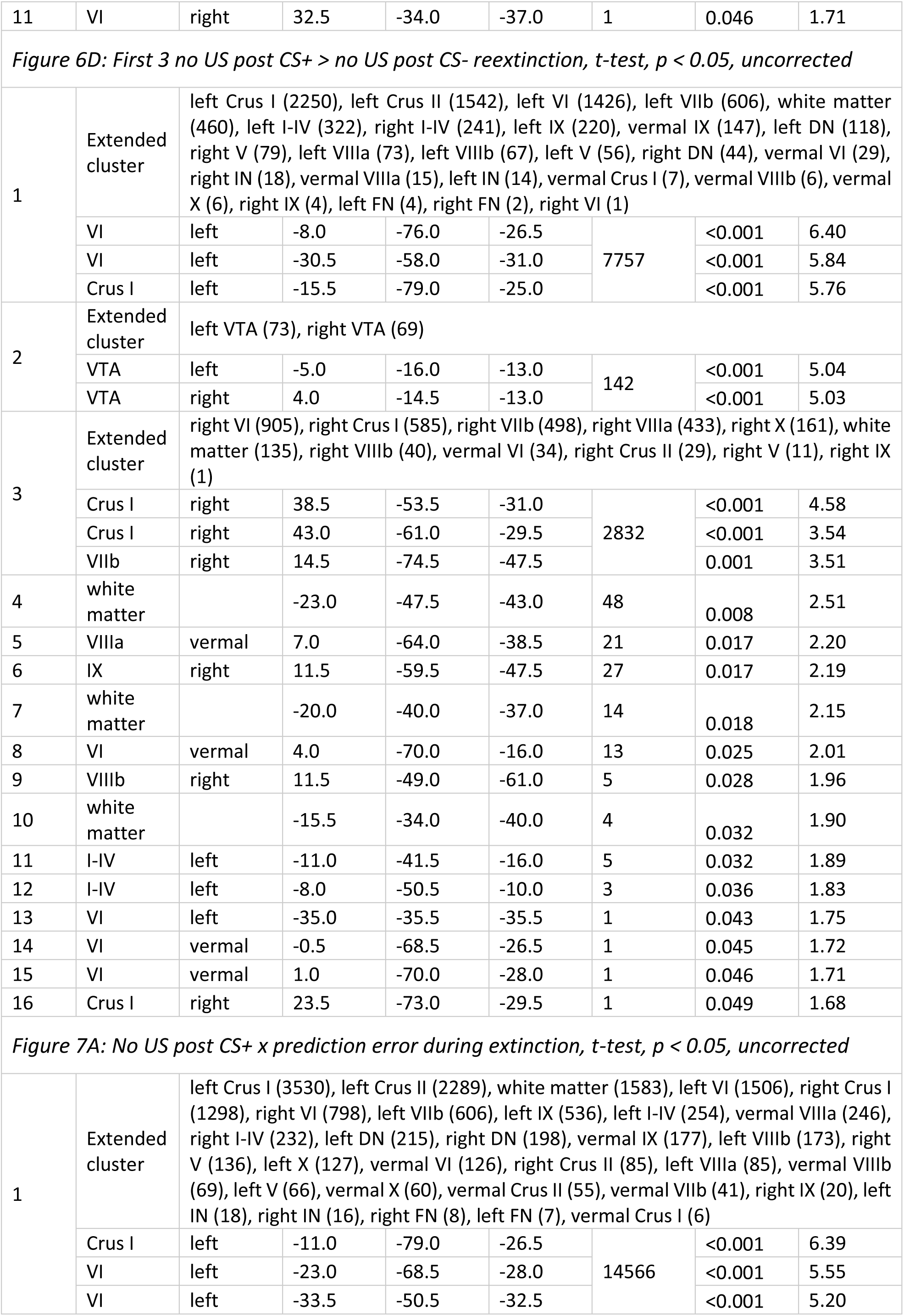
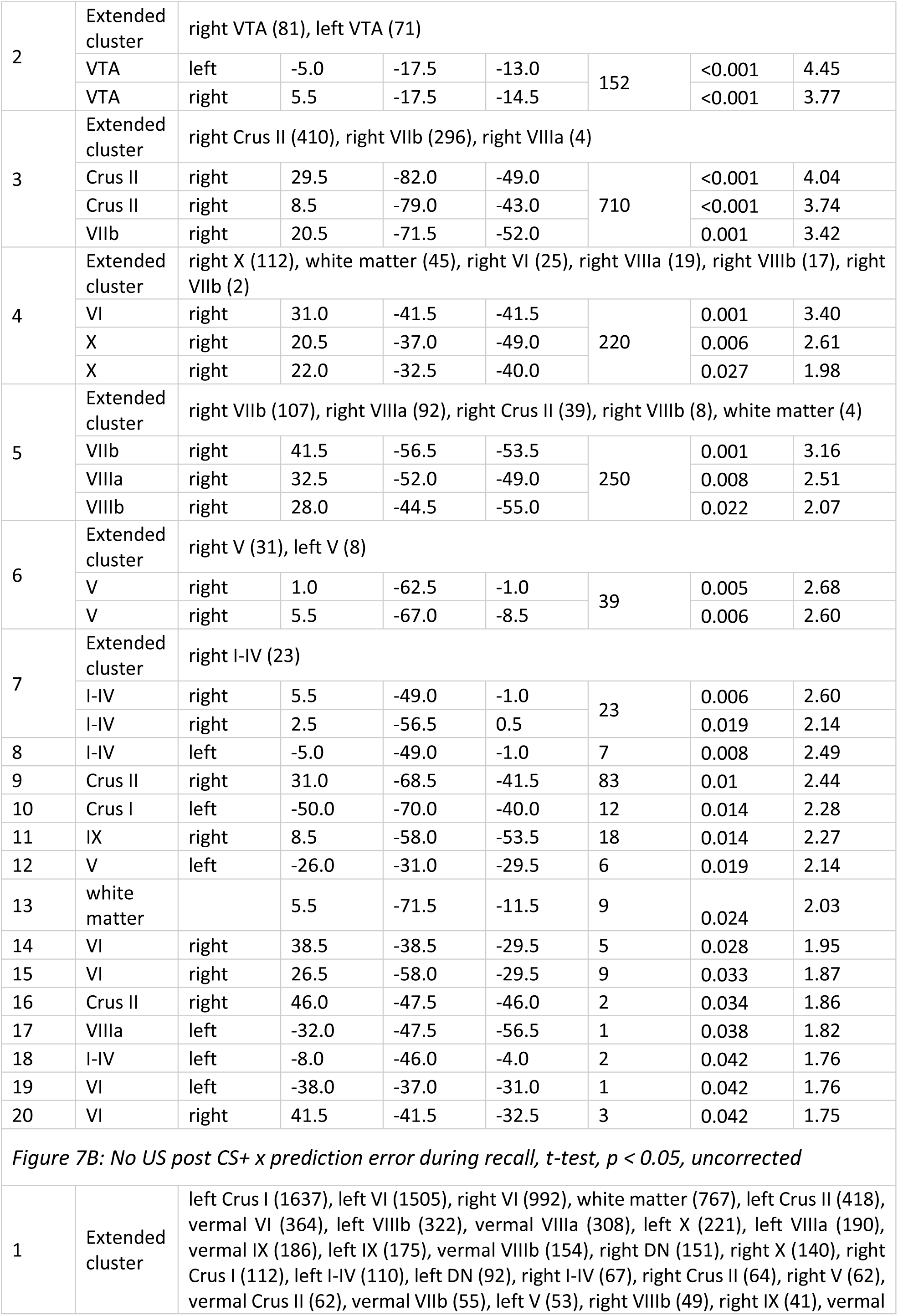
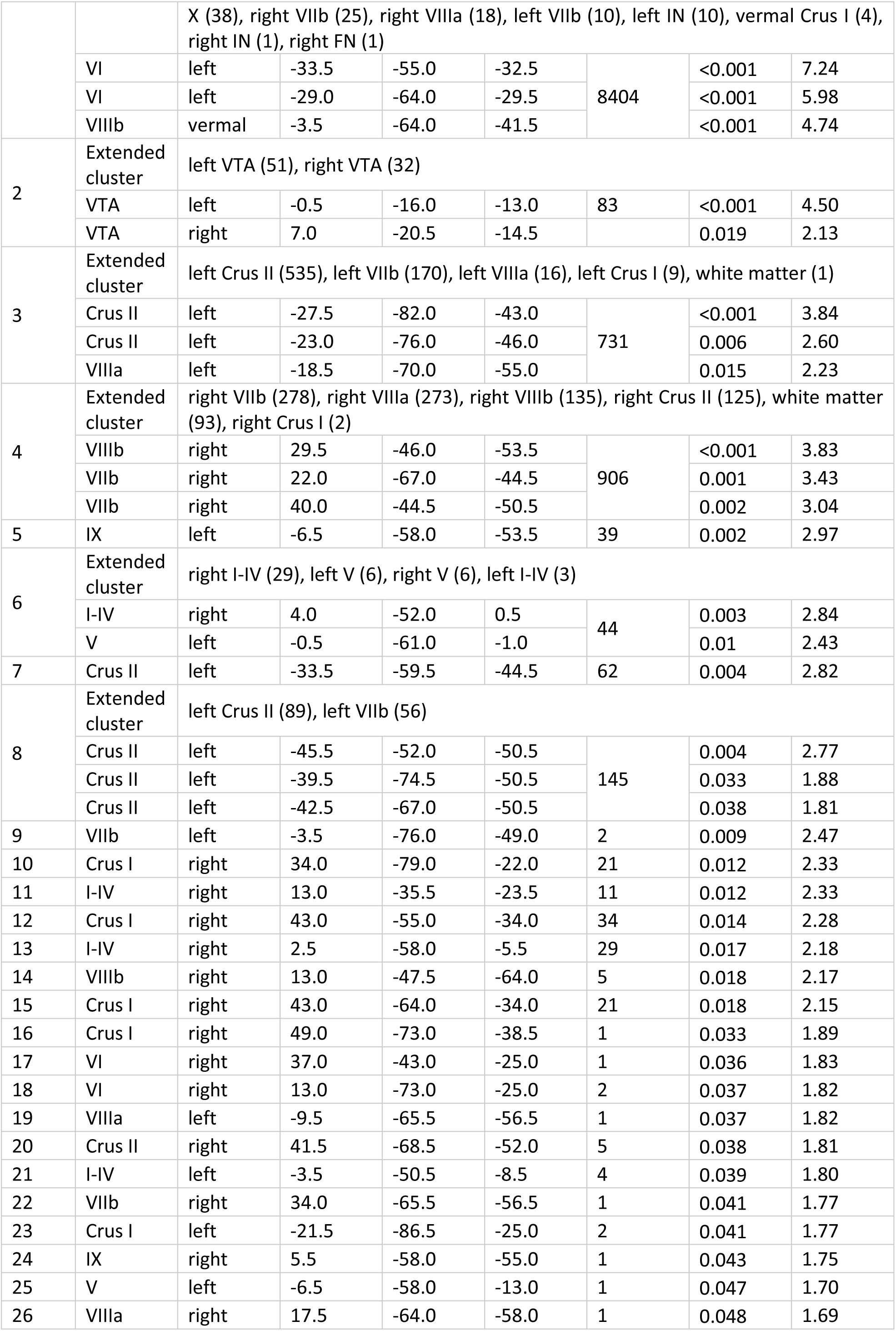
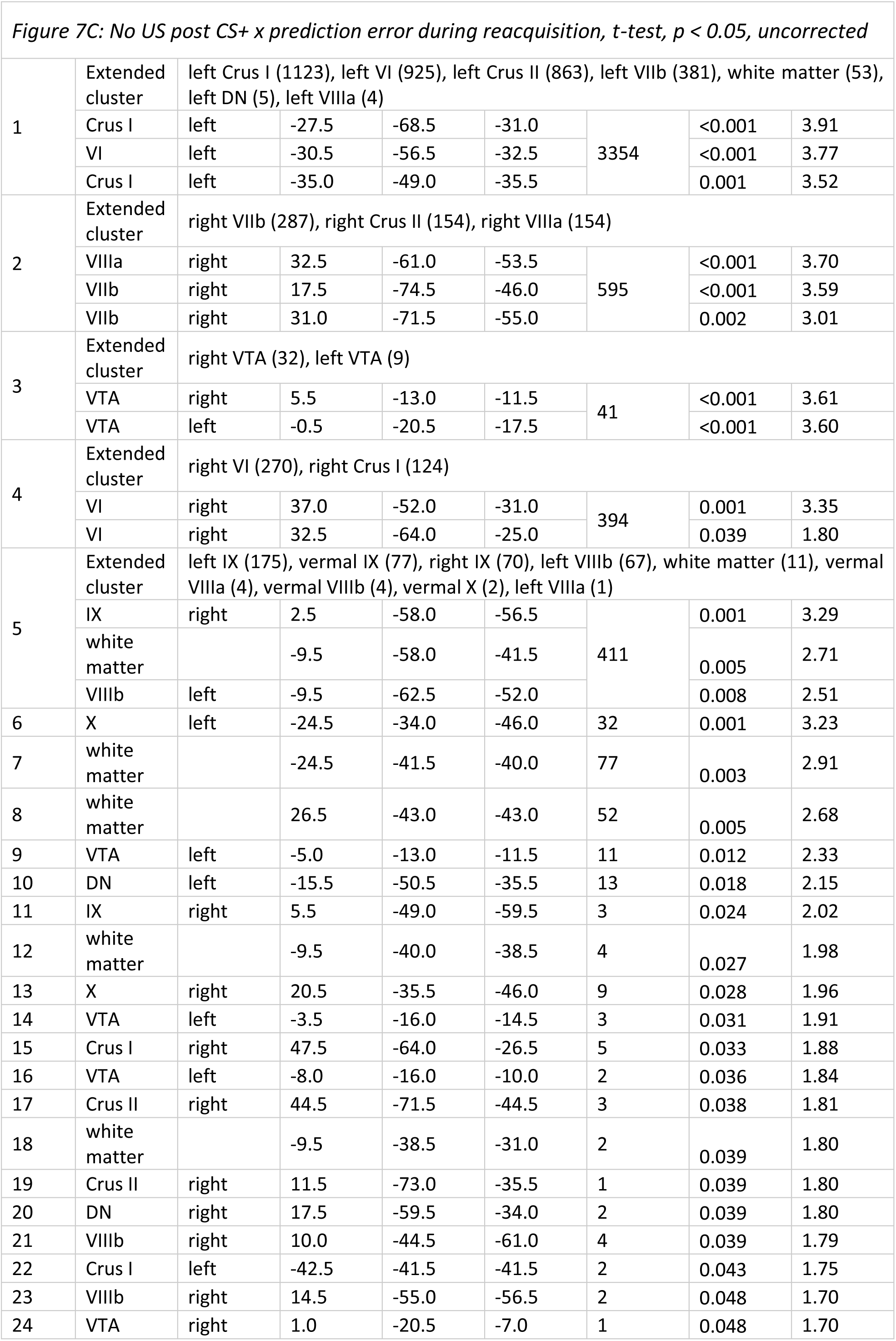
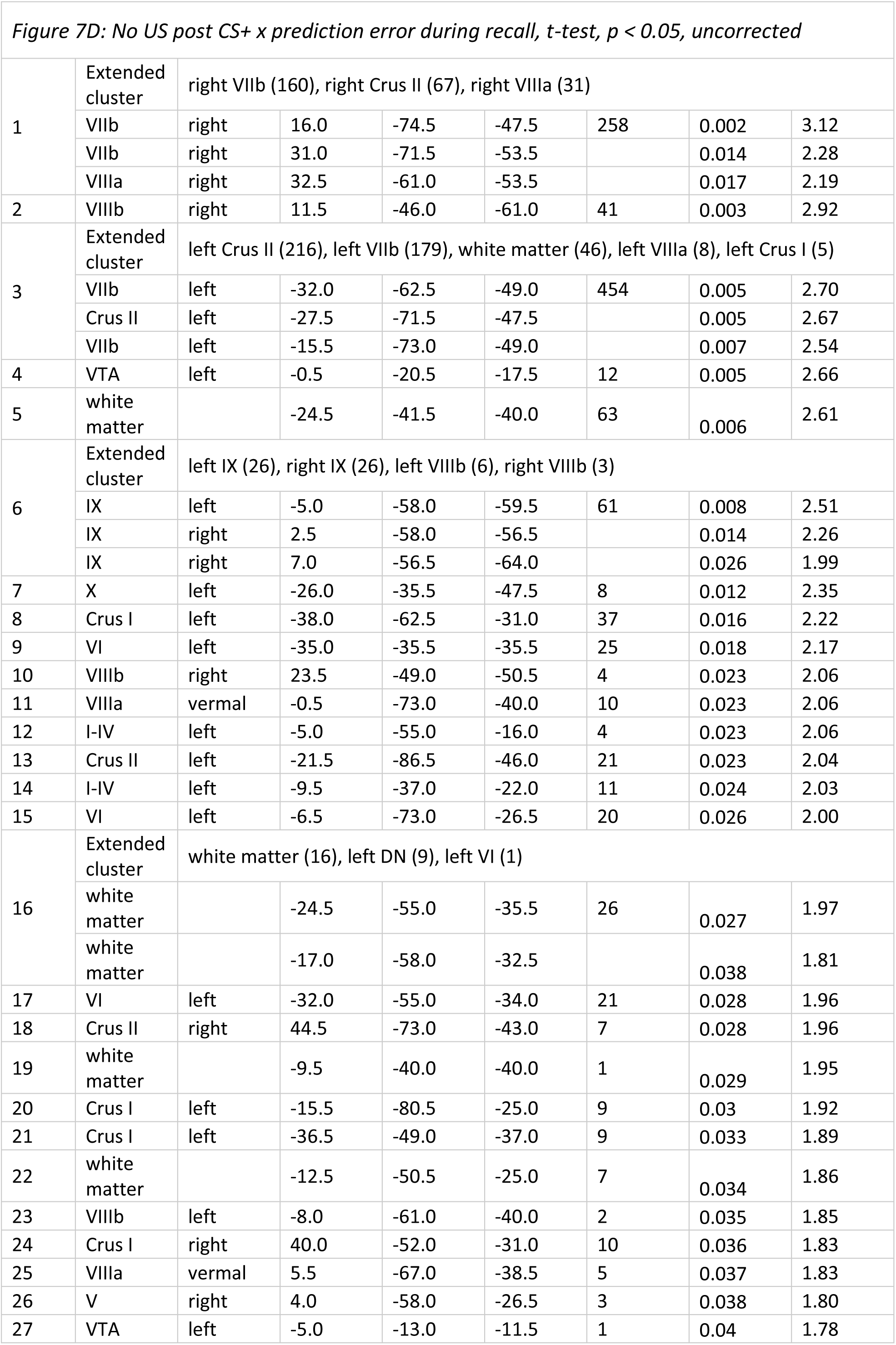
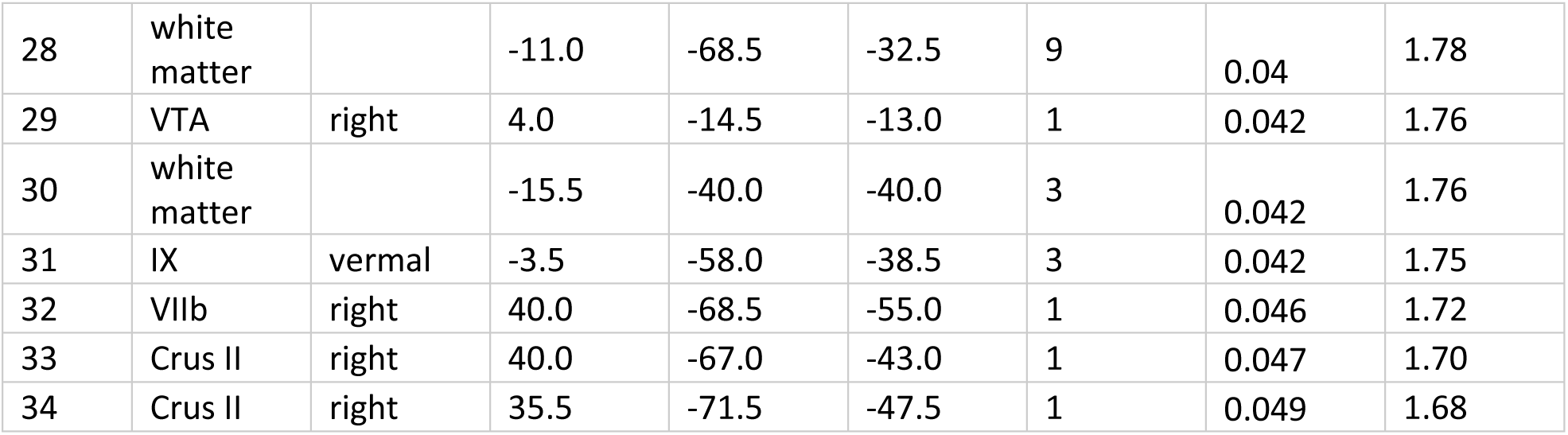
fMRI activation clusters (p < 0.05, uncorrected) related to the unexpected omission of the unconditioned stimulus (US) during extinction training (Figure 6 and 7). Clusters were identified in the cerebellar cortex, deep cerebellar nuclei (DCN), and ventral tegmental area (VTA). Up to three local maxima per cluster are reported, separated by at least 8 mm. Coordinates are given in MNI space (x, y, z). Cluster size is reported as number of voxels (voxel volume = 3.375 mm³). US: unconditioned stimulus; CS: conditioned stimulus; VTA: ventral tegmental area; DCN: deep cerebellar nuclei; DN: dentate nucleus; IN: interposed nucleus; FN: fastigial nucleus; MNI: Montreal Neurological Institute standard brain; t: t-statistic; punc: uncorrected p-value.

#### fMRI activations related to PPI connectivity during unexpected US omissions with a VTA seed. Uncorrected

**Table S12:**
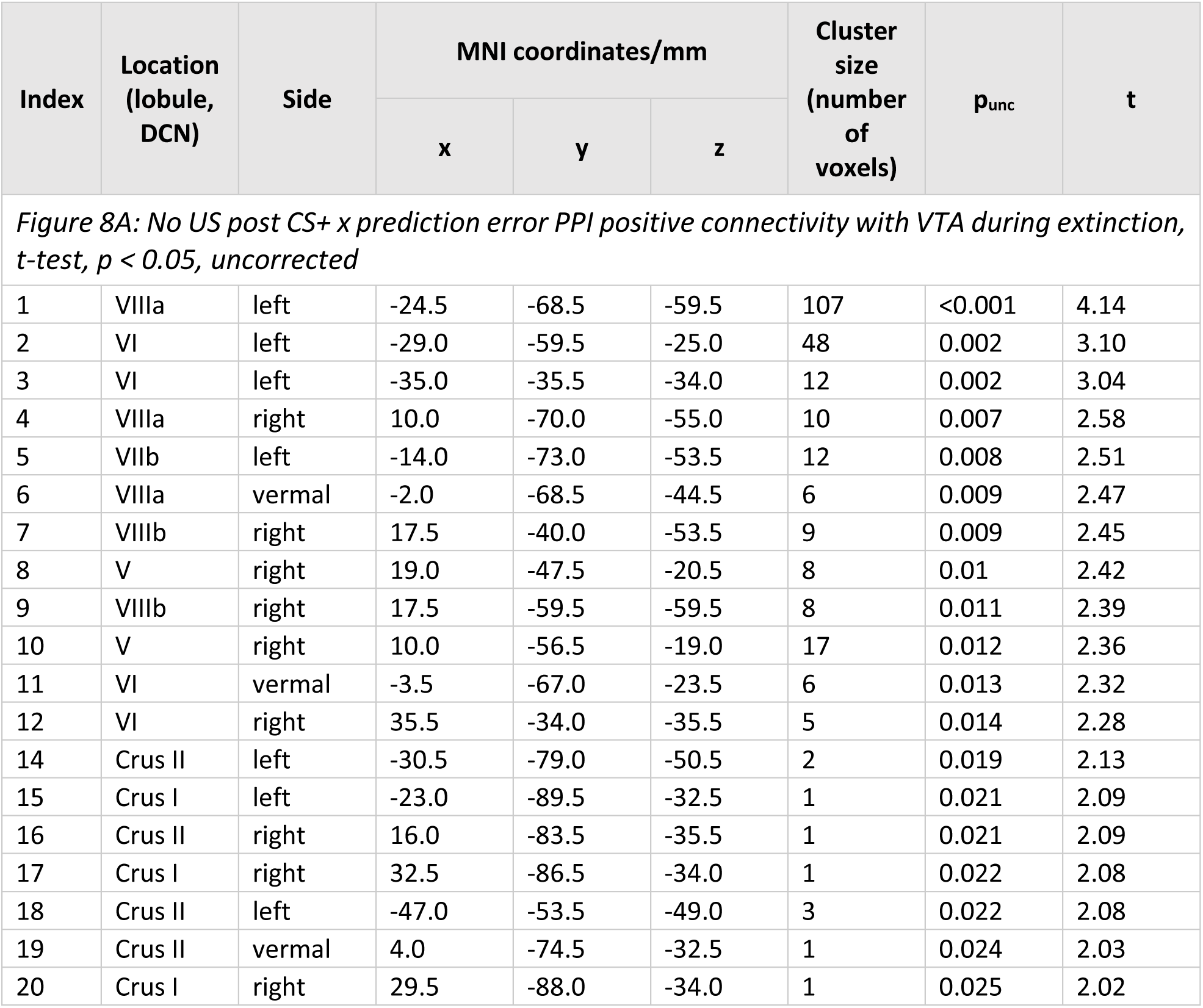
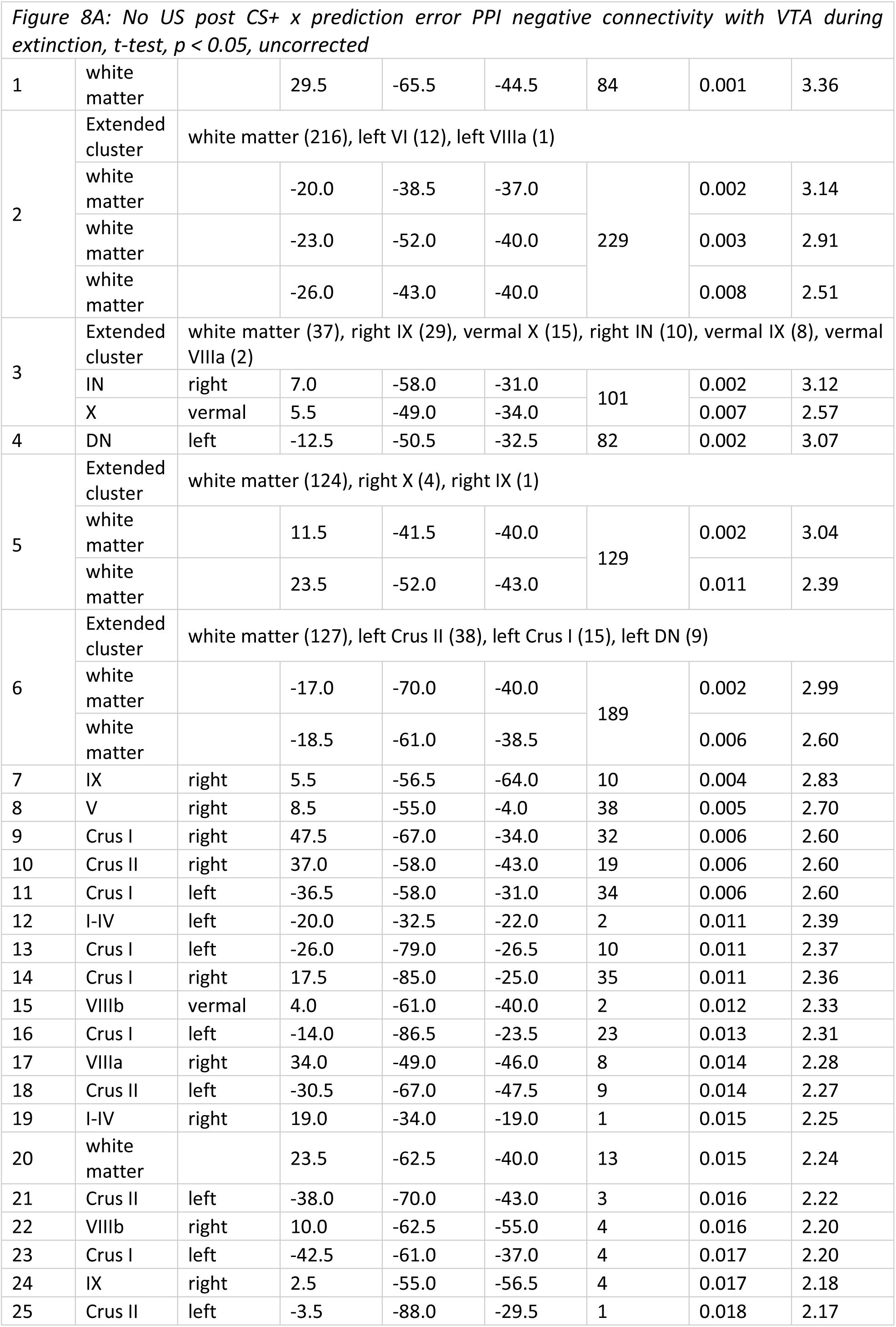
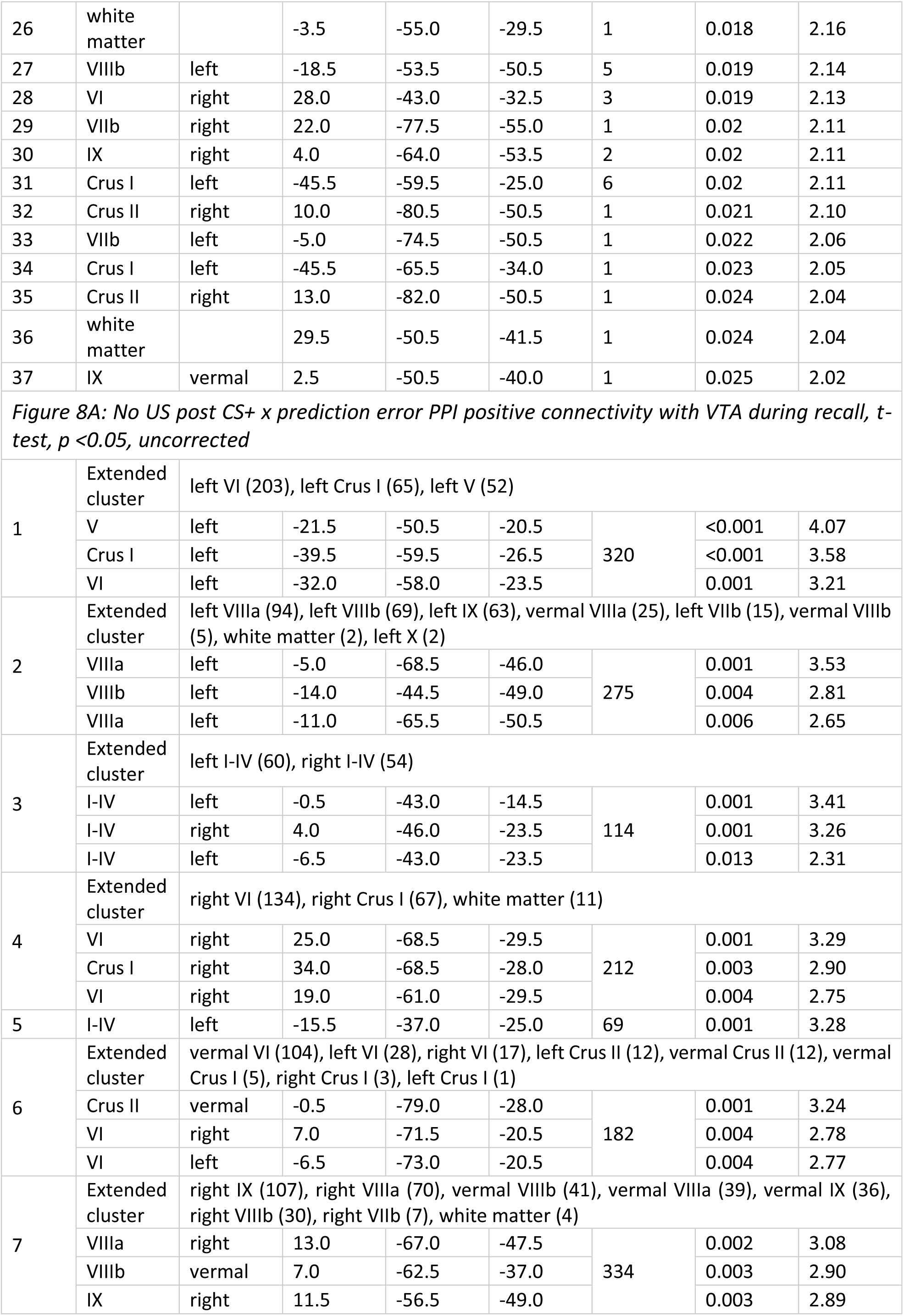
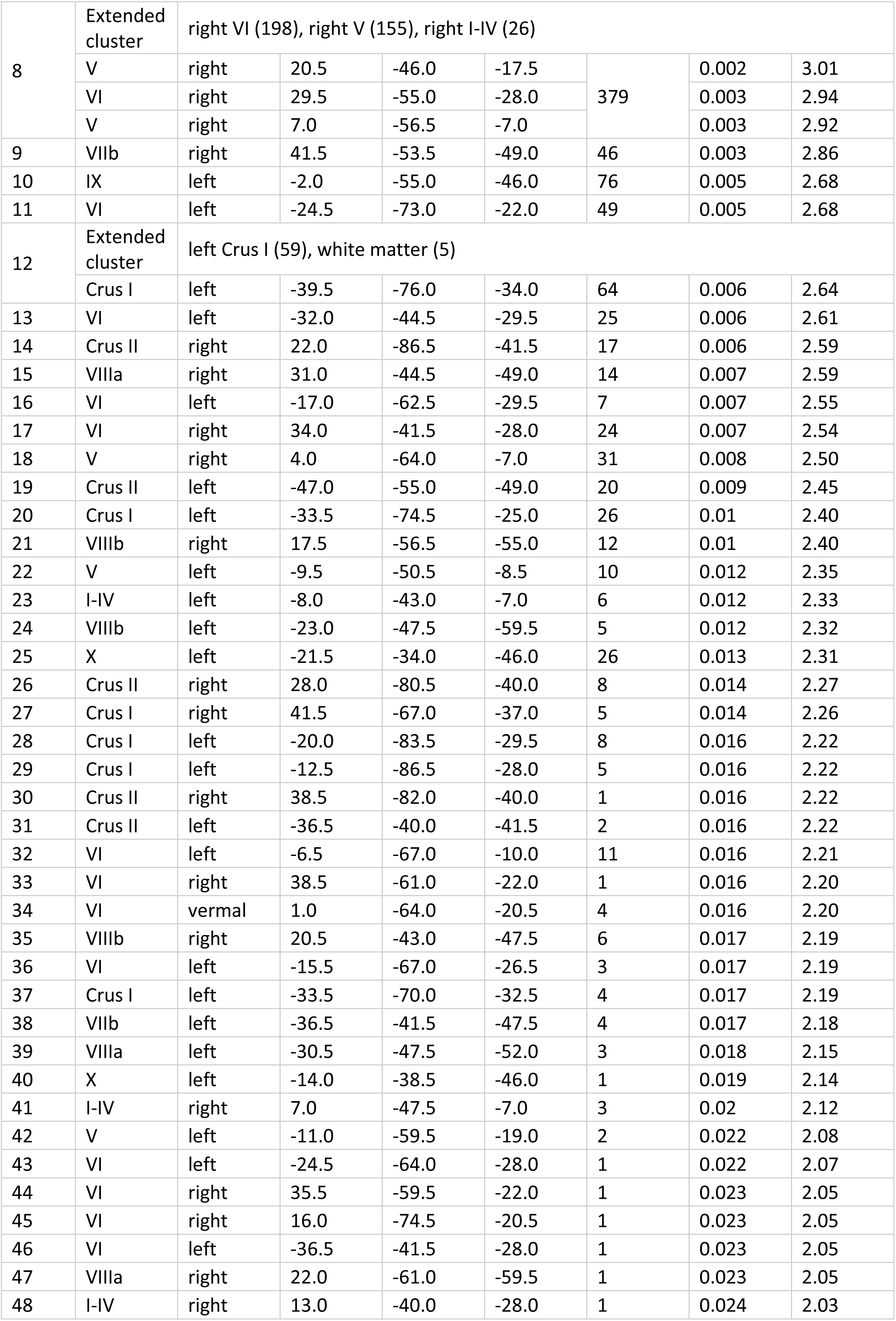
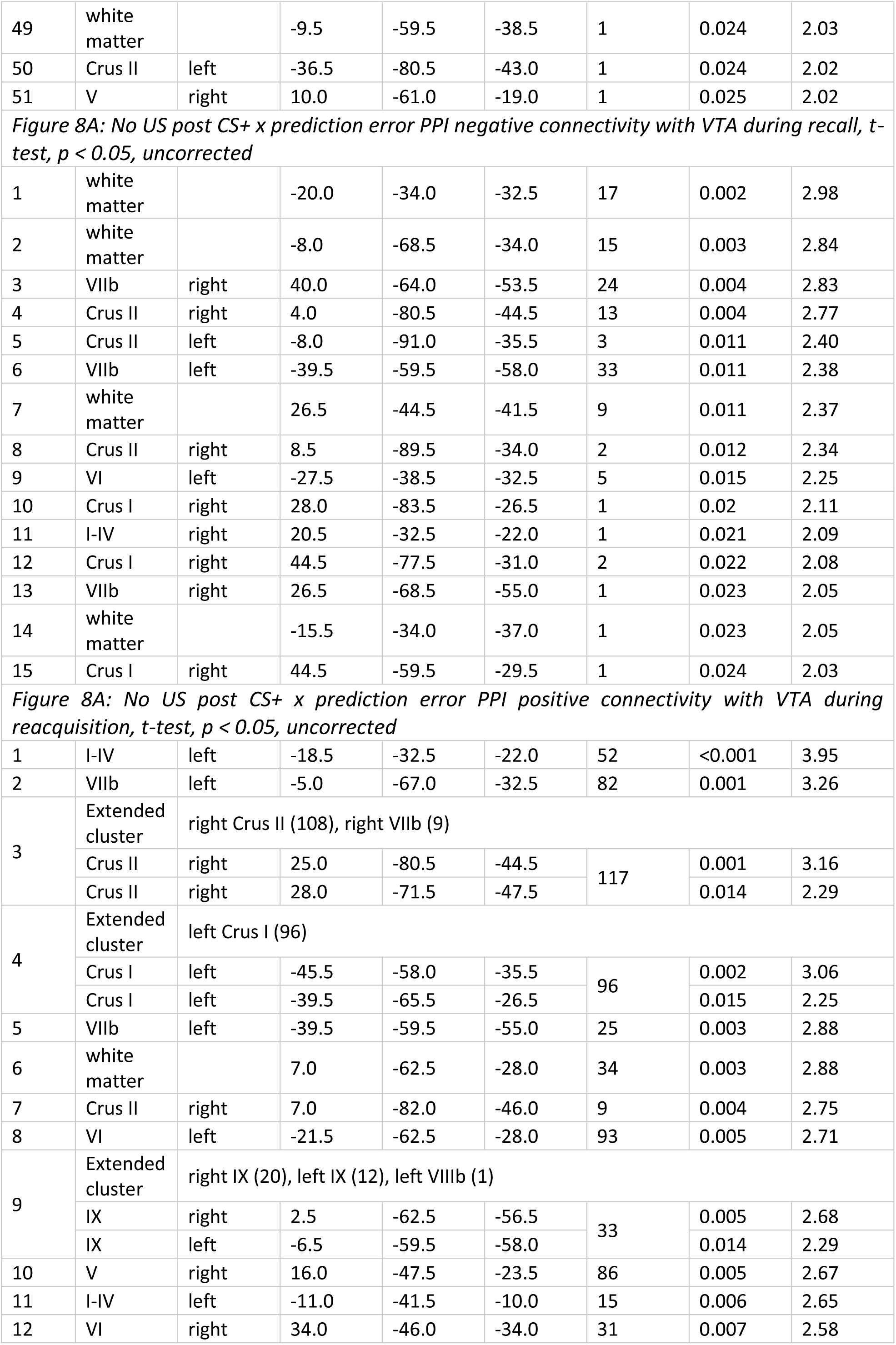
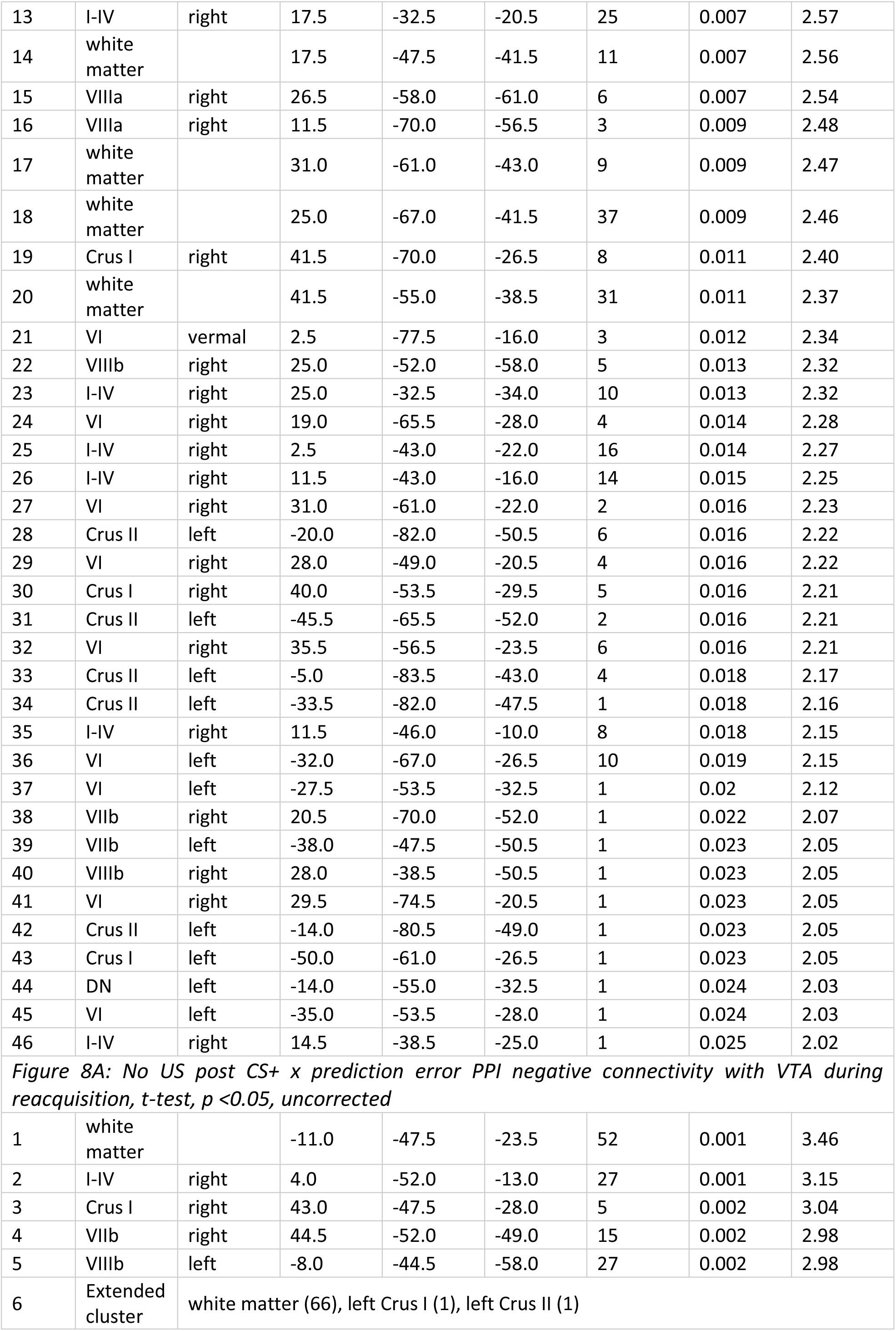
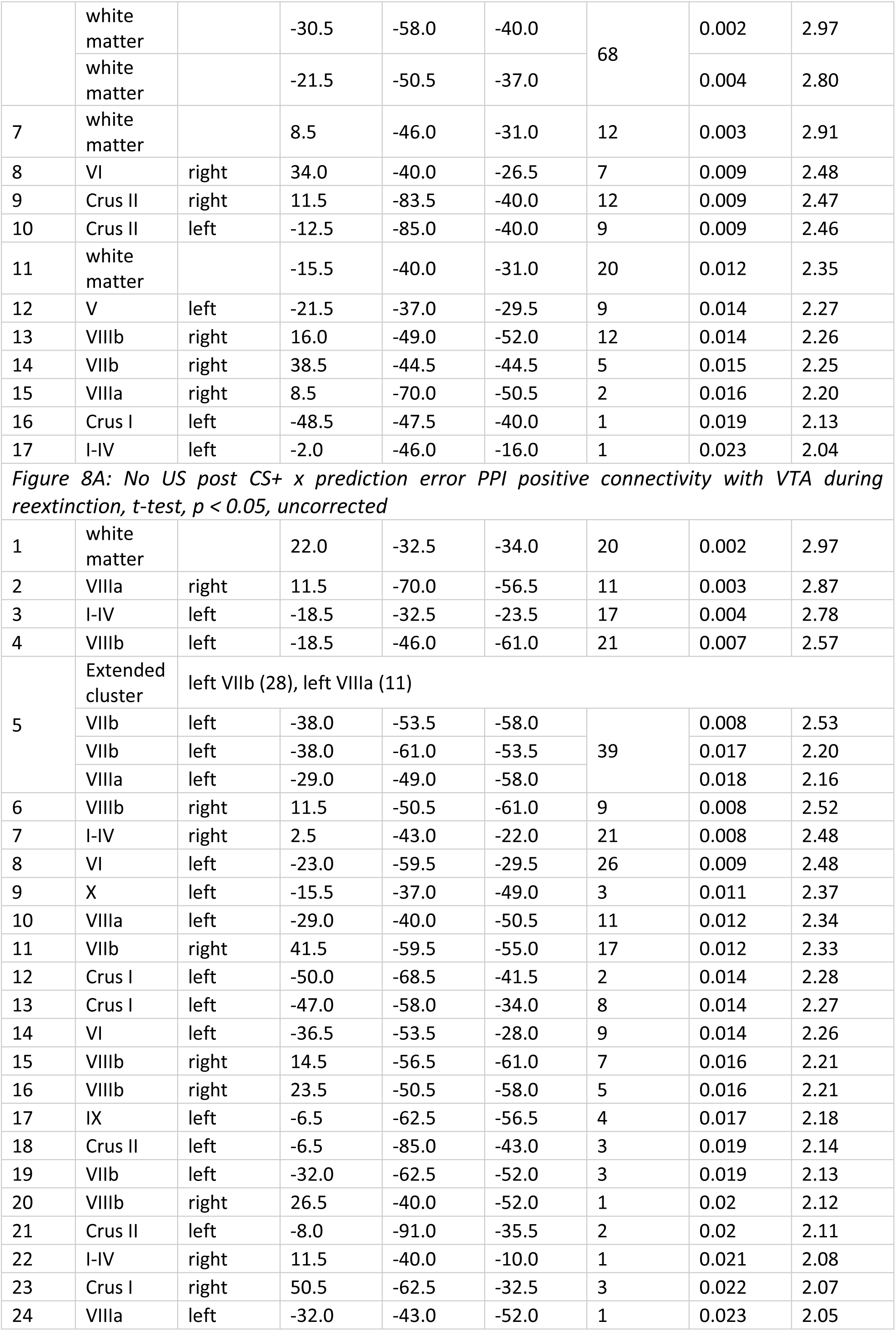
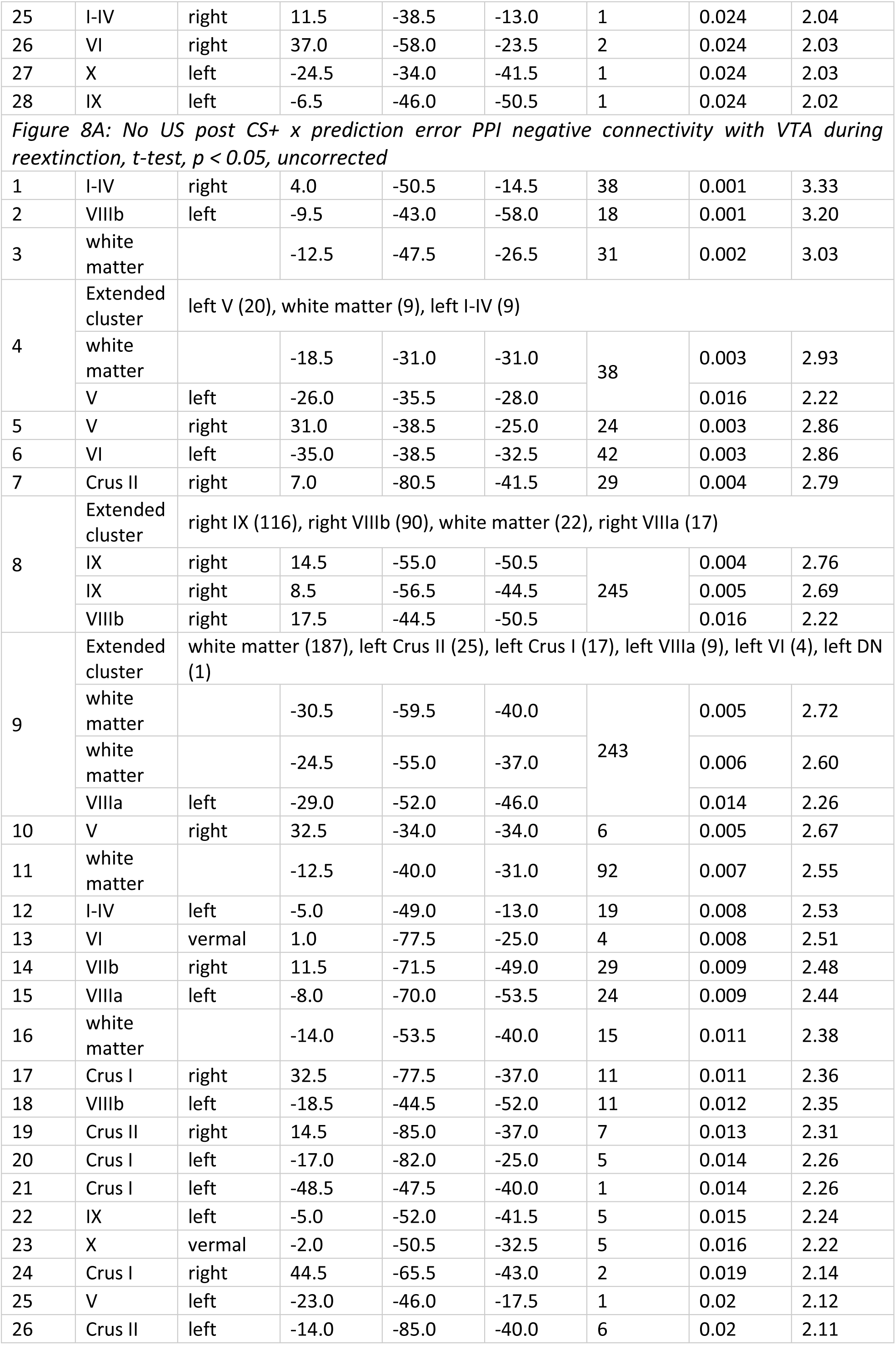
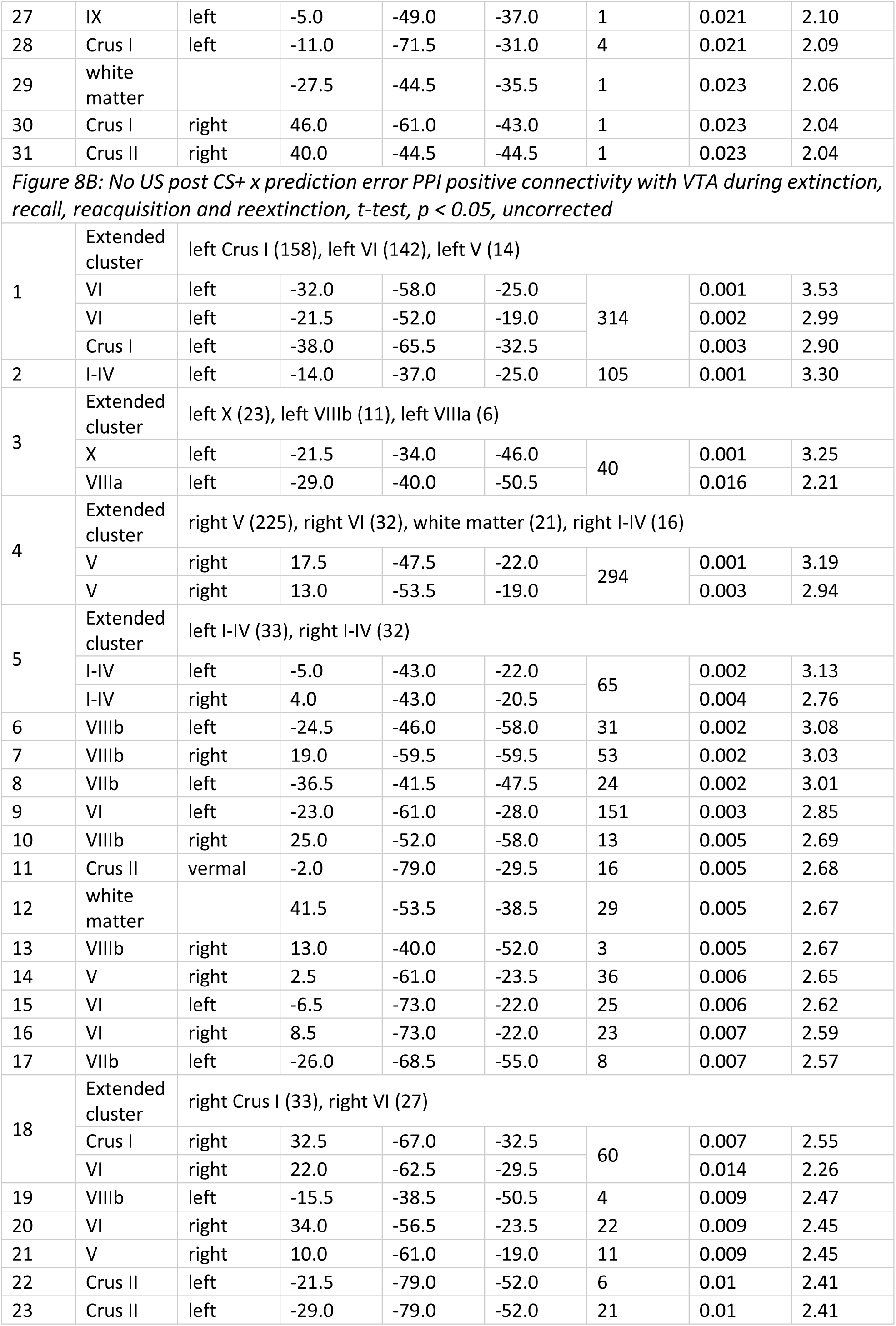
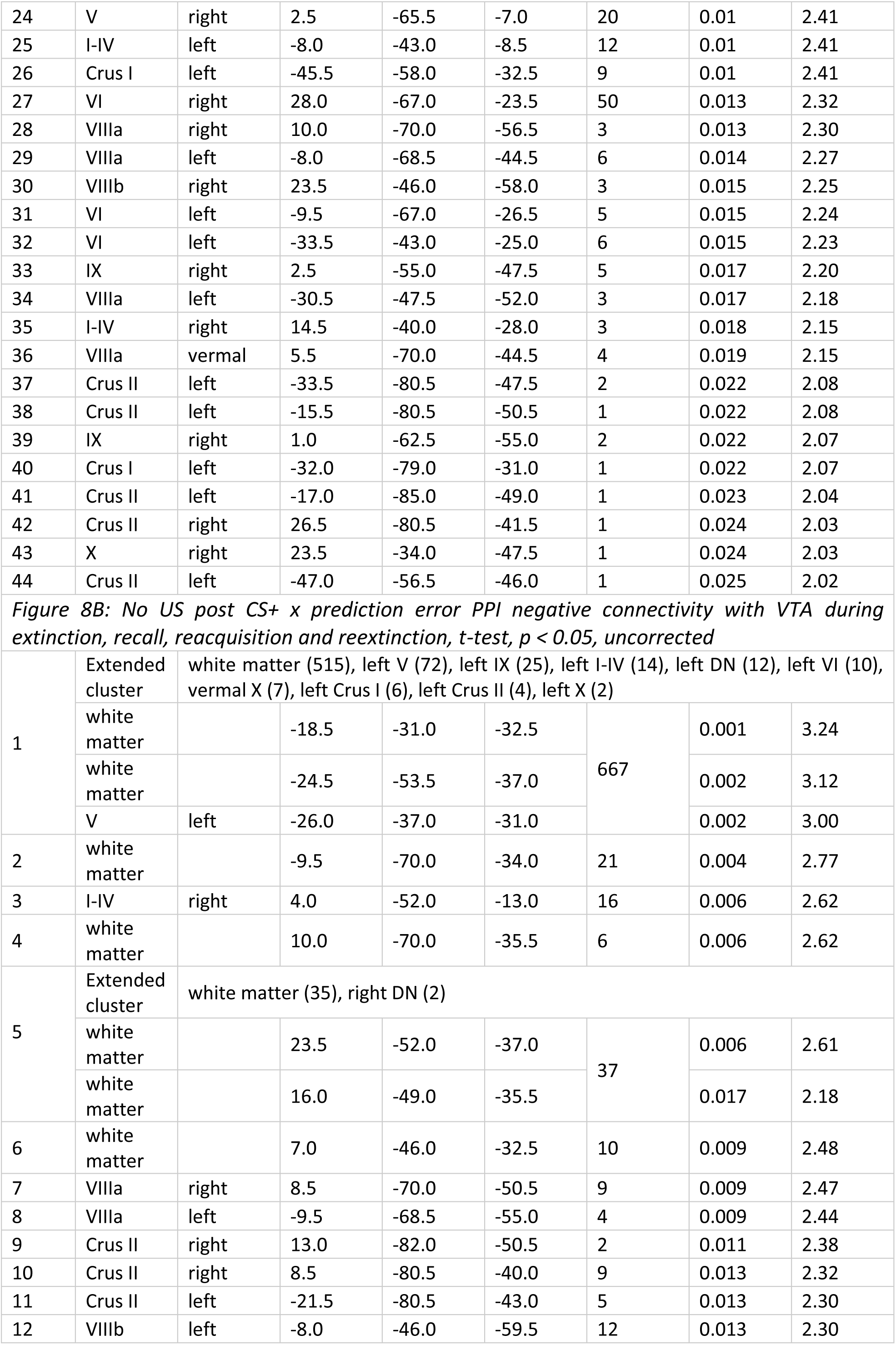
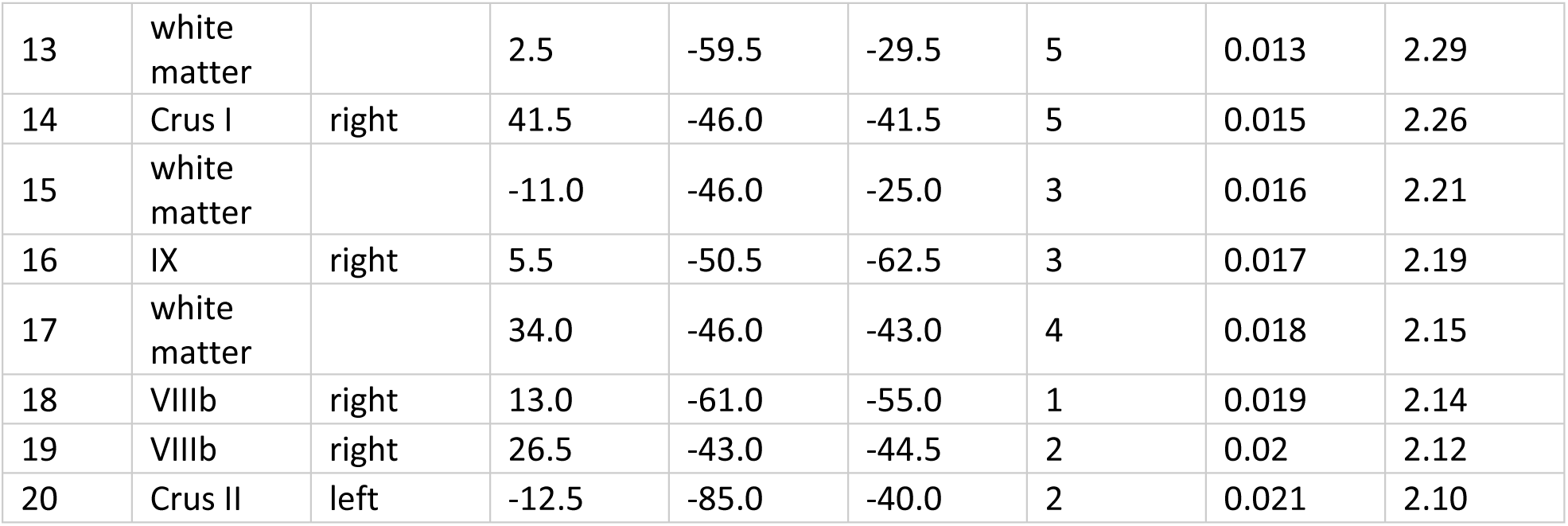
fMRI psychophysiological interaction (PPI) activation clusters (p < 0.05, uncorrected) related to connectivity with the ventral tegmental area (VTA) during unexpected omission of the unconditioned stimulus (US) using a VTA seed (Figure 8). Clusters were identified in the cerebellar cortex and deep cerebellar nuclei (DCN). Up to three local maxima per cluster are reported, separated by at least 8 mm. Coordinates are given in MNI space (x, y, z). Cluster size is reported as number of voxels (voxel volume = 3.375 mm³). US: unconditioned stimulus; CS: conditioned stimulus; VTA: ventral tegmental area; DCN: deep cerebellar nuclei; DN: dentate nucleus; IN: interposed nucleus; FN: fastigial nucleus; MNI: Montreal Neurological Institute standard brain; t: t-statistic; p_unc_: uncorrected p-value.

